# Genetically distinct clinical subsets, and associations with asthma and eosinophil abundance, within Eosinophilic Granulomatosis with Polyangiitis

**DOI:** 10.1101/491837

**Authors:** Paul A Lyons, James E Peters, Federico Alberici, James Liley, Richard M.R. Coulson, William Astle, Chiara Baldini, Francesco Bonatti, Maria C Cid, Heather Elding, Giacomo Emmi, Jörg Epplen, Loic Guillevin, David R. W. Jayne, Tao Jiang, Iva Gunnarsson, Peter Lamprecht, Stephen Leslie, Mark A. Little, Davide Martorana, Frank Moosig, Thomas Neumann, Sophie Ohlsson, Stefanie Quickert, Giuseppe A. Ramirez, Barbara Rewerska, Georg Schett, Renato A. Sinico, Wojciech Szczeklik, Vladimir Tesar, Damjan Vukcevic, The European Vasculitis Genetics Consortium, Benjamin Terrier, Richard A Watts, Augusto Vaglio, Julia U Holle, Chris Wallace, Kenneth G. C. Smith

## Abstract

Eosinophilic granulomatosis with polyangiitis (EGPA: formerly Churg-Strauss syndrome) is a rare inflammatory disease of unknown cause. 30% of patients have anti-neutrophil cytoplasm antibodies (ANCA) specific for myeloperoxidase (MPO). We performed a genome-wide association study (GWAS) of EGPA, testing 7.5 million genetic variants in 684 cases and 6,838 controls. Case-control analyses were performed for EGPA as a whole, and stratified by ANCA. To increase power, we used a conditional false discovery rate method to leverage findings from GWASs of related phenotypes. In total, 11 variants were associated with EGPA, two specifically with ANCA-negative EGPA, and one (*HLA-DQ*) with MPO+ANCA EGPA. Many variants were associated with asthma, eosinophilic and immune-mediated diseases and, strikingly, nine were associated with eosinophil count in the general population. Through Mendelian randomisation, we show that a primary tendency to eosinophilia underlies EGPA susceptibility. We demonstrate that EGPA comprises two genetically and clinically distinct syndromes, with ANCA-negative EGPA genetically more similar to asthma. MPO+ ANCA EGPA is an eosinophilic autoimmune disease sharing certain clinical features and an MHC association with MPO+ ANCA-associated vasculitis, while ANCA-negative EGPA may instead have a mucosal/barrier dysfunction origin. Five identified candidate genes are targets of therapies in development, supporting their exploration in EGPA.

## Introduction

Eosinophilic granulomatosis with polyangiitis (EGPA), once named Churg-Strauss syndrome, has a unique combination of clinical features that have some overlap with the other anti-neutrophil cytoplasmic antibody (ANCA)-associated vasculitis (AAV) syndromes, granulomatosis with polyangiitis (GPA) and microscopic polyangiitis (MPA)(1, 2). The initial report of Churg-Strauss syndrome described necrotizing vasculitis, eosinophilic tissue infiltration and extravascular granulomata at post-mortem(1). After a prodromal period characterized by asthma and eosinophilia that may last for some years, patients develop the more distinctive clinical features of EGPA. These include various combinations of neuropathy, pulmonary infiltrates, myocarditis, and ear, nose and throat (ENT), skin, gastrointestinal and renal involvement(2).

Despite being classified as a form of ANCA-associated vasculitis(3), vasculitis is not always evident (discussed in more detail **Supplementary Appendix**) and only 30-40% are ANCA positive (almost all against myeloperoxidase (MPO) rather than proteinase-3 (PR3)). These observations, together with increasing evidence that clinical distinctions can be drawn between the ANCA positive and negative subsets of EGPA(4–6), suggest that clinically important subsets may exist within it(2). The etiology of EGPA is unknown; it is too uncommon for familial clustering to have been quantified or major genetic studies performed. Candidate gene studies in small cohorts have reported associations of EGPA with *HLA-DRB4* and *DRB1*07* and protection by *DRB3* and *DRB1*13*(7, 8), suggesting a genetic contribution of uncertain size.

Here we describe the first genome-wide association study (GWAS) of EGPA. It demonstrates that EGPA is polygenic with genetic distinctions between MPO+ and ANCA-negative disease, correlating with different clinical features. The genetic associations themselves point to dysregulation of pathways controlling eosinophil biology, severe asthma and vasculitis, beginning to explain the development and clinical features of disease. These results suggest that EGPA might be comprised of two distinct diseases defined by ANCA status, and provide a scientific rationale for targeted therapy.

## Results

### The genetic contribution to EGPA

The EGPA patients’ clinical features are summarised in **Table 1**. Further details of the cohorts are presented in **Supplementary Tables 1-3**. Consistent with previous reports(2), 171 (32%) were positive for ANCA, and of the 166 who were positive for specific ANCA by ELISA, 161 (97%) had MPO+ANCA and five PR3+ANCA. We performed genome-wide association testing at 7.5 million genetic variants in 542 cases and 6717 controls. Genotyping was performed using the Affymetrix Axiom UK Biobank array and high-density genotype data was generated through imputation against the 1000 Genomes phase 3 reference panel (**Methods**). Despite attempting to control for population stratification by the inclusion of 20 genetic principal components (PCs) as covariates in the logistic regression model, the genomic inflation factor lambda remained 1.10. To account for residual genomic inflation, we calculated an adjusted genome-wide significance p-value threshold of 1×10^−8^ (**Methods**). Use of this adjusted threshold is equivalent to using a p-value threshold of 5×10^−8^ if there were no elevation of the genomic inflation factor (assuming that the level of inflation is uniform across the genome). Three genetic associations met this adjusted threshold (**Figure 1A**, **Supplementary Figure 1**, **Table 2**). The strongest association was with the MHC, and the others were on chromosome 2 near *BCL2L11* (encoding Bim), and on chromosome 5 near the *TSLP* gene (which encodes Thymic stromal lymphopoietin; TSLP). Since adjustment for genetic PCs did not completely abrogate genomic inflation, we used a linear mixed model as an alternative method to control for population stratification (**Methods**). This method more effectively controlled genomic inflation (lambda 1.047) and the 3 associations were preserved at genome-wide significance (**Supplementary Table 4**), providing strong evidence against these signals being driven by population stratification.

**Figure 1.**
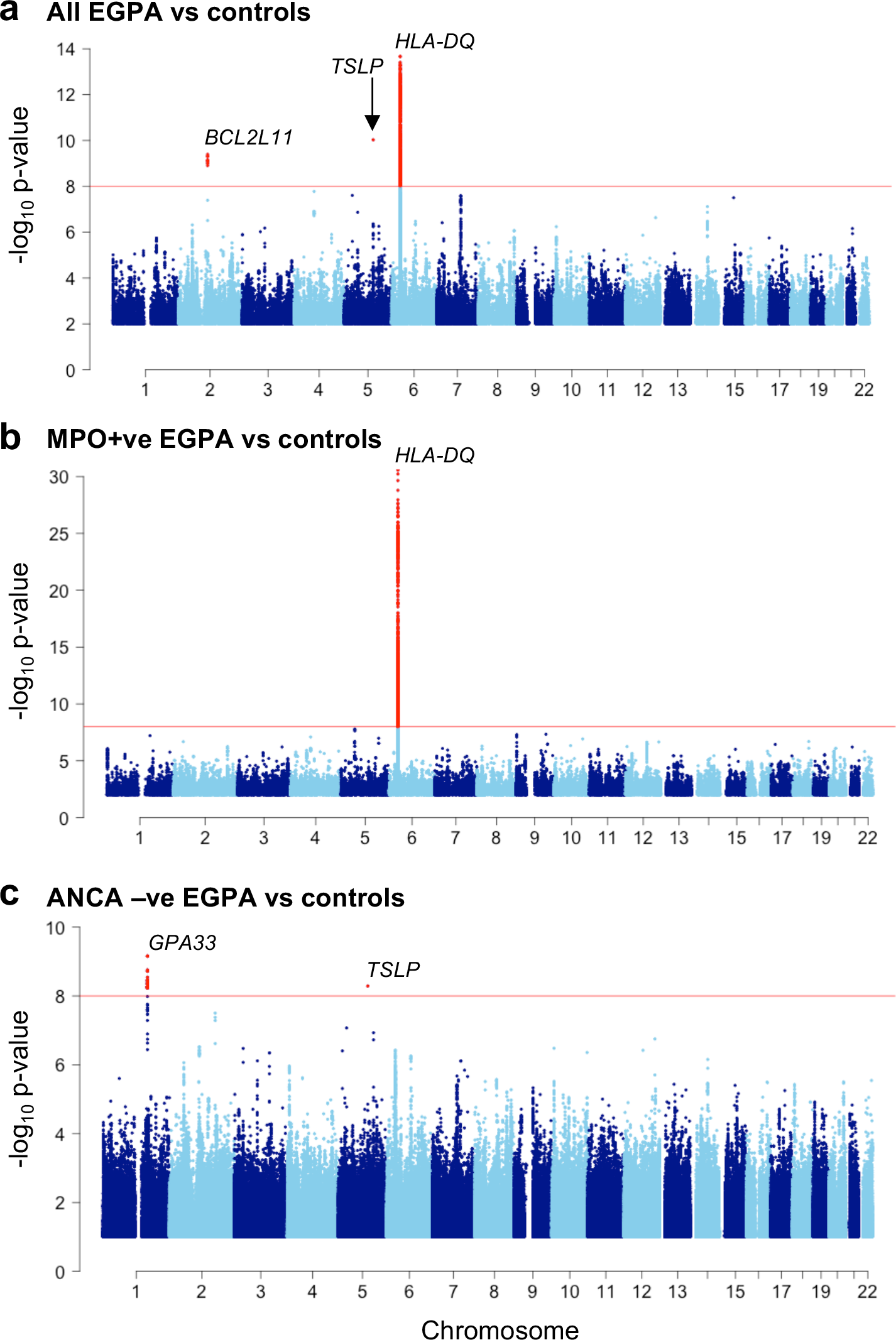
Manhattan plot of genetic associations with EGPA. Manhattan plots showing the association between SNPs and (**A**) all EGPA cases, (**B**) the subset of cases with MPO+ANCA EGPA (n=161), and **(C)** ANCA-negative EGPA cases (n=358). SNPs at loci reaching genome-wide significance are highlighted in red. The red horizontal lines indicate the threshold for declaring genome-wide significance (p = 1×10^−8^).

**Table 1.** Comparison of clinical features between MPO positive and ANCA negative EGPA patients.

|  | **All patients**<br>n=542<br>(%) | **ANCA –ve**<br>n=358<br>(%) | **MPO +ve**<br>n=161<br>(%) | **MPO +ve**<br>**versus ANCA**<br>**–ve**<br>**P-value** | **Bonferroni-**<br>**corrected**<br>**P-value** |
| --- | --- | --- | --- | --- | --- |
| Eosinophilia | 542 (100) |  |  |  |  |
| Asthma | 542 (100) |  |  |  |  |
| Neuropathy | 344 (63.5) | 204 (57.0) | 127 (78.9) | $2.6 \times 10^{-6}$ | **$2.1 \times 10^{-5}$** |
| Lung infiltrates | 306 (56.5) | 220 (61.5) | 73 (45.3) | 0.00087 | **0.0070** |
| ENT | 466 (86) | 315 (88.0) | 130 (80.7) | 0.041 | 0.32 |
| Cardiomyopathy | 136 (25.1) | 108 (30.2) | 23 (14.3) | 0.00018 | **0.0015** |
| Glomerulonephritis | 83 (15.3) | 33 (9.2) | 46 (28.6) | $2.9 \times 10^{-8}$ | **$2.3 \times 10^{-7}$** |
| Lung haemorrhage | 22 (4.1) | 14 (3.9) | 7 (4.3) | 1.0 | 1.0 |
| Purpura | 138 (25.5) | 92 (25.7) | 37 (23.0) | 0.58 | 1.0 |
| Positive biopsy* | 215 (41.2†) | 147 (42.7†) | 61 (38.6†) | 0.43 | 1.0 |

ENT= ear, nose and throat. P-values were calculated from 1 degree of freedom chi-squared tests with Yates' continuity correction. Bonferroni correction was undertaken to account for the 8 clinical features tested.
\* Defined as a biopsy showing histopathological evidence of eosinophilic vasculitis, or perivascular eosinophilic infiltration, or eosinophil-rich granulomatous inflammation or extravascular eosinophils in a biopsy including an artery, arteriole, or venule.
† Percentages are of those with available data. Biopsy data were unavailable for 21 (3.9%) of patients. Biopsy data was available for 158/161 (98.1%) MPO positive patients and 344/358 (96.1%) ANCA negative patients.

**Table 2.** Genetic Associations with EGPA.

| Chr | SNP | Gene | Cont<br>MAF | Total EGPA<br>N=542 |  |  | MPO+ve EGPA<br>N =161 |  | ANCA-ve EGPA<br>N=358 |  | Asthma<br>N= 10,365 |  | Eosinophil count<br>N=175,000 |  |
| --- | --- | --- | --- | --- | --- | --- | --- | --- | --- | --- | --- | --- | --- | --- |
|  |  |  |  | Case<br>MAF | OR | P | OR | P | OR | P | P | cFDR <sup>^</sup><br>EGPA\|asthma | P | cFDR <sup>‡</sup><br>EGPA\|Eos |
| 2 | rs72836352 | *BCL2L11* | 0.10 | 0.18 | 1.93 | 4.0x10<sup>-10</sup> | 2.18 | 3.7x10<sup>-5</sup> | 1.92 | 3.0x10<sup>-7</sup> |  |  | 9.3x10<sup>-26</sup> |  |
| 5 | rs1837253 | *TSLP* | 0.26 | 0.17 | 1.66 | 9.3x10<sup>-11</sup> | 1.57 | 9.4x10<sup>-4</sup> | 1.72 | 5.1x10<sup>-9</sup> | 2.3x10<sup>-5</sup> | 1.7x10<sup>-8</sup> | 1.9x10<sup>-22</sup> | 2.2x10<sup>-8</sup> |
| 6 | rs28724235 | *HLA-DQ* | 0.06 | 0.11 | 2.67 | 1.6x10<sup>-11</sup> | 20.7 | 2.4x10<sup>-31</sup> | 1.19 | 0.35 |  |  | 4.3x10<sup>-8</sup> |  |
| 1 | rs72689399 | *GPA33* | 0.01 | 0.03 | 3.83 | 1.8x10<sup>-6</sup> | 0.89 | 0.84 | 7.71 | 6.8x10<sup>-10</sup> |  |  | na |  |
| 5 | rs11745587<sup>†</sup> | *IRF1/IL5* | 0.35 | 0.40 | 1.42 | 1.1x10<sup>-6</sup> | 1.18 | 0.20 | 1.57 | 1.17x10<sup>-7</sup> | 1.8x10<sup>-3</sup> | 3.0x10<sup>-4</sup> | 1.6x10<sup>-29</sup> | 1.6x10<sup>-4</sup> |
| 6 | rs6454802 | *BACH2* | 0.40 | 0.31 | 0.71 | 4.7x10<sup>-7</sup> | 0.79 | 0.05 | 0.67 | 8.0x10<sup>-7</sup> | 2.2x10<sup>-3</sup> | 1.6x10<sup>-4</sup> | 1.0x10<sup>-18</sup> | 4.8x10<sup>-5</sup> |
| 10 | rs1623646<sup>†</sup> | - | 0.27 | 0.20 | 1.40 | 1.5x10<sup>-5</sup> | 1.23 | 0.13 | 1.54 | 4.0x10<sup>-6</sup> | 2.0x10<sup>-3</sup> | 1.4x10<sup>-3</sup> | 6.5x10<sup>-13</sup> | 7.1x10<sup>-4</sup> |
| 7 | rs42041<sup>\$</sup> | *CDK6* | 0.24 | 0.31 | 1.52 | 6.4x10<sup>-7</sup> | 1.47 | 0.005 | 1.45 | 8.3x10<sup>-5</sup> | | | 4.6x10<sup>-8</sup> | 5.1x10<sup>-4</sup> |
| 16 | rs4781047 | *SOCS1* | 0.43 | 0.46 | 0.74 | 2.0x10<sup>-5</sup> | 0.70 | 0.004 | 0.76 | 8.1x10<sup>-4</sup> |  |  | 7.8x10<sup>-14</sup> | 9.2x10<sup>-4</sup> |
| 3 | rs9290877 | *LPP* | 0.30 | 0.38 | 1.36 | 3.0x10<sup>-5</sup> | 1.51 | 0.001 | 1.29 | 0.004 |  |  | 9.1x10<sup>-14</sup> | 1.3x10<sup>-3</sup> |
| 12 | rs187564398 | *TBX3* | 0.02 | 0.04 | 3.72 | 2.3x10<sup>-7</sup> | 3.10 | 0.03 | 4.78 | 1.8x10<sup>-7</sup> |  |  | 6.9x10<sup>-3</sup> | 8.8x10<sup>-4</sup> |

<sup>^</sup> Reached GW-significance (adjusted for genomic inflation) in EGPA by cFDR. Significance threshold cFDR <1.1x10<sup>-3</sup>, FDR (excl MHC) <0.0053 (equivalent to P 1x10<sup>-8</sup>)
<sup>‡</sup> Reached GW-significance (adjusted for genomic inflation) in EGPA by cFDR. Significance threshold cFDR <2.4x10<sup>-3</sup>, FDR (excl MHC) <0.0053 (equivalent to P 1x10<sup>-8</sup>)
<sup>†</sup> SNP most associated with EGPA that was directly genotyped in asthma. Stronger EGPA associations exist at this locus (see Fig. 2)
<sup>\$</sup> SNP most associated with EGPA that was directly genotyped in eosinophil count. Strongest EGPA association (p = 2.6x10<sup>-8</sup>) at rs42046 (r<sup>2</sup> 0.8).
na data not available

While replication is the gold standard for GWAS studies, the rarity of EGPA (annual incidence 1-2 cases per million) makes recruitment of an adequately powered replication cohort challenging. Nonetheless, a second cohort of 142 EGPA patients from two European centres was identified; 43 (30%) had MPO+ANCA (**Supplementary Table 5**). Despite limited power, all three genetic loci were nominally significant in this replication cohort (**Supplementary Table 6**). Moreover, there was a strong correlation between the estimated effect sizes in the primary and replication cohorts (r = 0.78, p = 0.005), providing additional support for the reported associations (**Supplementary Figure 2**).

To quantify the genetic influence on EGPA, the total narrow-sense heritability (h^2^) was estimated: the genotyped variants additively explained approximately 22% of the total disease liability (EGPA h^2^≥0.22, standard error 0.082). Thus, while specific loci have substantial effect sizes in EGPA (**Table 2**), the contribution of genetics overall is similar to other immune-mediated diseases (e.g. h^2^ estimates for Crohn’s disease and ulcerative colitis are 26% and 19% respectively(9)).

### Conditional false discovery rate (cFDR) analysis reveals additional associations

One method to overcome limitations posed by GWAS sample size in rare diseases is to leverage results from GWAS of related phenotypes using the pleiotropy-informed Bayesian conditional false discovery rate (cFDR) method(10, 11). Asthma and eosinophil count were chosen as relevant traits, as they are both ubiquitous features of EGPA, and SNPs showing association with them showed a trend for greater association with EGPA (**Supplementary Figure 3**), with a consistent direction of effect (**Supplementary Table 7**), suggesting shared genetic architecture.

Use of the pleiotropy-informed Bayesian cFDR conditioning on asthma(12) revealed additional EGPA associations at 5q31.1, near *IRF1* and *IL5*, 6q15, in *BACH2* and an intergenic region (a ‘gene desert’) at 10p14. We then compared EGPA to data from a GWAS of eosinophil count in the general population(13), identifying a further 4 EGPA associations near *CDK6*, *SOCS1*, *LPP* and *TBX3* (**Table 2**). These 7 additional associations were preserved in the mixed model analysis. The identification of 11 associations with EGPA is consistent with it being a polygenic disease.

### Genetic and clinical distinctions between MPO+ANCA and ANCA-negative EGPA

EGPA has been classified alongside MPA and GPA as an AAV(3), despite ANCA being only found in the minority of cases. While asthma and eosinophilia are common to all patients with EGPA, other clinical features of the disease vary between ANCA positive and negative subgroups(2, 4–6, 14). We confirmed these phenotypic differences, with glomerulonephritis and neuropathy (clinical features consistent with vasculitis) more prevalent in the ANCA positive subgroup, whereas lung infiltrates and cardiac involvement were common in the ANCA negative subgroup (**Table 1**). These differential clinical associations remained statistically significant after adjustment for country of origin, suggesting a true biological difference between the subgroups (**Methods**, **Supplementary Tables 8-9**, **Supplementary Note**). These data, together with the analogous situation where patients with MPO+ANCA or PR3+ANCA GPA/MPA have distinct genetic associations(15, 16), suggested there may be genetic differences between MPO+ANCA and ANCA-negative EGPA.

We therefore compared these subsets to healthy controls, excluding the PR3+ANCA patients as there were too few to form a useful subset (**Figure 1B-C**, **Table 2**). The *HLA-DQ* association was only seen in the MPO-ANCA group, despite its smaller size (**Table 2**). In contrast, variants near *GPA33* and at *IRF1/IL5* reached genome-wide significance in the ANCA-negative subset (the latter when conditioning on asthma) but were not associated with the MPO+ANCA group. Three variants, at *BCL2L11*, *TSLP* and *CDK6*, had comparable odds ratios in each subgroup, suggesting their association with EGPA was independent of ANCA. Variants at *BACH2*, Chromosome 10, *SOCS1*, *LPP* and *TBX3* had similar odds ratios in both subsets, though genome-wide significant associations were only identified in the combined cohort when conditioning on asthma or eosinophil count, most likely due to limited power (**Table 2**).

A signal that appears specific to one subgroup might reflect lack of power to detect it in the other rather than necessarily indicating true genetic heterogeneity between the two subgroups. To formally address this issue, we compared genotypes of MPO+ and ANCA- samples directly with one another (i.e. a ‘within-cases’ analysis, independent of controls), and computed p-values for the 10 SNPs declared hits in our primary analysis agnostic of ANCA status, (ie either genome-wide significant in all EGPA cases vs controls or by cFDR). A significant difference in allelic frequency between the two subtypes at a given SNP indicates a subtype dependent effect on disease susceptibility. This analysis of MPO+ versus ANCA -ve cases revealed a genome-wide significant association at rs28724235 in the *HLA* (P 2.2×10^−11^). The variant in the *C5orf56-IRF1-IL5* region (rs11745587) showed a nominally significant association (P 6.8×10^−3^). There was no association identified at the other loci tested, either because of lack of power or because the signals at these loci were not subgroup-specific. In summary, this analysis provides robust evidence of a differential genetic basis of MPO+ and ANCA- EGPA at the *HLA* region and are suggestive of a subtype specific effect at *C5orf56-IRF1-IL5*.

In the small replication cohort, consistent with the primary cohort, the association at the *HLA-DQ* locus was stronger with the MPO-ANCA group than with EGPA overall (**Supplementary Table 10**). No evidence of association was observed between variants at *GPA33* and the ANCA-negative subgroup although, due to the low minor allele frequency and patient number, power to detect an effect was only 14%.

In light of the clinical and genetic differences identified between MPO positive and ANCA negative EGPA, we tested whether one subset was more genetically similar to asthma than was the other (**Methods**, **Supplementary Note**). With the *HLA* region removed, ANCA-negative EGPA was more genetically similar to asthma than was MPO+ EGPA (**Supplementary Note, Supplementary Figures 4-5**). This provides evidence for genetic differences between the two subtypes outside the *HLA* region, and suggests that the aetiology of ANCA-negative EGPA may more closely resemble that of asthma than does MPO+ EGPA. Differences are also seen in the genetic relatedness to IBD (see below) but we have insufficient power to formally demonstrate genetic variance between ANCA-negative and ANCA +ve EGPA.

Thus EGPA has a complex inheritance. Some loci are associated with both MPO+ANCA and ANCA-negative EGPA, consistent with the phenotypic overlap between the subsets and with their shared prodrome. Others are associated with only one subgroup. That these genetically distinct subsets of EGPA align with clinical disease phenotype (**Table 1**) suggests differences in pathogenesis and that the current clinical classification system should be re-visited (**Figure 3**).

**Figure 2.**
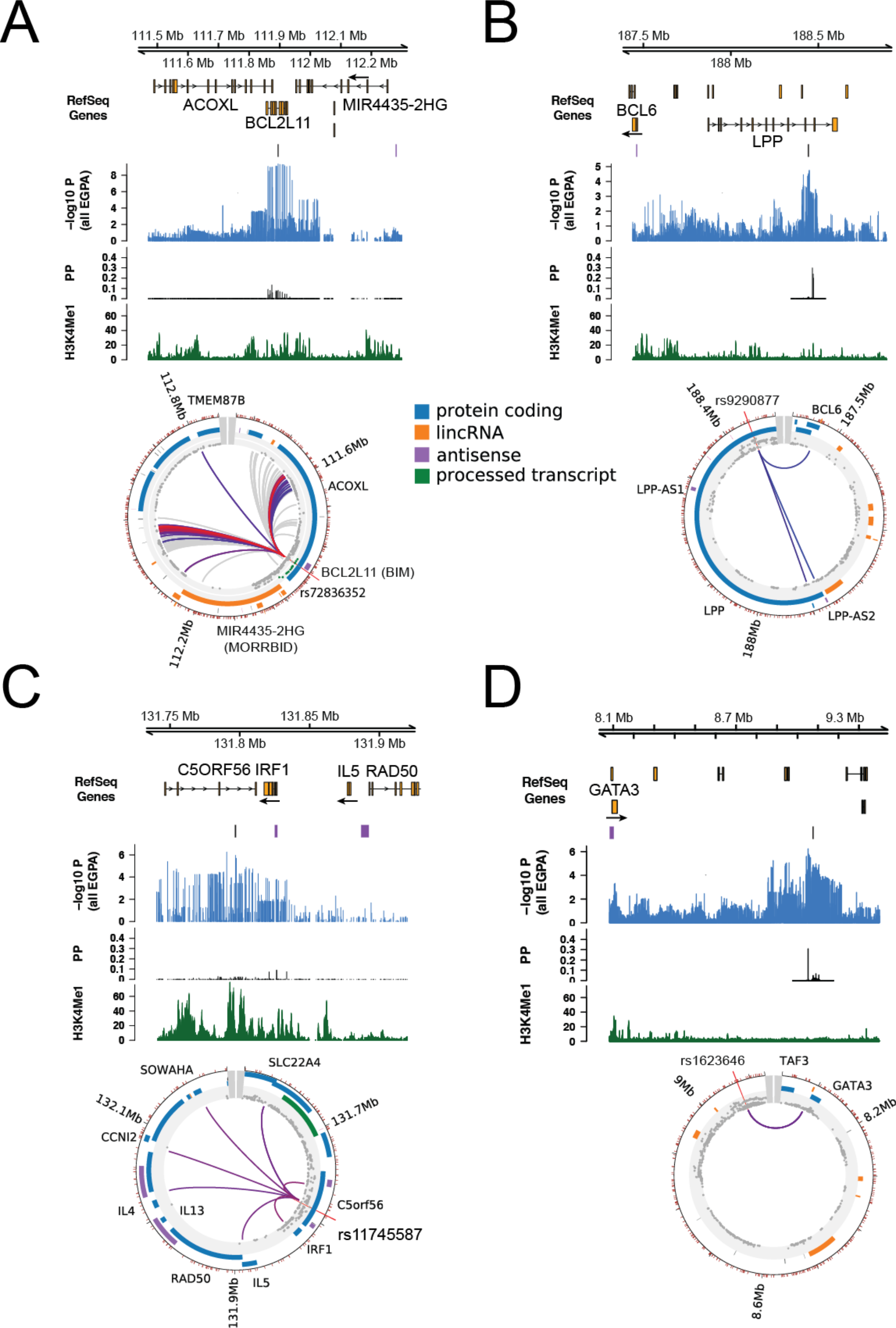
Genomic features at four EGPA-associated loci. Genomic positions from the hg19 genome build, representative RefSeq genes, SNP associations with EGPA, causal variant mapping (expressed as posterior probabilities, PP), H3K4 mono-methylation data and long-range DNA interactions are shown for **(A)** the *BCL2L11* region, **(B)** the *LPP* region, **(C)** the C5orf56-IRF1-IL5 region and **(D)** the 10p14 intergenic region. Arrows indicate direction of transcription. For further details regarding the promoter enhancer interaction mapping, including cell types analyzed at each locus, see **Methods** and **Supplementary Figure 5**.

**Figure 3.**
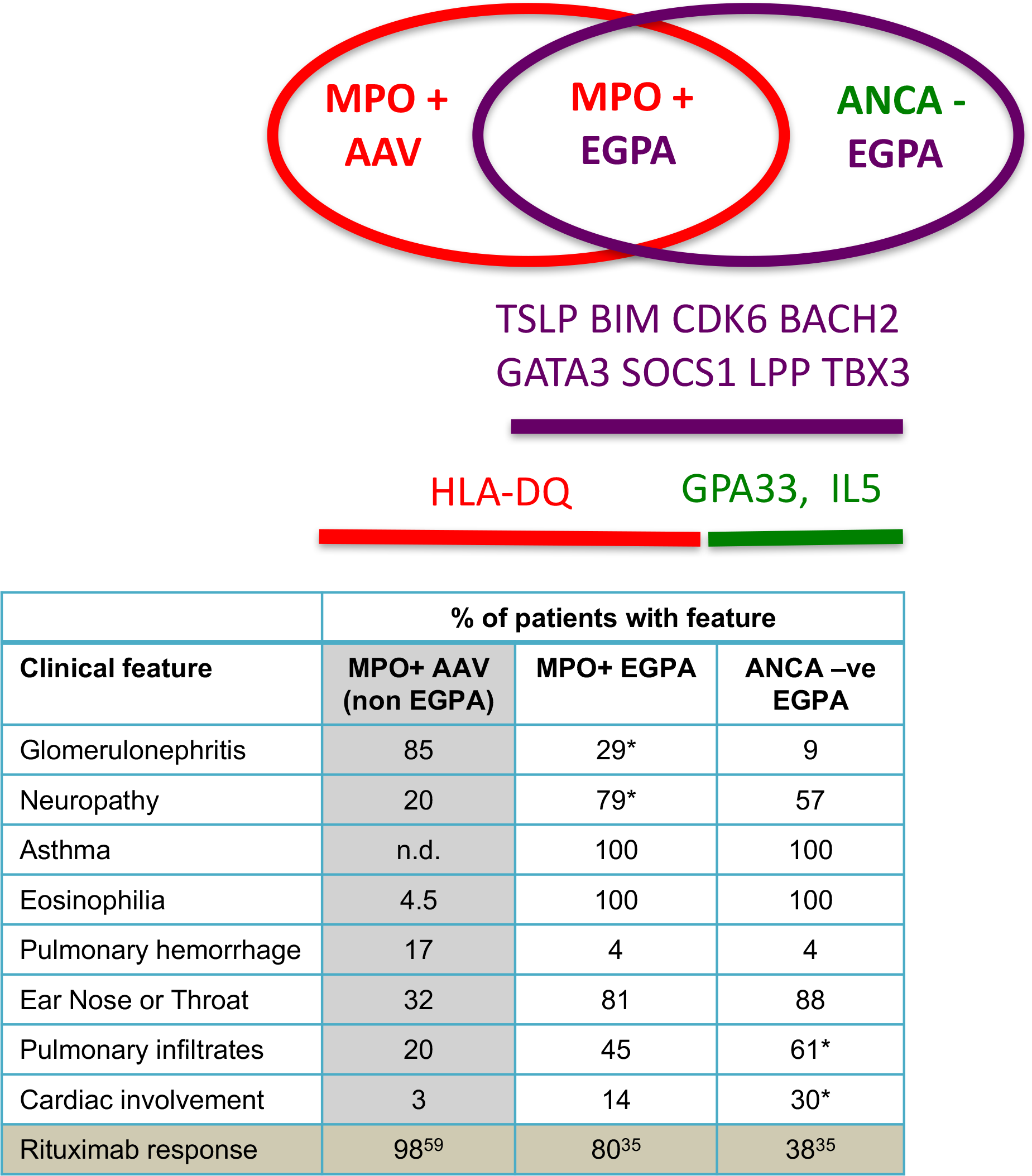
Clinically and genetically distinct subsets within EGPA, and their relation to MPO+ AAV. **Above:** schematic showing relationship between MPO+ AAV, MPO+ EGPA and ANCA -ve EGPA, and putative genes underlying this classification. **Below:** Unshaded cells in the table show a comparison of the clinical features of MPO+ANCA and ANCA-negative EGPA from this study as % (* p<0.0002 compared to other EGPA subset: see Table 1), but also see(4, 5). Shaded cells show data from external sources: MPO-AAV clinical data was derived from the EVGC AAV GWAS(15), and rituximab response rates for MPO-AAV from the RAVE study(59) and for EGPA from (35). n.d. = not determined.

### Candidate genes and EGPA pathogenesis

All genetic loci implicated in EGPA are detailed in **Supplementary Figures 6 and 7**, and four exemplars are shown in **Figure 2**. We cross-referenced the lead EGPA-associated variant at each locus with disease-associated variants in linkage disequilibrium from the NHGRI GWAS Catalog (**Supplementary Data Item 1**). It is interesting that 9 of the 11 alleles associated with increased EGPA risk are also associated with increased “physiological” eosinophil count at genome-wide significance (**Table 2**, **Figure 4A**), a correlation that persisted when all eosinophil-associated variants were assessed for EGPA risk (**Supplementary Figure 8**). In addition, 5 non-MHC EGPA risk alleles also confer increase risk of asthma (**Supplementary Table 7)**, and 4 confer increased risk of nasal polyps (**Supplementary Table 11, Supplementary Data Item 1**), consistent with the association between EGPA and these traits.

**Figure 4.**
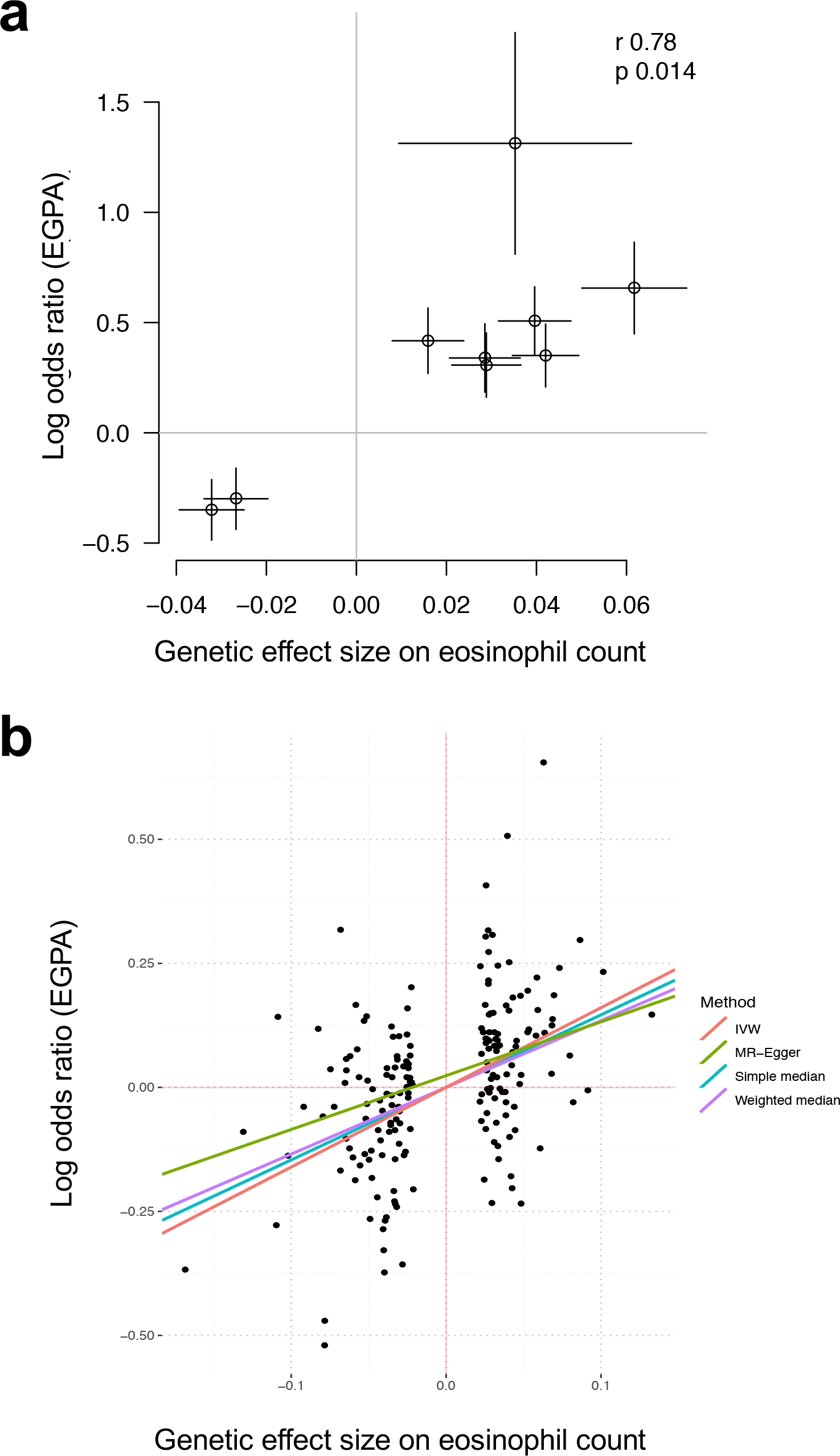
Relationship between genetic control of eosinophil count and risk of EGPA. (**A**) Correlation between the effect on EGPA risk and eosinophil count for the lead EGPA-associated genetic variants outside the *HLA* region (*GPA33* region variant rs72689399 was not present in eosinophil count dataset). Horizontal and vertical lines indicate 95% confidence intervals. (**B**) Mendelian randomization analysis supports a causal role for eosinophil abundance in EGPA aetiology. Points represent genome-wide significant conditionally independent variants associated with blood eosinophil count in the GWAS by Astle *et al*. (where typed or reliably imputed in the EGPA dataset). Coloured lines represent estimated causal effect of eosinophil count on risk of EGPA from Mendelian randomisation (MR) methods. IVW = inverse-variance weighted.

To identify potential candidate genes, we sought long-range interactions between the EGPA-associated SNP and gene promoters and regulatory regions in promoter capture Hi-C datasets using the CHiCP browser (**Methods**). We emphasise that whilst Hi-C data can suggest a link between a disease-associated variant and a candidate gene, it does not provide conclusive evidence of causality. In addition, we identified genes for which the sentinel EGPA-associated variants (or their proxies in LD) are eQTLs (**Supplementary Data Item 2**).

We sought external evidence from genomic databases (**Supplementary Data Items 1-3**) and the experimental literature (**Supplementary Table 12**) to provide corroboration for the candidate genes that we identified. For the majority of loci, we identified strong experimental evidence to implicate candidate genes in the pathogenesis of EGPA. Individual loci are discussed below.

#### BCL2L11

The sentinel EGPA-associated variant lies in an intron in *BCL2L11*, that encodes BIM, a Bcl2 family member essential for controlling apoptosis, immune homeostasis and autoimmune disease(17–19), and mast cell survival(20). Hi-C data showed interaction of the EGPA-associated variant with the promoter of *MORRBID*, that encodes a long non-coding RNA that regulates Bim transcription, controls eosinophil apoptosis and may be dysregulated in hypereosinophilic syndrome(21). These relevant functional associations suggest *BCL2L11* and *MORRBID* are more likely than *ACOXL* to be the causal gene at this locus. The EGPA risk allele is also associated with higher eosinophil count, and with increased risk of asthma, primary sclerosing cholangitis (PSC) and inflammatory bowel disease (**Figure 2A**, **Supplementary Table 11, Supplementary Data Item 1**), diseases in which eosinophils have been implicated.

#### TSLP

The EGPA susceptibility variant rs1837253 lies immediately upstream of *TSLP*. TSLP is released by stromal and epithelial cells in response to inflammatory stimuli, and drives eosinophilia and enhanced TH2 responses through effects on mast cells, group 2 innate lymphoid cells (ILC2), and dendritic cells. The risk allele, rs1837253:C, is associated with higher TSLP protein secretion in stimulated nasal epithelial cultures(22). No SNPs are in high linkage disequilibrium (LD) with rs1837253, with no variants with r2 >0.3 in European-ancestry populations in the 1000 Genomes phase 3 data, suggesting that it is likely to be the causal variant (or, less likely, that rs1837253 is tagging a rare variant that was not present in the individuals sequenced in the 1000 Genomes Project). rs1837253:C is also associated with higher risk of asthma, nasal polyps, and allergic rhinitis and with higher eosinophil counts (**Supplementary Table 11, Supplementary Data Item 1**), increasing the risk of EGPA more strongly than it does asthma (OR 1.7 vs. 1.12-1.27: **Supplementary Figure 9**). Other variants in the *TSLP* region, independent of rs1837253, are associated with asthma, eosinophil count, eosinophilic oesophagitis and allergic traits (**Supplementary Figure 6**).

#### GPA33

*GPA33* encodes a cell surface glycoprotein that maintains barrier function in the intestinal epithelium(23). Intriguingly, the EGPA-associated SNP correlates with GPA33 expression in bronchial tissue(24), suggesting GPA33 control of respiratory or intestinal barrier function might play a role in EGPA pathogenesis. In keeping with this hypothesis, we find shared genetic architecture between ANCA-negative EGPA and IBD, a disease commonly attributed to mucosal barrier dysfunction. This association was not seen with MPO+ EGPA, which is not associated with the *GPA33* variant (**Supplementary Figure 10**).

#### LPP

EGPA-associated rs9290877 is within *LPP*, encoding a LIM domain protein(25), and is associated with asthma, allergy and plasma IgE, (**Supplementary Table 11**). CHiCP analysis links the SNP to *BCL6* (**Figure 2B**), encoding a transcriptional repressor central to immune, and in particular TH2, regulation; BCL6-deficient mice die of overwhelming eosinophilic inflammation characterized by myocarditis and pulmonary vasculitis(26).

#### C5orf56-IRF1-IL5

rs11745587:A is associated with increased susceptibility to EGPA, higher eosinophil count(13), asthma (including the severe asthma subtype) and allergic rhinitis, as well as IBD and juvenile idiopathic arthritis (**Supplementary Table 11**). CHiCP analysis shows interactions with the putative regulatory regions of *IL4*, *IL5* and *IRF1*, all excellent candidates (**Figure 2C**). IL-4 and IL5 are archetypal Th2 cytokines, and IL-5 in particular drives eosinophilic inflammation(27).

#### BACH2

*BACH2* encodes a transcription factor that plays critical roles in B and T cell biology, BACH2 deficient mice die of eosinophilic pneumonitis(28), and polymorphisms in the *BACH2* region have been associated with susceptibility to numerous immune-mediated diseases (**Supplementary Table 11, Supplementary Figure 6**). The EGPA-associated variant is in LD with variants associated with asthma, nasal polyps, and allergy, as well as other immune-mediated diseases including celiac disease, IBD, PSC and multiple sclerosis (**Supplementary Data Item 1**).

#### CDK6

*CDK6* encodes a protein kinase that plays a role in cell cycle regulation. The SNP most associated with EGPA, rs42046, is also associated with eosinophil count(13) and rheumatoid arthritis (**Supplementary Table 11**).

### 10p14 intergenic region and GATA3

The associated SNP is not near any candidate genes, but CHiCP analysis shows interaction with the promoter of *GATA3* (**Figure 2D**). *GATA3* encodes a master regulator transcription factor expressed by immune cells including T, NK, NKT and ILC2 cells, that drives Th2 differentiation and secretion of IL-4, IL-5, and IL-13, and thus eosinophilic inflammation(29). The EGPA risk variant is also associated with asthma and allergic rhinitis (**Supplementary Table 11**, **Supplementary Data Item 1**).

#### TBX3-MED13L

*TBX3* encodes a transcriptional repressor important in embryonic development, and *MED13L* a subunit of a transcriptional co-activator. Neither have any known role in immune-mediated disease.

#### CLEC16A-SOCS1

CHiCP analysis shows links with a number of plausible candidate genes, including *CLEC16A* (implicated in autoimmunity in non-obese diabetic mice(30)), *SOCS1* (negative regulator of cytokine signaling associated with inflammation(31)) and DEXI (associated with Type I diabetes(32)). The EGPA-associated variant is associated with asthma, nasal polyps, and PSC (**Supplementary Table 11**).

#### HLA

MPO+ANCA EGPA was associated with a region encompassing the *HLA-DR* and –*DQ* loci (**Supplementary Figure 11**). The classical *HLA* alleles at 2 or 4 digit resolution, and amino acid variants at 8 *HLA* loci, were then imputed. Univariate logistic regression identified 8 *HLA* alleles conferring susceptibility to MPO+ANCA EGPA (**Supplementary Table 13**). Reciprocal stepwise conditional analysis revealed 3 signals conferred by 2 extended haplotypes encoding either *HLA-DRB1*0801-HLA-DQA1*04:01-HLA-DQB1*04:02*; or *HLA-DRB1*07:01-HLA-DQA1*02:01- HLA-DQB1*02:02/HLA-DQB1*03:03*; together with an additional signal at *HLA-DRB1*01:03* (**Supplementary Table 13** and **Supplementary Figure 11**). The strongest independent associations with disease risk were seen at *HLA-DQA1*04:01* (OR 7.18, p = 9.1×10^−20^), *HLA-DQA1*02:01* (OR 3.05, p = 2.0×10^−11^) and *HLA-DRB1*01:03* (OR 5.96, p = 2.3×10^−7^). Analysis of *HLA* allelic frequencies stratified by country of recruitment revealed a consistent pattern (**Supplementary Table 14**), indicating that our findings were not the result of residual population stratification.

Individual amino acid variants in *HLA-DRB1*, *HLA-DQA1*, and *HLA-DQB1* were associated with disease. Conditioning on the most significant (absence of Threonine at position 181 in *HLA-DRB1*; OR=4.5, P=3.3×10^−24^) eliminated most of the effect at all other variants (**Supplementary Figure 12**) and all the classical alleles except *HLA-DRB1*01:03* (OR=9.3, P=5.0×10^−10^ following conditioning). Conditioning on both *HLA-DRB1*01:03* and position 181 in *HLA-DRB1* accounted for the entire signal seen at the MHC locus. The *HLA-DQ* locus associated with MPO+ANCA EGPA was the same as that previously associated with MPO+ANCA vasculitis(15). To quantify the relative contribution of the MHC to the heritable phenotypic variance we partitioned the variance using BOLT-REML (33). Using this approach, 6% of the heritable variance could be attributed to the MHC, an observation similar to that seen in other autoimmune diseases where the MHC contribution ranges from 2% in systemic lupus erythematosus to 30% in type 1 diabetes (34).

### Mendelian randomisation supports a causal role for eosinophils in EGPA

The observation that eosinophilia is a ubiquitous clinical feature in EGPA does not necessarily imply that eosinophils play a causal role in the disease, since eosinophilia might instead be either an epiphenomenon or a downstream consequence of EGPA. The observation that 9 of the 11 alleles associated with increased EGPA risk are also associated with increased “physiological” eosinophil count, however, supports the notion of a causal role for eosinophils in EGPA. To formally test this hypothesis, we employed the technique of Mendelian randomisation (MR) (**Methods**). In contrast to observational associations, which are liable to confounding and/or reverse causation, MR analysis is akin to a “natural” randomised trial, exploiting the random allocation of alleles at conception to allow causal inference. MR analysis provided strong support (P < 2.8 ×10^−9^, inverse-variance weighted method) for a causal effect of eosinophil count on EGPA risk (**Figure 4B**). This result was robust to a number of sensitivity analyses (**Methods**, **Supplementary Figure 13, Supplementary Data Item 4**)

## Discussion

This study has identified 11 loci associated with EGPA, and reveals genetic and clinical differences between the MPO+ANCA EGPA subset, and the larger ANCA-negative subset (with PR3+ANCA patients too rare to be informative). There was a strong association of the MPO+ANCA subset with *HLA-DQ*, and no *HLA* association with ANCA-negative EGPA. The ANCA-negative group alone was associated with variants at the *GPA33* and *IL5/IRF1* loci. There was clear evidence of association of both EGPA subgroups with variants at the *TSLP, BCL2L11* and *CDK6* loci, and suggestive evidence for *BACH2*, Chromosome 10, *SOCS1*, *LPP* and *TBX3*. A number of these loci have previously been shown to be associated with other autoimmune diseases, including PSC (**Supplementary Table 11**). Thus EGPA is characterised by certain genetic variants that associate with the syndrome as a whole, but others that indicate a genetic distinction between the MPO+ANCA and ANCA-negative subsets. This distinction suggests important heterogeneity in the pathogenesis of the clinical syndrome, as the subsets have distinct clinical phenotypes and outcomes to therapy (**Figure 3**). The increased efficacy of rituximab in the MPO+ANCA subset of EGPA (35) demonstrates that trials of novel therapies might need to be stratified according to ANCA.

EGPA shares susceptibility variants with asthma (**Table 2**, **Supplementary Table 11**), and this genetic relatedness (**Supplementary Figure 3**) allowed the use of the cFDR technique to detect additional EGPA-associated variants. For the *TSLP* variant, the estimated effect size in EGPA was much greater than in asthma, with non-overlapping confidence intervals (**Supplementary Figure 9**), suggesting its association with asthma might be driven by a subset of patients. Consistent with this, rs1837253 (near *TSLP*) shows a trend towards association with a subset of severe asthmatics(12), as does the EGPA-associated variant at *C5orf56*/*IL5*. This raises the possibility that common EGPA-associated variants may be particularly associated with severe, adult-onset asthma– the asthma endotype typical of the EGPA prodrome. All or some of this subset of asthma patients may be at higher risk of subsequently developing EGPA.

Notably, 9 of 11 variants associated with EGPA were also associated with eosinophil count in normal individuals. The increase in EGPA risk conferred by each of the 9 genetic variants was proportional to its effect on eosinophil count (**Figure 4A**), a correlation that held true when all eosinophil-associated variants were assessed for EGPA risk (**Supplementary Figure 8**). The causal nature of this association was supported by Mendelian randomisation analysis. This dose-response relationship between the genetic effects on eosinophil count and on risk of EGPA is consistent with a scenario where the genetic control of “physiological” variation in the eosinophil count contributes directly to the risk of pathological eosinophilia and eosinophilic inflammatory disease. This inference is possible because the genetic effects on eosinophil count were derived in an independent dataset of healthy individuals (and thus the eosinophil count in this cohort is not confounded by the presence of disease). This is a novel concept that might extend beyond eosinophil-driven conditions – that genetic variants that control a cell number within the “normal range” might, when combined, predispose to and perhaps initiate disease. By analogy to studies establishing the relationship between genetic associations with LDL-C and with coronary artery disease risk(36), our data supporting a causal role for eosinophils could pave the way for eosinophil-targeted therapies in EGPA.

The shared eosinophil count and asthma-associated variants may explain the shared clinical features of the two EGPA subsets – both in the prodromal phase and after the development of frank EGPA. Moreover, they raise the possibility that persistent eosinophilia driven by variation in genes controlling “physiological” eosinophil levels can predispose to, or even directly give rise to, adult-onset asthma and to the EGPA prodrome (or to a progression from adult-onset asthma to the eosinophilic prodrome). After some time, an unknown proportion of people with this prodrome proceed to develop the clinical features characteristic of EGPA. Whether this progression is stochastic, is related to the severity of ongoing eosinophilic inflammation, is influenced by genetic factors or is triggered by an environmental factor is unknown.

Around a third of EGPA patients develop MPO+ANCA and clinical features that overlap with MPO+AAV, such as vasculitis and necrotising glomerulonephritis (**Figure 3**). These patients appear to have a classic *HLA* class II-associated autoimmune disease with prominent eosinophilic features. The remaining patients, in contrast, develop disease characterised more by tissue eosinophilia, and its cardiac and pulmonary manifestations, in the absence of both autoantibodies and an *HLA* association. This subgroup has a genetic association with expression of the barrier
protein *GPA33* (24) and shared genetic architecture with IBD, suggesting that ANCA-negative EGPA might arise as a mucosal/barrier, rather than autoimmune, disease.

EGPA has traditionally been treated with similar non-selective immunosuppression to AAV, such as steroids, cyclophosphamide or rituximab(2, 35). This study suggests investigation of further therapies may be warranted. Anti-IL5 (mepolizumab) has been developed for severe asthma, and efficacy in EGPA was recently confirmed in the MIRRA study(37). Given the genetic variant at *IRF1/IL5* is associated with ANCA-negative EGPA, it would be interesting to specifically analyse this subset. Anti-TSLP agents are undergoing trials in asthma and might be considered in EGPA. BCL2 antagonists (e.g. ABT737) act by disrupting the sequestration of BIM by BCL2 and are being developed as cancer chemotherapeutics(38). Their immunomodulatory effect(39) and exquisite sensitivity for driving mast cell apoptosis(40) suggests they may be effective in EPGA at low, perhaps non-toxic, doses. CDK6 inhibitors and SOCS1 mimetics are also under development, and their impact in preclinical models(41, 42) makes them additional therapeutic candidates for EGPA.

Our study had potential limitations. Whilst the MIRRA criteria are the most suitable diagnostic criteria for this study, it is possible that some patients with idiopathic hypereosinophilic syndrome may also meet these criteria. The choice of diagnostic criteria is discussed in detail in the **Supplementary Appendix**. We observed elevation of the genomic inflation factor even after adjustment for genetic principal components, suggesting potential residual population stratification, and so we used a more stringent significance threshold adjusted for residual genomic inflation to mitigate against this. In addition the use of a linear mixed model analysis improved the value of lambda and confirmed the reported associations.

The rarity of EGPA makes GWAS challenging. Compared to GWAS of common diseases, our sample sizes were necessarily small and consequently our power was limited, particularly for uncommon or rare variants. Nevertheless, the 3 variants that achieved genome-wide significance in our primary cohort achieved nominal significance in our small replication cohort. Moreover, we observed strong correlation between estimated odds ratios between our primary and replication cohorts, with excellent agreement for 10 of the 11 loci. In addition, we were able to identify strong, and in some cases overwhelming, functional data and experimental literature to support the associations at the majority of loci identified (**Supplementary Table 12**).

In summary, this GWAS has demonstrated that EGPA is a polygenic disease. Most genes associated with EGPA are also associated with control of the normal eosinophil count in the general population, suggesting a primary tendency to eosinophilia underlies susceptibility. Given the rarity of EGPA, it is likely that additional as yet unidentified environment or genetic factors are necessary to trigger disease. After the asthma/eosinophilia prodrome, EGPA develops and comprises two genetically and clinically distinguishable syndromes with different treatment responsiveness. MPO+ANCA EGPA is an eosinophilic autoimmune disease sharing both clinical features and an MHC association with anti-MPO AAV. ANCA-negative EGPA may instead have a mucosal/barrier origin. Thus the identification of genes associated with EGPA helps explain its pathogenesis, points to logical therapeutic strategies, and supports a case for formally recognising the two distinct conditions that comprise it.

## Methods

### Inclusion criteria

The important issue of diagnostic criteria is discussed in the **Supplementary Appendix**. We used the recently developed diagnostic criteria used in the Phase III clinical trial “Study to Investigate Mepolizumab in the Treatment of Eosinophilic Granulomatosis With Polyangiitis” (MIRRA: **Supplementary Table 1**)(37). These define EGPA diagnosis based on the history or presence of *both* asthma and eosinophilia (>1.0 \times 10^9^/L and/or > 10% of leukocytes) *plus* at least two additional features of EGPA.

### Subjects

We recruited 599 individuals with a clinical diagnosis of EGPA from 17 centers in 9 European countries (**Supplementary Table 2**). Nine individuals were excluded because they did not fulfil the MIRRA criteria. In total, 542 patients were included in the GWAS after poor quality samples and patients with non-European ancestry were excluded (see ‘Genotyping and quality control’ below). Of these, 299 (55%) were female and 243 (45%) male. Clinical characteristics are shown in **Table 1**. 358 patients were ANCA negative, 161 patients were MPO-ANCA positive, and 5 were PR3-ANCA positive. For 5 patients, there was no data on ANCA status. 13 patients were ANCA positive with either no data on specific antibodies to PR3 or MPO, or with positive ANCA immunofluorescence without detectable antibodies to PR3 or MPO.

Genotype data for 6000 UK controls was obtained from the European Prospective Investigation of Cancer (EPIC) Consortium. 496 individuals with a history of asthma were excluded. After QC, 5466 individuals remained. In addition, we recruited and genotyped controls from 6 European countries (**Supplementary Table 2**).

A further 150 patients with a clinical diagnosis of EGPA that fulfilled the MIRRA criteria were recruited from Germany and Italy for replication purposes, along with 125 controls from these countries. In total, 142 cases were included in the study following the removal of poor quality samples, of these 43 were MPO ANCA +ve. All individuals provided written informed consent.

### Genotyping and quality control (QC)

Genomic DNA was extracted from whole blood using magnetic bead technologies at the Centre for Integrated Genomic Medical Research (Manchester, UK) according to manufacturer’s instructions. Patients with EGPA and healthy controls were genotyped using the Affymetrix UK Biobank Axiom array according to the manufacturer’s protocol. Genotyping of cases and non-UK controls was performed by AROS Applied Biotechnology (Aarhus, Denmark). Genotyping of UK controls had been performed previously by the EPIC-Norfolk consortium(43), also using the Affymetrix Axiom UK Biobank array.

Genotype calling was performed using the Affymetrix Powertools software. Calling was performed in batches of contemporaneously run plates as per the manufacturer’s advice. After genotype calling, SNP data was processed in the following sequence using PLINK v1.9(44). Samples with a sex mismatch, abnormal heterozygosity, or proportion of missing SNPs >5%, were removed. SNPs with missing calls >2%, deviation from Hardy-Weinberg Equilibrium (p-value <1×10^−6^), or which were monomorphic were removed. The QC process was performed separately for each batch. The post-QC genotype data from each batch was then merged. Following this merger, duplicated or related samples (identified using Identity by State) were removed, and the SNP QC was performed again so that SNPs with missing calls >2% or deviation from Hardy-Weinberg Equilibrium (p-value <1×10^−6^) in the combined data were excluded. SNPs that showed significant differential missingness (Benjamini-Hochberg adjusted p-value <0.05) between cases and controls were removed. Finally, principal components analysis (PCA) of the post-QC genotype calls combined with calls from 1000 Genome individuals was performed (**Supplementary Figure 14**). Samples of non-European ancestry by PCA were excluded as described below. The means and standard deviations of PC1 and PC2 were calculated for the EUR subset of the 1000 Genomes samples. Cases or controls lying outside +/− 3 standard deviations from the mean on either PC1 or PC2 were removed. Following these QC steps, 542 patients and 6717 controls remained (see **Supplementary Table 2**). 543,639 autosomal SNPs passed QC.

### Replication cohort genotyping

Replication cohort samples were genotyped using the Affymetrix UK Biobank Axiom array according to the manufacturer’s protocol by Cambridge Genomic Services (Cambridge, UK). Genotype calling and QC was performed as outlined above except that samples were processed as a single batch. Following these steps, 142 cases, 121 controls and 626,229 autosomal SNPs passed QC.

### Association testing on the directly genotyped data

Case-control association testing on the 543,639 directly genotyped autosomal SNPs was performed using logistic regression with the SNPTEST software. Principal components (PCs) were included as covariates to adjust for confounding factors. We calculated PCs on the genotype matrix containing both cases and controls. Whilst this carries a small risk of masking true disease associations, we felt that this more conservative approach was appropriate since i) the UK controls were genotyped in a different facility thereby potentially leading to confounding from a batch effect, and ii) we did not have French controls. QQ plots of expected test statistics under the null hypothesis of no genotype-disease association versus the observed test statistics are shown in **Supplementary Figure 15**. The genomic inflation factor (lambda, the ratio of the median of the observed chi-squared statistics to the median expected chi-squared statistic under the null hypothesis) was calculated. We assessed the effect of using increasing numbers of PCs as covariates on lambda. Using the first 20 PCs produced a reduction in lambda to 1.09, with little benefit from the inclusion of further PCs.

In addition to testing all EGPA cases versus controls, we also tested MPO-ANCA positive cases (n=161) against controls, and ANCA negative cases (n= 358) against controls. 5 individuals who were PR3-ANCA positive and 18 individuals in whom we had either no data on ANCA status, or who were ANCA positive by immunofluorescence but in whom specific antibodies to MPO and PR3 were negative or unknown were excluded from these subset analyses.

### Pre-phasing and imputation

Imputation was performed using the 1000 genomes phase 3 individuals as a reference panel (data download date 7^th^ July 2015). Pre-phasing was first performed with SHAPEIT(45), and then imputation was performed using IMPUTE2(46, 47). For the IMPUTE2 Monte Carlo Markov Chain algorithm we used the default settings (30 iterations, with the first 10 discarded as the ‘burn-in’ period). The ‘k_hap’ parameter was set to 500. IMPUTE2 was provided with all available reference haplotypes from the 1000 Genomes individuals, as the software chooses a custom reference panel for each individual to be imputed.

### Association testing following imputation

Association testing on the imputed data was carried out using the SNPTEST software. We used an additive genetic model (option ‘--frequentist 1’), with the first 20 PCs as covariates to adjust for population stratification. Uncertainty in the imputed genotypes was taken into account in the association testing by using a missing data likelihood score test. We limited association testing of imputed SNPs to those with an ‘info’ metric of 0.9 or higher. An info metric near 1 indicates that a SNP has been imputed with high confidence. 7,585,056 autosomal SNPs either directly typed or imputed with an info metric greater than 0.9 were taken forward for association testing. The genomic inflation factor for the association tests using the imputed data was 1.10 for the primary cohort and 1.07 for the replication cohort. The results of the two cohorts were meta-analysed using the META software using an inverse-variance method based on a fixed-effects model (option ‘—method 1’).

### Adjustment of genome-wide level of significance to account for residual genomic inflation

Since lambda was 1.10 despite inclusion of 20 PCs as covariates, we computed an adjusted genome-wide significance threshold accounting for residual genomic inflation, using Devlin and Roeder’s method(48). An adjusted chi-squared statistic threshold was calculated by multiplying the 1 degree of freedom chi-squared corresponding to a p-value of 5 x 10^−8^ by lambda (1.10). The p-value corresponding to this, 1.0×10^−8^, is the adjusted significance threshold. Use of this threshold on unadjusted p-values is equivalent to using a threshold of 5 x 10^−8^ on adjusted p-values. A potential limitation of this genomic inflation correction procedure is that it assumes the level of inflation is uniform across the genome.

### Association testing using a linear mixed model to correct for population stratification

As an alternative strategy to mitigate against the effects of population stratification, we performed the GWAS using a linear mixed model with the LMM-BOLT software (49). This approach resulted in an improved lambda value of 1.047.

### Fine-mapping

To identify the most likely causal variant we performed fine-mapping as follows. We computed approximate Bayes factors for each variant using the Wakefield approximation(50), with a prior parameter W=0.09, indicating our prior expectation that true effect sizes (relative risks) exceed 2 only 1% of the time. Thus, we calculated posterior probabilities that each variant is causal, assuming a single causal variant per LD-defined region as previously proposed(51). Code to perform these steps is available at https://github.com/chr1swallacw/lyons-et-al.

### Haplotype block estimation

Haplotype blocks around the EGPA SNPs reported in Table 2 were calculated using the “–blocks” option in PLINK v1.9, based on individuals of European ancestry from the 1000 Genomes Project phase 3 data. PLINK v1.9 calculates haplotype blocks based on Haploview’s interpretation of the block definition suggested by Gabriel et al (52). The following options were used: --blocks-max-kb 500 -blocks-strong-lowci 0.7005 –blocks-min-maf 0.01 –window 500.

### Leveraging association statistics from related traits

In order to increase power to detect EGPA-associated genetic variants, we used a conditional FDR (cFDR) method(10, 11) to leverage findings from other GWAS studies of related phenotypes. We used the summary statistics from a GWAS of asthma(12) and a population-scale GWAS of peripheral blood eosinophil counts(13) to calculate a conditional FDR for each SNP for association with EGPA conditional on each of these other traits.

Given p-values *P*_*i*_ for EGPA and *P*_*j*_ for the conditional trait (eg asthma), and a null hypothesis of no association with EGPA *H*_*0*_, the cFDR for p-value thresholds *p*_*i*_ and *p*_*j*_ is defined as

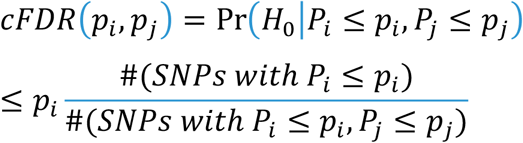

roughly analogous to Storey’s q-value computed only on the SNPs for which *P*_*j*_ ≤ *p*_*j*_.

The cFDR has the advantage of asymmetry, only testing against one phenotype at a time. Intuitively, if we know that associations are frequently shared between asthma and EGPA, and attention is restricted to a set of SNPs with some degree of association with asthma, we may relax our threshold for association with EGPA. The cFDR formalizes this intuition in a natural way, with the adjustment in the threshold responding to the total degree of observed overlap between disease associations.

The cFDR lacks a convenient property of the Q-value limiting the overall FDR to the significance threshold for the test statistic. Namely, if we reject *H*_*0*_ at all SNPs with cFDR < *α*, the overall FDR is not necessarily bound above by *α*(11). An upper bound on the overall FDR can be obtained by considering the region of the unit square for which *p*_*i*_, *p*_*j*_ reaches significance, with the bound typically larger than *α*.

We used a threshold = cFDR (1×10^−8^,1), chosen to be the most stringent threshold for which all SNPs reaching genome-wide significance (following adjustment for residual genomic inflation, hence p_i_= 1×10^−8^ rather than 5×10^−8^) in a univariate analysis will also reach significance in the cFDR analysis. The value α is equal to the FDR for this univariate analysis on the set of SNPs for which the cFDR is defined. Put more intuitively, cFDR thresholds for the analyses conditioning on asthma and eosinophilia were chosen so that the overall FDR was less than the FDR corresponding to a P value for (adjusted) genome-wide significance in the standard univariate analysis.

Clearly, the cFDR method requires that genotype data for a given SNP is available for both traits. The cFDR analysis was limited to SNPs that were directly typed in each GWAS. The cFDR analysis for EGPA conditional on asthma examined 74,776 SNPs and that for EGPA conditional on eosinophil count 513,801.

### HLA imputation

Two thousand seven hundred and seventy seven SNPs, 343 classical HLA alleles to 2 or 4 digit resolution and 1,438 amino acid variants were imputed at 8 HLA loci (*HLA-A*, *HLA-B*, *HLA-C*, *HLA-DRB1*, *HLA-DQA1*, *HLA-DQB1*, *HLA-DPA1* and *HLA-DPB1*) from phased genotype data using the HLA*IMP:03 software(53). Association testing for HLA variants was performed in Plink v1.9 using logistic regression adjusting for population structure.

### Narrow-sense heritability estimation

The narrow-sense heritability (h^2^) of EGPA was estimated using LD-score regression(54). LD scores were calculated using the European 1000 Genomes Project reference panel. To estimate the contribution of the MHC to heritable phenotypic variance we performed a variance-components analysis using BOLT-REML(33).

### Promoter enhancer interaction mining

Long-range interactions between genetic variants associated with EGPA and gene promoter and regulatory regions were identified in promoter capture Hi-C datasets from a range of primary cell types(55) and cell lines(56) using the CHiCP browser(57).

### Data mining

Other traits associated with EGPA-associated loci were identified using PhenoScanner(58). Phenoscanner searches GWAS data from multiple sources including the NHGRI-EBI (National Human Genome Research Institute-European Bioinformatics Institute) GWAS Catalog and NHLBI GRASP (Genome-Wide Repository of Associations Between SNPs and Phenotype) Catalog, and accounts for LD between queried SNPs and those in the catalogs of trait-associated variants. For each locus associated with EGPA, we searched for traits associated with SNPs in LD (r^2^ 0.6 or higher) with the lead EGPA SNP. We searched for all associations with a p-value of 1 ×10^−5^ or lower so that in addition to identifying all associations that have achieved ‘genome-wide’ significance, we also identified associations that are ‘suggestive’ or ‘sub-genome wide’. We also used PhenoScanner to identify eQTLs at EGPA-associated loci.

### Associations of clinical characteristic with ANCA status

Comparisons of clinical features between MPO+ANCA EGPA patients (n=161) and ANCA negative patients (n=358) were performed from 2×2 contingency tables using 1 degree of freedom chi-squared tests with Yates’ continuity correction. Bonferroni correction of P values was undertaken to account for the multiple testing of 8 clinical features. Associations with PR3 antibodies were not assessed as this subgroup (n=5) was too small for statistical analysis. Patients with missing or incomplete ANCA data, and those who were ANCA positive by immunofluorescence without MPO or PR3 antibodies were excluded from this analysis.

ANCA status was tested for association with country of origin using a chi-squared test on the 8×2 contingency table (**Supplementary Table 8**). Columns represented ANCA status (ANCA negative or MPO positive) and rows represented country of origin. The Republic of Ireland and the UK were considered as one entity for the purposes of this analysis. We found a significant association between ANCA status and country of origin (P 1.9×10^−10^). Since there were only 5 cases from the Czech Republic, in sensitivity analysis, we repeated the chi-squared test after merging the counts for Czech and German cases. This did not materially affect the association (P 5.0 ×10^−10^). This association is likely to reflect the differing specialities of recruiting centres (ANCA positive patients are more likely to be found in nephrology clinics than rheumatology clinics). Nevertheless, to ensure that the clinical associations that we identified with ANCA status were not in fact driven by geographical differences, we repeated the association testing adjusting for country of origin. We did this by performing logistic regression of each clinical feature on ANCA status (MPO +ve versus negative), with country of origin (coded using dummy variables) as a covariate (**Supplementary Table 8**).

### Mendelian Randomisation

“Two-sample” Mendelian randomisation (MR) was performed to assess whether there is a causal effect of eosinophil count (the exposure) on EGPA (the outcome). MR analysis was performed using the MendelianRandomization R package. Summary statistics were obtained for 209 conditionally independent variants associated with peripheral blood eosinophil count in a population study by Astle et al **(13)**. 193 of these were typed or imputed in the EGPA dataset. MR analysis was performed using the beta coefficients and standard errors for these 193 variants in the eosinophil count GWAS and in our EGPA GWAS (all EGPA versus controls). The primary analysis was conducted using the inverse-variance weighted method. Additional sensitivity analyses were performed using alternative methods (**Supplementary Figure 13**).

### Study approval

Written informed consent was received from participants prior to inclusion in the study. Details of ethical approval for each participating center are shown in **Supplementary Table 3**.

## Supporting information

## Author Contributions

KGCS and PAL conceived and designed the study. PAL, JEP, FA, JL, RMRC, SL, DV, WA, HE, TJ and CW analysed data. FA, CB, MCC, JE, LG, DRWJ, IG, PL, MAL, DM, FM, SO, GAR, BR, GS, RAS, WS, VT, BT, RAW, AV, JUH and the EVGC provided samples and collected clinical data. KGCS wrote the manuscript with the assistance of PAL, JEP and JL. All authors read and approved the final version of the manuscript.

## Acknowledgements

This work was funded primarily by Project Grants from Arthritis Research UK (20593 to Drs. Smith and Lyons) and the British Heart Foundation (PG/13/64/30435 to Drs. Smith and Lyons). Additional support was provided by the NIHR Cambridge Biomedical Research Centre, the West Anglia Comprehensive Research Network, a Medical Research Council Programme Grant (MR/L019027/1 to Dr. Smith), a Wellcome Trust Investigator Award (200871/Z/16/Z to Dr. Smith), a NIHR Senior Investigator Award (to Dr. Smith), a Wellcome Trust Senior Research Fellowship (WT107881 to Dr. Wallace), a Medical Research Council grant (MC_UU_00002/4 to Dr. Wallace), a Wellcome Trust Mathematical Genomics and Medicine Programme Studentship (to Dr. Liley), a Career Development Award from the Cambridge British Heart Foundation Centre for Research Excellence and a UK Research Innovation Fellowship (RE/13/6/30180 and MR/S004068/1 to Dr. Peters). Additional aspects of this work were supported by the following funding: a Science Foundation Ireland Grant (11/Y/B2093 to Dr Little); an Australian National Health and Medical Research Council (NH&MRC), Career Development Fellowship (ID 1053756) and a Victorian Life Sciences Computation Initiative (VLSCI) grant number (VR0240) on its Peak Computing Facility at the University of Melbourne, an initiative of the Victorian Government, Australia (to Dr Leslie); Research at the Murdoch Children’s Research Institute was supported by the Victorian Government’s Operational Infrastructure Support Program; Project RVO 64 165 of the Ministry of Health of Czech Republic (to Dr Tesar); Ministerio de Economía y Competitividad (SAF 2014-57708-R), FEDER una manera de hacer Europa) and CERCA programme (to Drs Cid and Hernández-Rodríguez) and Instituto de Salud Carlos III (PI 15/00092) (to Drs Espígol-Frigolé and Prieto-Gonzalez); Prof Bruce is an NIHR Senior Investigator and is funded by Arthritis Research UK, the National Institute for Health Research Manchester Biomedical Research Unit and the NIHR/Wellcome Trust Manchester Clinical Research Facility.

We thank the Centre for Integrated Genomic Medical Research Biobank for sample storage and preparation, and AROS Applied Biotechnology (Aarhus, Denmark) for genotyping.

See **Supplementary Appendix** for consortium details.

## References

1. Churg J, and Strauss L. Allergic granulomatosis, allergic angiitis, and periarteritis nodosa. Am J Pathol. 1951;27(2):277–301.

2. Mahr A, Moosig F, Neumann T, Szczeklik W, Taille C, Vaglio A, and Zwerina J. Eosinophilic granulomatosis with polyangiitis (Churg-Strauss): evolutions in classification, etiopathogenesis, assessment and management. Curr Opin Rheumatol. 2014;26(1):16–23.

3. Jennette JC, Falk RJ, Bacon PA, Basu N, Cid MC, Ferrario F, Flores-Suarez LF, Gross WL, Guillevin L, Hagen EC, et al. 2012 revised International Chapel Hill Consensus Conference Nomenclature of Vasculitides. Arthritis Rheum. 2013;65(1):1–11.

4. Sable-Fourtassou R, Cohen P, Mahr A, Pagnoux C, Mouthon L, Jayne D, Blockmans D, Cordier JF, Delaval P, Puechal X, et al. Antineutrophil cytoplasmic antibodies and the Churg-Strauss syndrome. Ann Intern Med. 2005;143(9):632–8.

5. Sinico RA, Di Toma L, Maggiore U, Bottero P, Radice A, Tosoni C, Grasselli C, Pavone L, Gregorini G, Monti S, et al. Prevalence and clinical significance of antineutrophil cytoplasmic antibodies in Churg-Strauss syndrome. Arthritis Rheum. 2005;52(9):2926–35.

6. Sada KE, Amano K, Uehara R, Yamamura M, Arimura Y, Nakamura Y, Makino H, and Research Committee on Intractable Vasculitides tMoHLWoJ. A nationwide survey on the epidemiology and clinical features of eosinophilic granulomatosis with polyangiitis (Churg-Strauss) in Japan. Mod Rheumatol. 2014;24(4):640–4.

7. Vaglio A, Martorana D, Maggiore U, Grasselli C, Zanetti A, Pesci A, Garini G, Manganelli P, Bottero P, Tumiati B, et al. HLA-DRB4 as a genetic risk factor for Churg-Strauss syndrome. Arthritis Rheum. 2007;56(9):3159–66.

8. Wieczorek S, Hellmich B, Gross WL, and Epplen JT. Associations of Churg-Strauss syndrome with the HLA-DRB1 locus, and relationship to the genetics of antineutrophil cytoplasmic antibody-associated vasculitides: comment on the article by Vaglio et al. Arthritis Rheum. 2008;58(1):329–30.

9. Chen GB, Lee SH, Brion MJ, Montgomery GW, Wray NR, Radford-Smith GL, Visscher PM, and International IBDGC. Estimation and partitioning of (co)heritability of inflammatory bowel disease from GWAS and immunochip data. Hum Mol Genet. 2014;23(17):4710–20.

10. Andreassen OA, Thompson WK, Schork AJ, Ripke S, Mattingsdal M, Kelsoe JR, Kendler KS, O’Donovan MC, Rujescu D, Werge T, et al. Improved detection of common variants associated with schizophrenia and bipolar disorder using pleiotropy-informed conditional false discovery rate. PLoS Genet. 2013;9(4):e1003455.

11. Liley J, and Wallace C. A pleiotropy-informed Bayesian false discovery rate adapted to a shared control design finds new disease associations from GWAS summary statistics. PLoS Genet. 2015;11(2):e1004926.

12. Moffatt MF, Gut IG, Demenais F, Strachan DP, Bouzigon E, Heath S, von Mutius E, Farrall M, Lathrop M, Cookson WO, et al. A large-scale, consortium-based genomewide association study of asthma. N Engl J Med. 2010;363(13):1211–21.

13. Astle WJ, Elding H, Jiang T, Allen D, Ruklisa D, Mann AL, Mead D, Bouman H, Riveros-Mckay F, Kostadima MA, et al. The Allelic Landscape of Human Blood Cell Trait Variation and Links to Common Complex Disease. Cell. 2016;167(5):1415–29 e19.

14. Comarmond C, Pagnoux C, Khellaf M, Cordier JF, Hamidou M, Viallard JF, Maurier F, Jouneau S, Bienvenu B, Puechal X, et al. Eosinophilic granulomatosis with polyangiitis (Churg-Strauss): clinical characteristics and long-term followup of the 383 patients enrolled in the French Vasculitis Study Group cohort. Arthritis Rheum. 2013;65(1):270–81.

15. Lyons PA, Rayner TF, Trivedi S, Holle JU, Watts RA, Jayne DR, Baslund B, Brenchley P, Bruchfeld A, Chaudhry AN, et al. Genetically distinct subsets within ANCA-associated vasculitis. N Engl J Med. 2012;367(3):214–23.

16. Millet A, Pederzoli-Ribeil M, Guillevin L, Witko-Sarsat V, and Mouthon L. Antineutrophil cytoplasmic antibody-associated vasculitides: is it time to split up the group? Ann Rheum Dis. 2013;72(8):1273–9.

17. Bouillet P, Metcalf D, Huang DC, Tarlinton DM, Kay TW, Kontgen F, Adams JM, and Strasser A. Proapoptotic Bcl-2 relative Bim required for certain apoptotic responses, leukocyte homeostasis, and to preclude autoimmunity. Science. 1999;286(5445):1735–8.

18. Bouillet P, Purton JF, Godfrey DI, Zhang LC, Coultas L, Puthalakath H, Pellegrini M, Cory S, Adams JM, and Strasser A. BH3-only Bcl-2 family member Bim is required for apoptosis of autoreactive thymocytes. Nature. 2002;415(6874):922–6.

19. Enders A, Bouillet P, Puthalakath H, Xu Y, Tarlinton DM, and Strasser A. Loss of the pro-apoptotic BH3-only Bcl-2 family member Bim inhibits BCR stimulation-induced apoptosis and deletion of autoreactive B cells. J Exp Med. 2003;198(7):1119–26.

20. Alfredsson J, Puthalakath H, Martin H, Strasser A, and Nilsson G. Proapoptotic Bcl-2 family member Bim is involved in the control of mast cell survival and is induced together with Bcl-XL upon IgE-receptor activation. Cell Death Differ. 2005;12(2):136–44.

21. Kotzin JJ, Spencer SP, McCright SJ, Kumar DB, Collet MA, Mowel WK, Elliott EN, Uyar A, Makiya MA, Dunagin MC, et al. The long non-coding RNA Morrbid regulates Bim and short-lived myeloid cell lifespan. Nature. 2016;537(7619):239–43.

22. Hui CC, Yu A, Heroux D, Akhabir L, Sandford AJ, Neighbour H, and Denburg JA. Thymic stromal lymphopoietin (TSLP) secretion from human nasal epithelium is a function of TSLP genotype. Mucosal Immunol. 2015;8(5):993–9.

23. Williams BB, Tebbutt NC, Buchert M, Putoczki TL, Doggett K, Bao S, Johnstone CN, Masson F, Hollande F, Burgess AW, et al. Glycoprotein A33 deficiency: a new mouse model of impaired intestinal epithelial barrier function and inflammatory disease. Dis Model Mech. 2015;8(8):805–15.

24. Consortium GT. Human genomics. The Genotype-Tissue Expression (GTEx) pilot analysis: multitissue gene regulation in humans. Science. 2015;348(6235):648–60.

25. Grunewald TG, Pasedag SM, and Butt E. Cell Adhesion and Transcriptional Activity - Defining the Role of the Novel Protooncogene LPP. Transl Oncol. 2009;2(3):107–16.

26. Dent AL, Shaffer AL, Yu X, Allman D, and Staudt LM. Control of inflammation, cytokine expression, and germinal center formation by BCL-6. Science. 1997;276(5312):589–92.

27. Sehmi R, Wardlaw AJ, Cromwell O, Kurihara K, Waltmann P, and Kay AB. Interleukin-5 selectively enhances the chemotactic response of eosinophils obtained from normal but not eosinophilic subjects. Blood. 1992;79(11):2952–9.

28. Kim EH, Gasper DJ, Lee SH, Plisch EH, Svaren J, and Suresh M. Bach2 regulates homeostasis of Foxp3+ regulatory T cells and protects against fatal lung disease in mice. J Immunol. 2014;192(3):985–95.

29. Ho IC, Tai TS, and Pai SY. GATA3 and the T-cell lineage: essential functions before and after T-helper-2-cell differentiation. Nat Rev Immunol. 2009;9(2):125–35.

30. Schuster C, Gerold KD, Schober K, Probst L, Boerner K, Kim MJ, Ruckdeschel A, Serwold T, and Kissler S. The Autoimmunity-Associated Gene CLEC16A Modulates Thymic Epithelial Cell Autophagy and Alters T Cell Selection. Immunity. 2015;42(5):942–52.

31. Marine JC, Topham DJ, McKay C, Wang D, Parganas E, Stravopodis D, Yoshimura A, and Ihle JN. SOCS1 deficiency causes a lymphocyte-dependent perinatal lethality. Cell. 1999;98(5):609–16.

32. Davison LJ, Wallace C, Cooper JD, Cope NF, Wilson NK, Smyth DJ, Howson JM, Saleh N, Al-Jeffery A, Angus KL, et al. Long-range DNA looping and gene expression analyses identify DEXI as an autoimmune disease candidate gene. Hum Mol Genet. 2012;21(2):322–33.

33. Loh PR, Bhatia G, Gusev A, Finucane HK, Bulik-Sullivan BK, Pollack SJ, Schizophrenia Working Group of Psychiatric Genomics C, de Candia TR, Lee SH, Wray NR, et al. Contrasting genetic architectures of schizophrenia and other complex diseases using fast variance-components analysis. Nat Genet. 2015;47(12):1385–92.

34. Matzaraki V, Kumar V, Wijmenga C, and Zhernakova A. The MHC locus and genetic susceptibility to autoimmune and infectious diseases. Genome Biol. 2017;18(1):76.

35. Mohammad AJ, Hot A, Arndt F, Moosig F, Guerry MJ, Amudala N, Smith R, Sivasothy P, Guillevin L, Merkel PA, et al. Rituximab for the treatment of eosinophilic granulomatosis with polyangiitis (Churg-Strauss). Ann Rheum Dis. 2016;75(2):396–401.

36. Ference BA, Robinson JG, Brook RD, Catapano AL, Chapman MJ, Neff DR, Voros S, Giugliano RP, Davey Smith G, Fazio S, et al. Variation in PCSK9 and HMGCR and Risk of Cardiovascular Disease and Diabetes. N Engl J Med. 2016;375(22):2144–53.

37. Wechsler ME, Akuthota P, Jayne D, Khoury P, Klion A, Langford CA, Merkel PA, Moosig F, Specks U, Cid MC, et al. Mepolizumab or Placebo for Eosinophilic Granulomatosis with Polyangiitis. N Engl J Med. 2017;376(20):1921–32.

38. Delbridge AR, Grabow S, Strasser A, and Vaux DL. Thirty years of BCL-2: translating cell death discoveries into novel cancer therapies. Nat Rev Cancer. 2016;16(2):99–109.

39. Carrington EM, Vikstrom IB, Light A, Sutherland RM, Londrigan SL, Mason KD, Huang DC, Lew AM, and Tarlinton DM. BH3 mimetics antagonizing restricted prosurvival Bcl-2 proteins represent another class of selective immune modulatory drugs. Proc Natl Acad Sci U S A. 2010;107(24):10967–71.

40. Karlberg M, Ekoff M, Huang DC, Mustonen P, Harvima IT, and Nilsson G. The BH3-mimetic ABT-737 induces mast cell apoptosis in vitro and in vivo: potential for therapeutics. J Immunol. 2010;185(4):2555–62.

41. Ahmed CM, Larkin J, 3rd, and Johnson HM. SOCS1 Mimetics and Antagonists: A Complementary Approach to Positive and Negative Regulation of Immune Function. Front Immunol. 2015;6(183.

42. Sekine C, Sugihara T, Miyake S, Hirai H, Yoshida M, Miyasaka N, and Kohsaka H. Successful treatment of animal models of rheumatoid arthritis with small-molecule cyclin-dependent kinase inhibitors. J Immunol. 2008;180(3):1954–61.

43. Day N, Oakes S, Luben R, Khaw KT, Bingham S, Welch A, and Wareham N. EPIC-Norfolk: study design and characteristics of the cohort. European Prospective Investigation of Cancer. Br J Cancer. 1999;80 Suppl 1(95–103.

44. Chang CC, Chow CC, Tellier LC, Vattikuti S, Purcell SM, and Lee JJ. Second-generation PLINK: rising to the challenge of larger and richer datasets. Gigascience. 2015;4(7.

45. Delaneau O, Zagury JF, and Marchini J. Improved whole-chromosome phasing for disease and population genetic studies. Nat Methods. 2013;10(1):5–6.

46. Howie B, Fuchsberger C, Stephens M, Marchini J, and Abecasis GR. Fast and accurate genotype imputation in genome-wide association studies through pre-phasing. Nat Genet. 2012;44(8):955–9.

47. Howie BN, Donnelly P, and Marchini J. A flexible and accurate genotype imputation method for the next generation of genome-wide association studies. PLoS Genet. 2009;5(6):e1000529.

48. Devlin B, and Roeder K. Genomic control for association studies. Biometrics. 1999;55(4):997–1004.

49. Loh PR, Tucker G, Bulik-Sullivan BK, Vilhjalmsson BJ, Finucane HK, Salem RM, Chasman DI, Ridker PM, Neale BM, Berger B, et al. Efficient Bayesian mixed-model analysis increases association power in large cohorts. Nat Genet. 2015;47(3):284–90.

50. Wakefield J. Bayes factors for genome-wide association studies: comparison with P-values. Genet Epidemiol. 2009;33(1):79–86.

51. Wellcome Trust Case Control C, Maller JB, McVean G, Byrnes J, Vukcevic D, Palin K, Su Z, Howson JM, Auton A, Myers S, et al. Bayesian refinement of association signals for 14 loci in 3 common diseases. Nat Genet. 2012;44(12):1294–301.

52. Gabriel SB, Schaffner SF, Nguyen H, Moore JM, Roy J, Blumenstiel B, Higgins J, DeFelice M, Lochner A, Faggart M, et al. The structure of haplotype blocks in the human genome. Science. 2002;296(5576):2225–9.

53. Motyer AM, Vukcevic D, Dilthey A, Donnelly P, McVean G, and Leslie S. Practical Use of Methods for Imputation of HLA Alleles from SNP Genotype Data. http://biorxiv.org/content/early/2016/12/09/091009.

54. Bulik-Sullivan BK, Loh PR, Finucane HK, Ripke S, Yang J, Schizophrenia Working Group of the Psychiatric Genomics C, Patterson N, Daly MJ, Price AL, and Neale BM. LD Score regression distinguishes confounding from polygenicity in genome-wide association studies. Nat Genet. 2015;47(3):291–5.

55. Javierre BM, Burren OS, Wilder SP, Kreuzhuber R, Hill SM, Sewitz S, Cairns J, Wingett SW, Varnai C, Thiecke MJ, et al. Lineage-Specific Genome Architecture Links Enhancers and Non-coding Disease Variants to Target Gene Promoters. Cell. 2016;167(5):1369–84 e19.

56. Mifsud B, Tavares-Cadete F, Young AN, Sugar R, Schoenfelder S, Ferreira L, Wingett SW, Andrews S, Grey W, Ewels PA, et al. Mapping long-range promoter contacts in human cells with high-resolution capture Hi-C. Nat Genet. 2015;47(6):598–606.

57. Schofield EC, Carver T, Achuthan P, Freire-Pritchett P, Spivakov M, Todd JA, and Burren OS. CHiCP: a web-based tool for the integrative and interactive visualization of promoter capture Hi-C datasets. Bioinformatics. 2016;32(16):2511–3.

58. Staley JR, Blackshaw J, Kamat MA, Ellis S, Surendran P, Sun BB, Paul DS, Freitag D, Burgess S, Danesh J, et al. PhenoScanner: a database of human genotype-phenotype associations. Bioinformatics. 2016;32(20):3207–9.

59. Jones RB, Tervaert JW, Hauser T, Luqmani R, Morgan MD, Peh CA, Savage CO, Segelmark M, Tesar V, van Paassen P, et al. Rituximab versus cyclophosphamide in ANCA-associated renal vasculitis. N Engl J Med. 2010;363(3):211–20.

