## Supplementary material for "Genetically distinct clinical subsets, and associations with asthma and eosinophil abundance, within Eosinophilic Granulomatosis with Polyangiitis"

**Supplementary Appendix**

Additional authors and affiliations.
Supplementary Methods.
Supplementary Figure 1. Manhattan plots showing only directly genotyped SNPs.
Supplementary Figure 2. Correlation of estimated effect sizes between primary and replication cohorts.
Supplementary Figure 3. Enrichment of asthma and eosinophil-associated variants in EGPA.
Supplementary Figure 4. Genetic similarity between EGPA subsets and asthma.
Supplementary Figure 5. Contour and density plots of test statistics for quantifying pleiotropy.
Supplementary Figure 6. Genomic features and associations with other traits at non-MHC EGPA-associated loci.
Supplementary Figure 7. Locus zoom plots of loci associated with EGPA.
Supplementary Figure 8. Genetic effects on eosinophil count correlate with risk of EGPA.
Supplementary Figure 9. The *TSLP* promoter region variant rs1837253 has a greater effect size in EGPA than in asthma.
Supplementary Figure 10. QQ plots of genetic associations in EGPA according to increasing degrees of association in IBD.
Supplementary Figure 11. The MHC association with MPO +ve EGPA is localized to the Class II region.
Supplementary Figure 12. Amino acid positions in HLA-DRB1, HLA-DQA1 and HLA-DQB1 associated with susceptibility to EGPA.
Supplementary Figure 13. Forest plot of Mendelian randomization estimates for the causal effect of eosinophil count on EGPA.
Supplementary Figure 14. Principal components analysis (PCA) of genotype data.
Supplementary Figure 15. QQ plots for all EGPA cases vs controls.

Supplementary Table 1. Criteria for the diagnosis of EGPA from the 'Study to Investigate Mepolizumab in the Treatment of Eosinophilic Granulomatosis With Polyangiitis' (MIRRA[$]).

Supplementary Table 2. Breakdown of 542 cases and 6717 controls by country and center.

Supplementary Table 3. Ethics approval from each contributing centre.

Supplementary Table 4. Genetic association analysis using a linear mixed model (LMM-Bolt).

Supplementary Table 5. Replication cohort case demographics by country of origin.

Supplementary Table 6. Meta-analysis of genetic associations with EGPA in the primary and replication cohorts.

Supplementary Table 7. Direction of effect at EGPA variants on eosinophil count and asthma risk.

Supplementary Table 8: ANCA status according to country of recruitment.

Supplementary Table 9: associations of ANCA status with clinical features using logistic regression with adjustment for country of origin.

Supplementary Table 10. Meta-analysis of genetic associations with EGPA subsets stratified by ANCA status in the primary and replication cohorts.

Supplementary Table 11. Non-MHC EGPA-associated loci and other diseases.

Supplementary Table 12**.** Evidence to support biological plausibility of EGPA-associated variants.

Supplementary Table 13: Association of classical MHC alleles with MPO+ve EGPA.

Supplementary Table 14**.** Minor allele frequencies at HLA alleles associated with EGPA stratified by country.

Legend for Supplementary Data Item 1 (cross-referencing of EGPA-associated variants with variants in high LD in the NHGRI GWAS Catalog).

Legend for Supplementary Data Item 2 (cross-referencing of EGPA-associated variants with eQTLs).

Legend for Supplementary Data Item 3 (variants associated with traits in NHGRI GWAS Catalog that lie within +/- 1MB of the EGPA-associated variants).

Legend for Supplementary Data Item 4 (Mendelian randomisation estimates).

Legend for Supplementary Data Item 5. Full EGPA GWAS summary statistics.

Supplementary References.

**Additional authors and affiliations**

**The European Vasculitis Genetics Consortium**

Mohammed Akil[31], Jonathan Barratt[32], Neil Basu[33], Adam S. Butterworth[4], Ian Bruce[34,35], Michael Clarkson[36], Niall Conlon[37], Bhasker DasGupta[38], Timothy W. R. Doulton[39], Georgina Espígol-Frigolé[8], Oliver Flossmann[40], Armando Gabrielli[41], Jolanta Gasior[42], Gina Gregorini[43], Giuseppe Guida[44], José Hernández-Rodríguez[8], Zdenka Hruskova[27], Amy Hudson[18], Ann Knight[45], Peter Lanyon[46], Raashid Luqmani[47], Malgorzata Magliano[48], Angelo A. Manfredi[23], Christopher Marguerie[49], Federica Maritati[30], Chiara Marvisi[30], Neil J. McHugh[50], Eamonn Molloy[51], Allan Motyer[16], Chetan Mukhtyar[52], Leonid Padyukov[53], Alberto Pesci[54], Sergio Prieto-Gonzalez[8], Marc Ramentol-Sintas[55], Petra Reis[25], Dario Roccatello[56], Patrizia Rovere-Querini[23], Carlo Salvarani[57], Francesca Santarsia[58], Roser Solans-Laque[55], Nicole Soranzo[9, 59], Jo Taylor[60], Julie Wessels[61], & Jochen Zwerina[25].

[31]Sheffield Royal Hallamshire Hospital, Sheffield, S10 2JF, UK.

[32]Department of Infection, Immunity and Inflammation, University of Leicester, Leicester, LE1 9HN UK.

[33]Institute of Medical Sciences, Aberdeen, AB25 2ZD, UK.

[34]NIHR Manchester Musculoskeletal Biomedical Research Unit, Central Manchester University Hospitals NHS Foundation Trust, Manchester, UK.

[35]Arthritis Research UK Centre for Epidemiology, Centre for Musculoskeletal Research, The University of Manchester, Manchester Academic Health Science Centre; Manchester, UK.

[36]Cork University Hospital, Cork, Ireland.

[37]Department of Immunology, St James' Hospital Dublin, Dublin 8, Ireland.

[38]Southend University Hospital, Westcliff-on-Sea, SS0 0RY, UK.

[39]East Kent Hospitals University NHS Foundation Trust, Kent and Canterbury Hospital, Canterbury, CT1 3NG, UK.

[40]Royal Berkshire Hospital NHS Trust, Reading, RG1 5AN, UK.

[41]Department of Internal Medicine, Ospedali Riuniti-Università, Politecnica delle Marche, Ancona, Italy.

[42]University Hospital, Department of allergy and immunology, Krakow, Poland.

[43]Nephrology Unit, Spedali Civili, Brescia, Italy.

[44]Internal Medicine II, Immunology and Allergology Outpatient Clinic, Medical Science Department, ASL TO2 Birago di Vische Hospital, and the University of Torino, Turin, Italy.

[45]Department of Medical Sciences, Uppsala University, 751 85 Uppsala, Sweden.

[46]Nottingham University Hospitals NHS Trust, Nottingham, NG7 2UH, UK.

[47]Nuffield Orthopaedic Centre, Oxford, OX3 7LD, UK.

[48]Stoke Mandeville Hospital, Aylesbury, HP21 8AL, UK.

[49]South Warwickshire NHS Foundation Trust, UK.

[50]Royal National Hospital for Rheumatic Disease, Bath, BA1 1RL, UK.

[51]St Vincent's Hospital Dublin, Dublin, Ireland.

[52]Norfolk and Norwich University Hospital, Norwich NR4 7UY, UK.

[53]Department of Medicine, Karolinska Univesity Hospital, 171 76 Stockholm, Sweden.

[54]Pneumology Unit, University of Milano Bicocca, Milan, Italy.

[55]Research Unit in Systemic Autoimmune Diseases, Vall d'Hebron Research Institute, Hospital Vall d'Hebron, Barcelona, Spain.

[56]Department of Rare, Immunologic, Hematologic and Immunohematologic Diseases, Center of Research of Immunopathology and Rare Diseases, University of Torino, Italy.

[57]Rheumatology Unit, Arcispedale S. Maria Nuova, Reggio Emilia, Italy.

[58]Nephrology Unit, University Hospital of Parma, Parma, Italy.

[59]Department of Haematology, University of Cambridge, Cambridge Biomedical Campus, Cambridge CB2 0PT, UK.

[60]Dorset County Hospital, Dorchester, DT1 2JY, UK.

[61]Royal Stoke University Hospital, Stoke-on-Trent, ST4 6QG, UK.

**Supplementary Methods**

**Inclusion criteria**

There are no validated diagnostic criteria for EGPA. The 2012 Chapel Hill Consensus Conference (CHCC) described EGPA as a disease characterized by `eosinophil-rich and necrotizing granulomatous inflammation often involving the respiratory tract, and necrotizing vasculitis predominantly affecting small to medium vessels, and associated with asthma and eosinophilia'(1). However, as acknowledged in the CHCC publication itself, the product of the CHCC is a nomenclature system, and not diagnostic or classification criteria, and thus the CHCC definition is not suitable for diagnosis.

The emphasis on a histopathological definition of EGPA can be traced back to Churg and Strauss's original description of the syndrome, which was made largely on the basis of post-mortem studies of patients with untreated long-standing disease(2). In modern clinical practice, overt vasculitis is much harder to detect. In a series of 23 EGPA patients published by Reid *et al* in 1998, only 4 met the original histopathological criteria of Churg and Strauss(3). There are multiple reasons for this. Many patients who present with EGPA are already on chronic corticosteroid therapy for asthma control. Patients who present with organ- or life-threatening disease and the typical clinical, radiological, hematological and serological findings are treated empirically with high-dose corticosteroids, and so if a biopsy is taken it is usually after treatment has been instituted. Affected tissues may not be easily accessible for biopsy, and the tissue samples that are obtained are small, making vasculitis much harder to detect than on post-mortem studies.

Lanham *et al.* first identified the limitations of diagnostic criteria that focus narrowly on fulfilling the pathological features of necrotizing vasculitis, extravascular granulomata and tissue infiltration by eosinophils, since these pathological features often do not co-exist spatially or temporally, or indeed at all(4).  In recognition of the limitations of diagnostic criteria that required histopathological evidence of necrotizing vasculitis and granulomata, Lanham *et al* proposed diagnostic criteria(4) that required the presence of asthma, peripheral blood eosinophilia (peak count >$1.5 \times 10^9$/L) and systemic vasculitis involving two or more extra-pulmonary organs. The evidence of vasculitis could be clinical or radiological, and did not have to be confirmed histopathologically. These criteria are not widely used, as they fail to identify the many patients who do not have overt evidence of vasculitis.

The 1990 American College of Rheumatology (ACR) classification criteria were designed to classify patients with already documented vasculitis(5). The ACR criteria were derived through analysis of 20 EGPA patients and 787 patients with other forms of vasculitis, in which multivariate modelling was used to select a set of 6 features that most effectively discriminated EGPA from other forms of vasculitis when 4 or more were present. In this dataset, the presence of 4 or more of the 6 yielded a sensitivity of 85% and a specificity of 99.7%. However, it is important to note the ACR criteria were not developed for making a diagnosis in individual patients and have not been validated for this purpose(6). Indeed, The ACR criteria for other vasculitides developed concurrently using this dataset have been shown to perform poorly when used for diagnosis(7). In the series of 23 patients with a clinical diagnosis of EGPA reported by Reid et al, only 14 met the ACR criteria(3). Therefore the ACR criteria are unsuitable for most clinical studies.

More appropriate for use in clinical or genetic studies are the recently developed diagnostic criteria used in the Phase III clinical trial “Study to Investigate Mepolizumab in the Treatment of Eosinophilic Granulomatosis With Polyangiitis” (MIRRA: **Supplementary Table 1**)(8). These define EGPA diagnosis based on the history or presence of *both* asthma and eosinophilia (>1.0 \times 10$^{9}$/L and/or > 10% of leukocytes) *plus* at least two additional features of EGPA. Of note, the MIRRA criteria include a wider range of clinical features than the ACR criteria (e.g. cardiac involvement and glomerulonephritis), and also the results of ANCA testing, which was not widely available at the time the ACR criteria were developed.

**Supplementary Note: comparison of genetic similarity of ANCA negative EGPA and MPO positive EGPA to asthma**

We compared $Z$-scores from the MPO+ vs control analysis to $Z$-scores from asthma (which we denote $Z_a$) using the test statistic $X_{p,}$ defined below. To assess whether $X_{p,}$ was significant, we compared its observed value to distributions estimated under 3 sampling schema. We repeatedly resampled 161 samples with the same geographic distribution as the MPO+ cases, without replacement, under the following three schema:

A) from ANCA- cases, using all controls
B) from all EGPA cases, using all controls
C) from controls, using remaining controls as the control set.

For each draw $i$, we conducted a GWAS case/control analysis using the same methodology and covariates as for the MPO+/control analysis and calculated a value for $X_{p,}$, to generate sets of test statistics $X_A$, $X_B$, $X_C$ from draws under schemas A, B and C respectively.

We estimated a p-value for the deviation of $X_p$ from what would be expected under a null hypothesis $H_0^{hom}$ of non-HLA genetic homogeneity between MPO+ and ANCA- subtypes. We assumed a sampling distribution of $X_p$ under $H_0^{hom}$ as

$$X_p | H_0^{hom} \sim N\big(mean(X_A), var(X_B)\big)$$

Under $H_0^{hom}$, by assumption, the mean of $X_A$ is equal to the expected value of $X_p$. The variance of $X_A$ underestimates the sampling variance of $X_p$, since there are fewer ANCA- cases than EGPA cases in total, so we estimated the sampling variance of $X_p$ with the mean of $X_B$. We note that we take the sampling variance of $X_p$ to correspond to a sampling schema of 'select 161 cases from the 542 available EGPA cases' rather than 'sample 161 cases from the EGPA population'. Since $X_p$ is an average across multiple SNPs, we assumed a normal sampling distribution for $X_p$ under CLT.

**Results**

Using this schema, we obtained a p-value of 1.7 x $10^{-3}$ against $H_0^{hom}$, indicating reasonable evidence for differential genetic basis between MPO+ and ANCA- EGPA outside the MHC region. $X_p$ was typical of the values $X_C$ (quantile 0.46), so we were unable to determine if MPO+ EGPA had any heritability outside the MHC region on the basis of this analysis. Densities of $X_A$, $X_B$, and $X_C$ compared to $X_p$ are shown in **Supplementary Figure 4**. If the MHC region was included, the MPO+ subtype was more similar to asthma than the ANCA- subtype (p=1.05 x $10^{-6}$).

We show that EGPA and asthma share associated variants (**Table 2**, **Supplementary Table 7**). Therefore the question arises as to whether the SNPs driving the difference in genetic architecture between ANCA- and MPO+ EGPA are or are not those associated with asthma. Expressed another way, are the two EGPA subsets genetically distinct because one subset more closely resembles asthma, or because they are simply intrinsically different, independent of any relationship with asthma? To address this, we chose a test statistic so as to be maximally sensitive to joint association with asthma and the EGPA subtype (see below), but that retains power to detect different genetic architectures between ANCA- and MPO+ EGPA at non-asthma associated variants. In other words, the value of the test statistic generally reflects 'similarity to asthma' but is also responsive to 'greater overall heritability'.

If the different genetic architectures between ANCA- and MPO+ EGPA were primarily due to different effect sizes at non-asthma associated SNPs we would expect that the difference would be largely retained if we removed any dependence between Z scores for asthma and Z scores for the EGPA subtype. To check this, we reproduced test statistics $X_A$, $X_p$ in the same with the $Z_a$ scores randomly shuffled to give test statistics $X_A'$, $X_p'$.

This removed any dependence between the sets of Z scores, but retained the marginal distributions of Z scores for asthma and for the EGPA subtypes. We found that both $X_A'$ and $X_p'$ were indistinguishable from $X_C$ (quantile of mean($X_A'$) in $X_C$ = 0.2, quantile of mean($X_p'$) in $X_C$ = 0.3), by contrast to $X_A$ and $X_p$ which were significantly different from $X_C$. This indicated that the observed difference in genetic architecture between MPO+ and ANCA-EGPA was generally at variants which were also associated with asthma, and that the two subtypes differ in their genetic similarity to asthma.

Graphically, a case in which the EGPA subtype showed no shared association with asthma would appear on **Supplementary Figure 5** as a set of points displaced significantly from the origin along the x-axis, but not the y-axis (variants associated with the EGPA subtype, but not with asthma), and a separate set of points displaced along the y-axis but not the x-axis (variants associated with asthma but not with the EGPA subtype). By contrast, SNPs associated with both asthma and the EGPA subtype would be displaced from the origin in both the x- and the y-axes simultaneously.

**Choice of statistic for pleiotropy**

To characterise the degree of pleiotropy between two phenotypes characterised by sets of Z scores, we sought a test statistic to detect concurrently high Z scores in both phenotypes. We considered two metrics for this; for Z-scores $Z_a(i)$ for asthma and $Z_t(i)$ for the trait $t$ under investigation at SNP $i$ of $n$ SNPs in total, these were

$$X_{1,\alpha}(Z_a, Z_t) = \left(\frac{\sum_i |Z_a(i) Z_t(i)|^\alpha}{n}\right)^{\frac{1}{\alpha}}$$

$$X_{2,\alpha}(Z_a, Z_t) = \left(\frac{\sum_i \left(\sqrt{Z_a(i)^4 + 1} + \sqrt{Z_t(i)^4 + 1} - \sqrt{(Z_a(i)^2 - Z_t(i)^2)^2 + 1} - 1\right)^\alpha}{n}\right)^{\frac{1}{\alpha}}$$

with $\alpha$ in {1,2}. Contours of the two test statistics are shown in **Supplementary Figure 5**.

The form of test statistic 2 was chosen so that SNPs with simultaneously high $|Z_a|$ and $|Z_t|$ values would contribute to the statistic, but there would be minimal contribution from SNPs for which the point $(Z_a, Z_t)$ was near the *x* or *y* axis, even if one of $Z_a$ , $Z_t$ were very large.

We determined the statistic to use in the above analysis by determining which statistic best separated values $X_B$ from $X_C$ (defined as in the previous section), assessing separation by the value of a t-statistic score between the two sets of values. The best-performing test statistic was $X_2$ with $\alpha$=2.

**Discussion**

In this analysis, we showed that ANCA-negative EGPA individuals have systematic genetic differences to MPO+ EGPA individuals outside of the MHC region, by establishing that the former showed greater pleiotropy with asthma. The phenotypic similarity of EGPA with asthma more generally indicates that these genetic differences are likely to correspond to clinically important pathophysiological processes. Given that our sampling maintained the geographic distribution of cases, this finding is unlikely to be confounded by geography.

An obvious metric for assessment of pleiotropy between asthma and the EGPA subtype of interest is genetic correlation ($r_g$). However, estimation of $r_g$ is complicated and estimates have prohibitively large variance when made using the small number of cases in this study. The aim of the analysis described above was simply to indicate genetic differences between the two EGPA subtypes, rather than to estimate genetic correlation. While the metric we used was somewhat simpler, it was difficult to compare its distribution across different study sizes, which necessitated the downsampling of ANCA-negative EGPA cases.

### Supplementary Figures

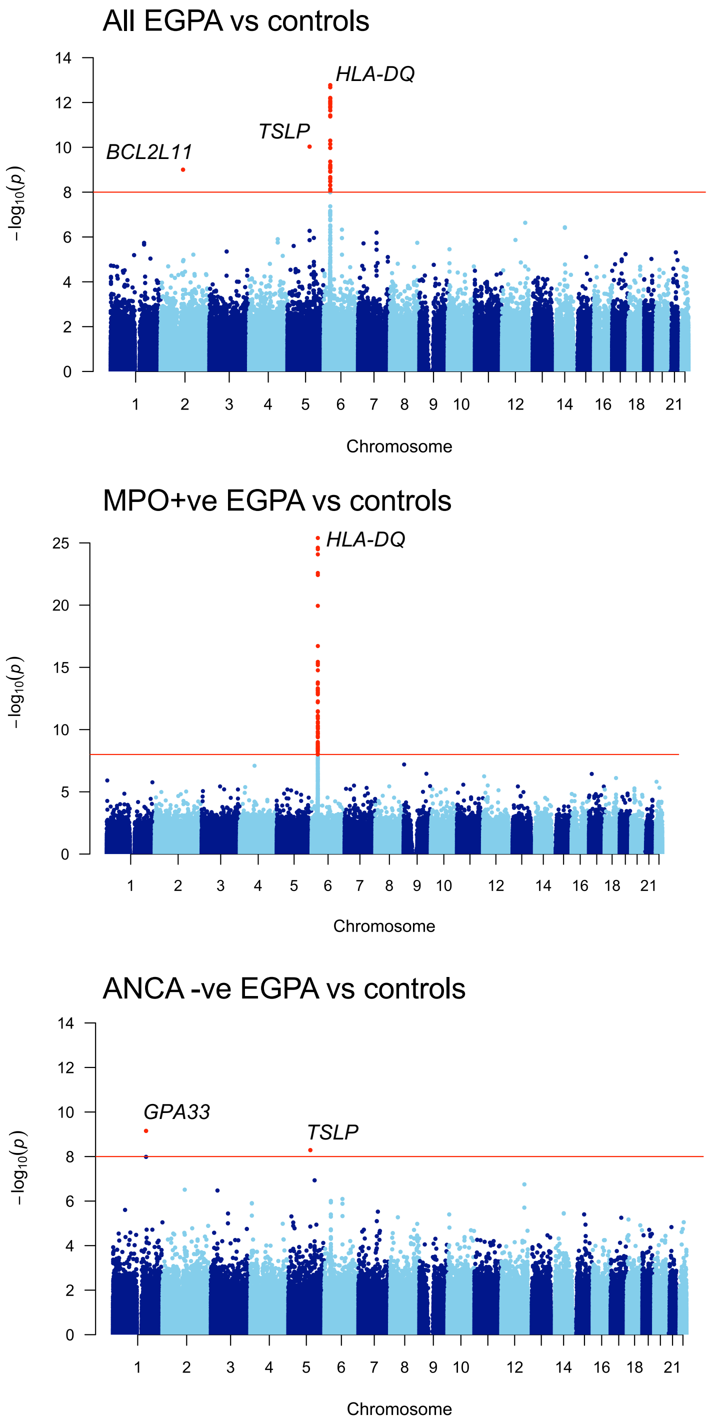

**Supplementary Figure 1. Manhattan plots showing only directly genotyped SNPs.** The horizontal red line indicates genome-wide significance (P $1x10^{-8}$, adjusted for genomic inflation). Significant SNPs are coloured in red.

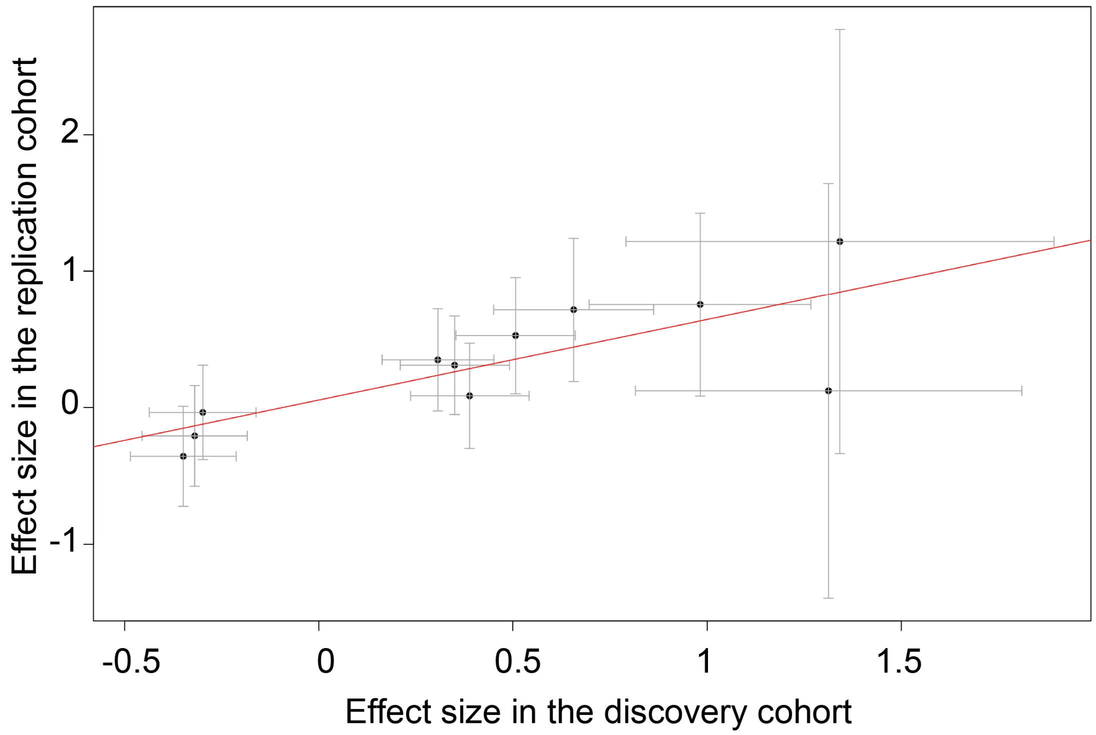

**Supplementary Figure 2. Correlation of estimated effect sizes between primary and replication cohorts**. Log (estimated ORs) (+/- 95% confidence intervals) are shown. Pearson r = 0.78, p = 0.005.

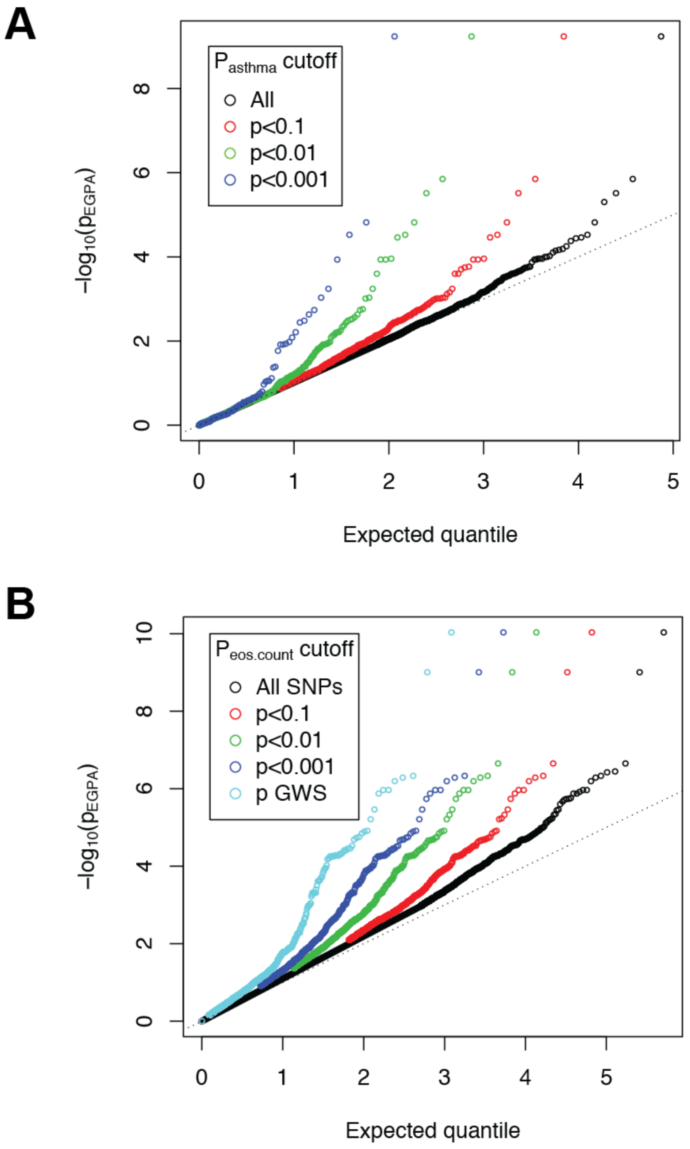

**Supplementary Figure 3. Enrichment of asthma and eosinophil-associated variants in EGPA.** QQ plots of observed –log10 p-values for the association of genotype with EGPA versus the expected –log10 p-values under the null hypothesis of no association, conditional on varying degrees of association with **A**) asthma and **B**) eosinophil count. The MHC region has been excluded. Each circle represents a SNP. The coloured circles indicate sets of SNPs with increasing degrees of association with asthma (panel A) or eosinophil count (panel B). Their QQ-plots demonstrate progressive departure from the line y=x (dotted), indicating shared genetic architecture between EGPA and these traits.

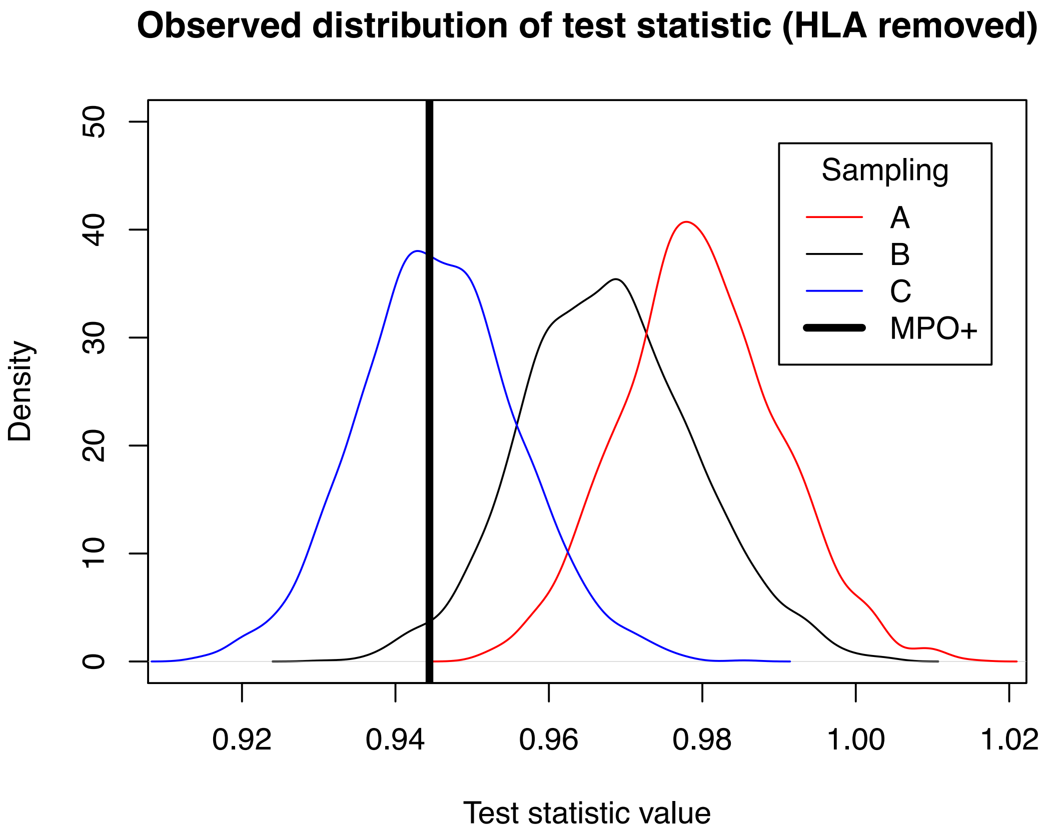

**Supplementary Figure 4. Genetic similarity between EGPA subsets and asthma**. The distributions of $X_A$, $X_B$, and $X_C$ under the sampling schemas described in the Supplementary Note are shown (MHC region removed). Greater right-displacement indicates greater genetic similarity with asthma. Samples from ANCA negative cases (A) show greater genetic similarity with asthma than do MPO+ cases (vertical line) or samples from all EGPA cases (B). MPO+ cases are indistinguishable from controls (C) on the basis of this data.

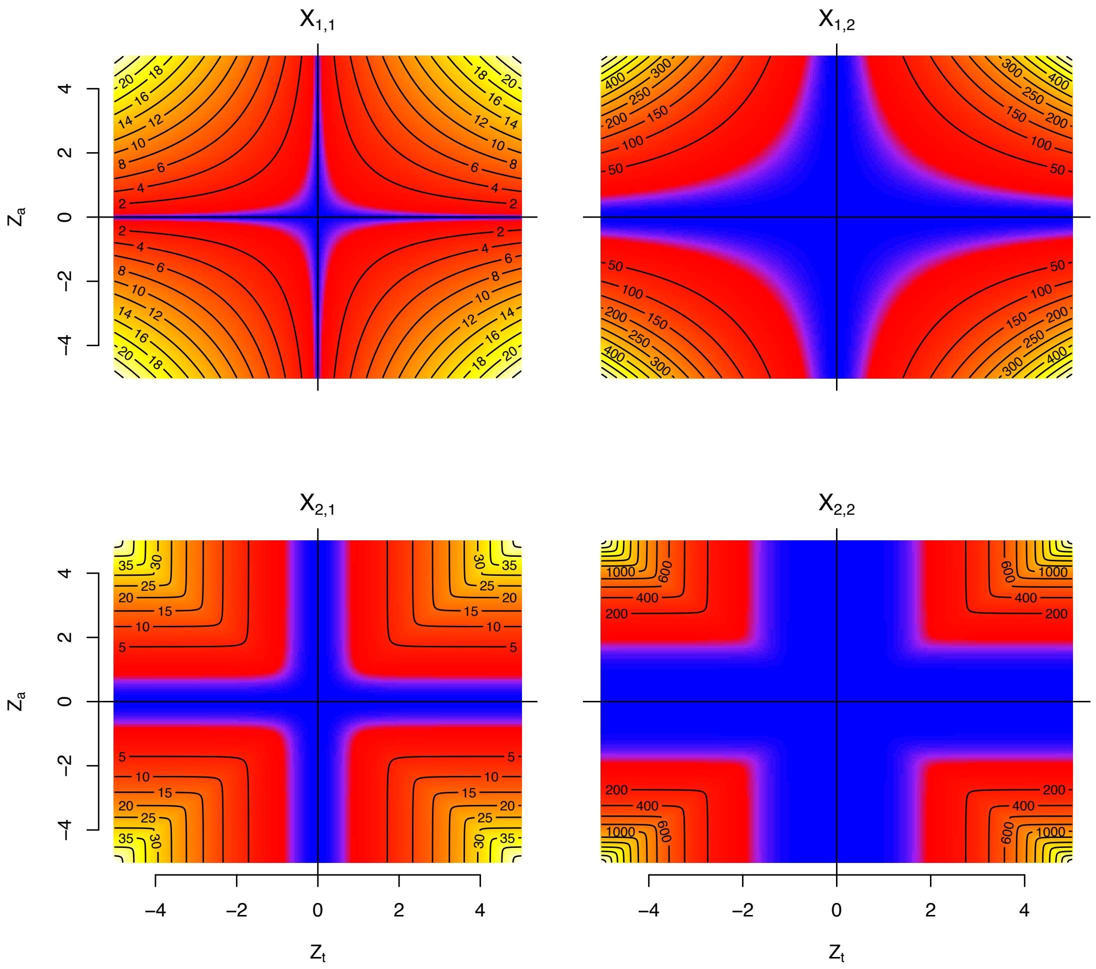

**Supplementary Figure 5**. **Contour and density plots of test statistics for quantifying pleiotropy.**

Blue colours correspond to values near zero, yellow to large values. The contribution to $X_1$ of a SNP with $(Z_a, Z_t)=(1,x)$ becomes arbitrarily large as $x\rightarrow\infty$, but the contribution to $X_2$ is bounded with increasing *x.*

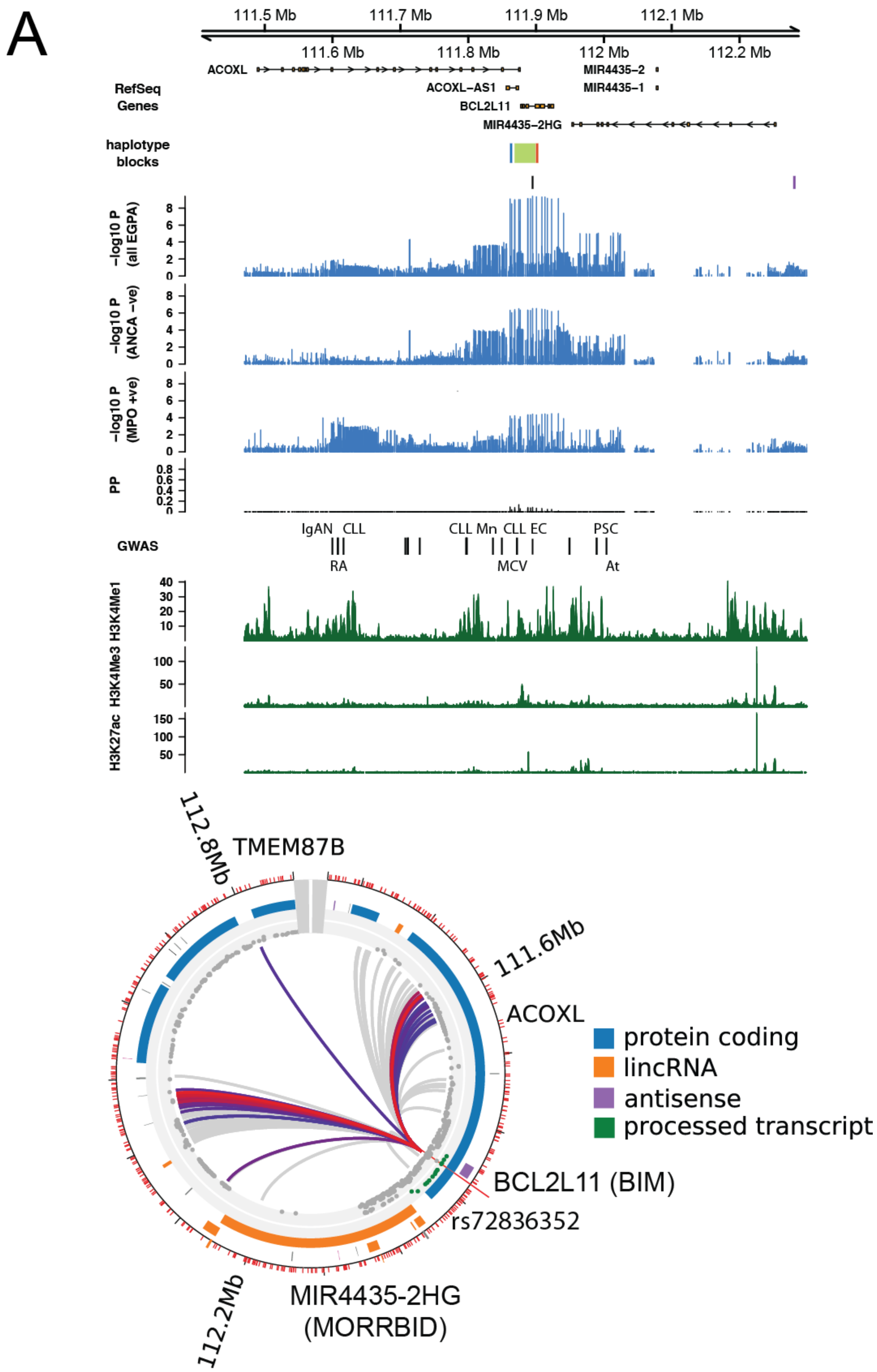
A
111.5 Mb
111.7 Mb
111.9 Mb
112.1 Mb
111.6 Mb
111.8 Mb
112 Mb
112.2 Mb
RefSeq Genes
ACOXL
ACOXL−AS1
BCL2L11
MIR4435−2HG
MIR4435−2
MIR4435−1
haplotype blocks
−log10 P (all EGPA)
−log10 P (ANCA −ve)
−log10 P (MPO +ve)
PP
GWAS
IgAN CLL
RA
CLL Mn CLL EC
MCV
PSC
At
H3K4Me1
H3K4Me3
H3K27ac
112.8Mb
TMEM87B
111.6Mb
ACOXL
BCL2L11 (BIM)
rs72836352
112.2Mb
MIR4435-2HG (MORRBID)
protein coding
lincRNA
antisense
processed transcript

B

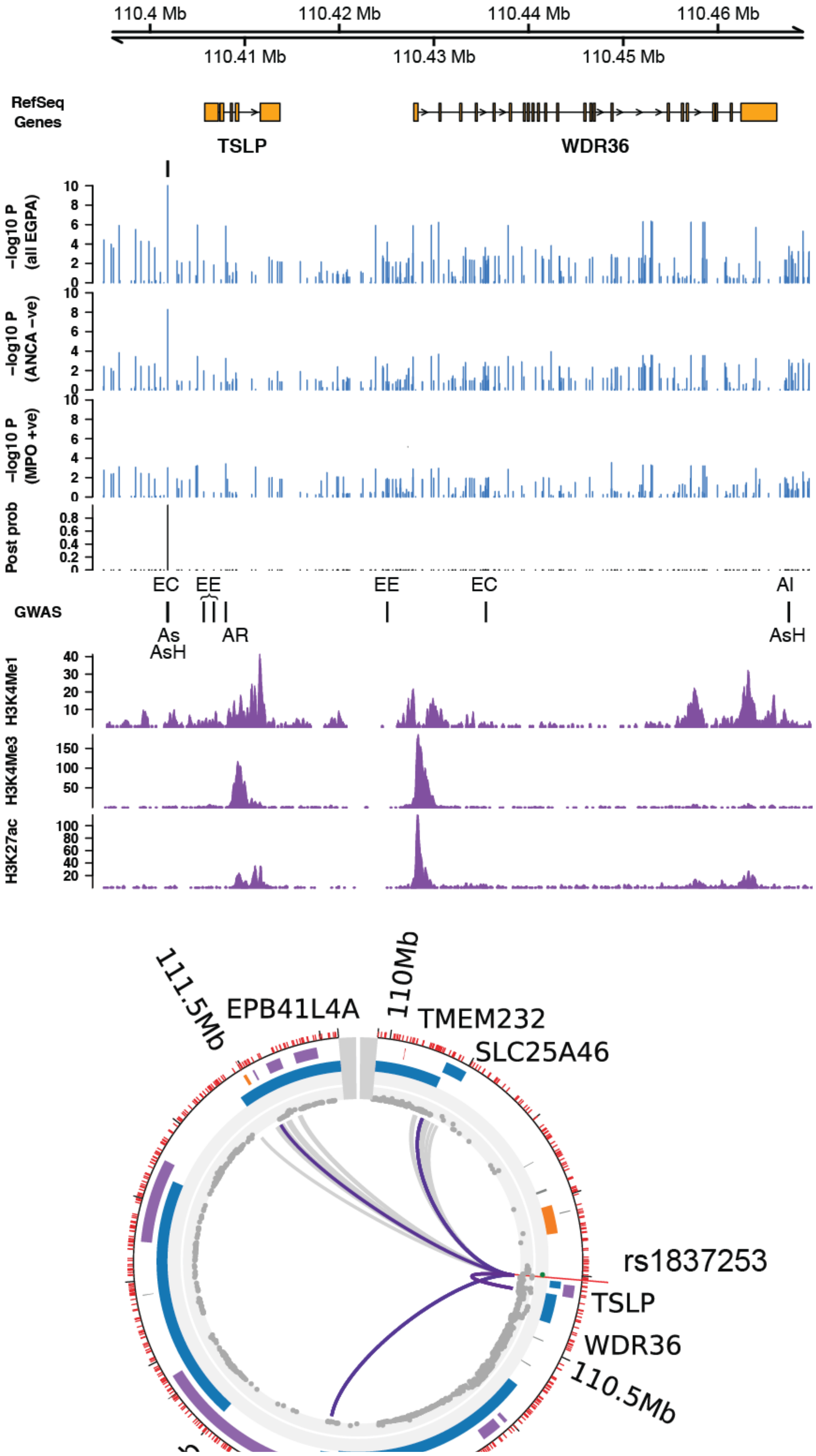

110.4 Mb
110.42 Mb
110.44 Mb
110.46 Mb
110.41 Mb
110.43 Mb
110.45 Mb
RefSeq Genes
TSLP
WDR36
−log10 P (all EGPA)
−log10 P (ANCA −ve)
−log10 P (MPO +ve)
Post prob
GWAS
EC
EE
EE
EC
AI
As
AsH
AR
AsH
H3K4Me1
H3K4Me3
H3K27ac
111.5Mb
110Mb
EPB41L4A
TMEM232
SLC25A46
rs1837253
TSLP
WDR36
110.5Mb

C

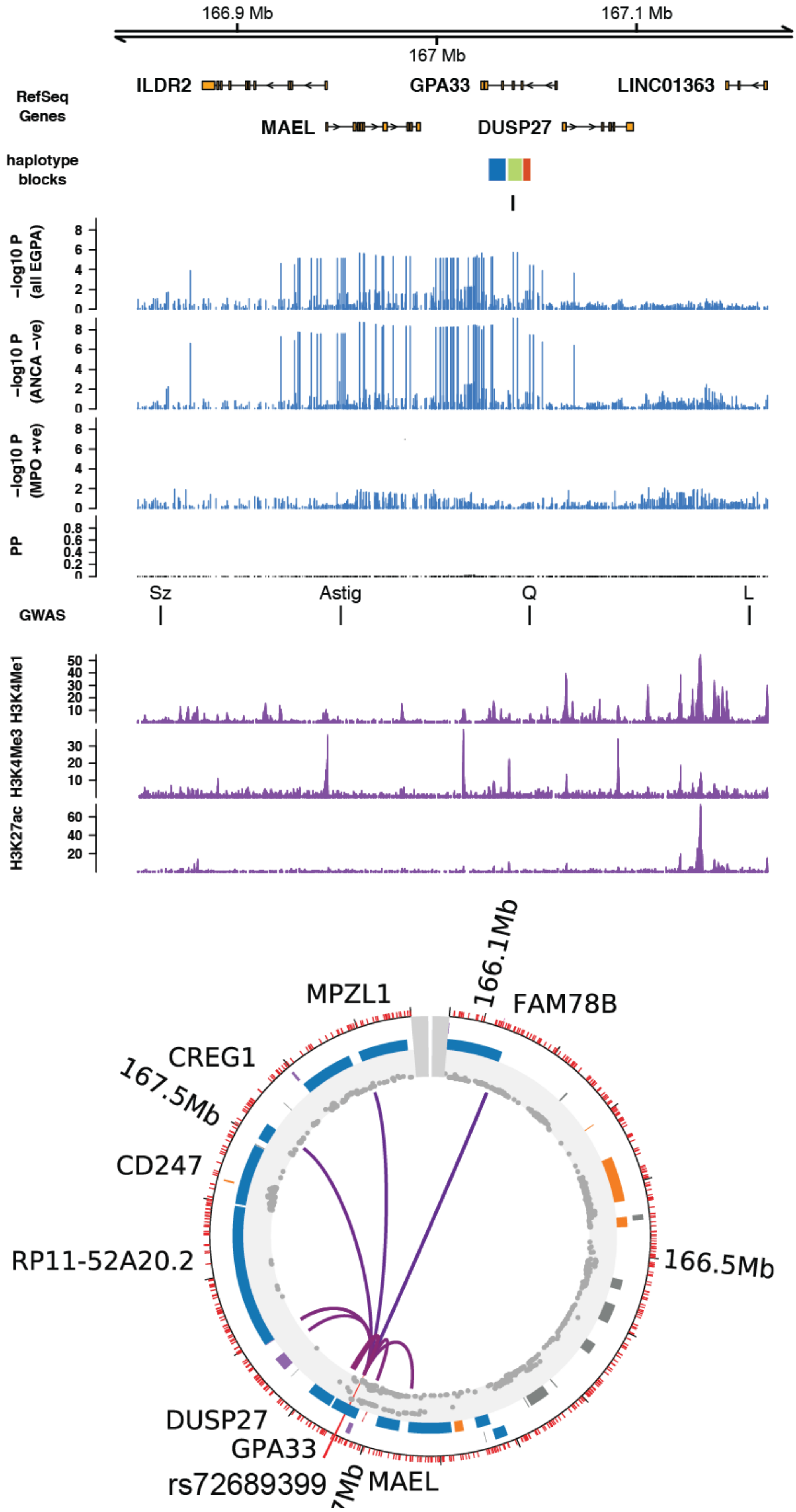
166.9 Mb
167 Mb
167.1 Mb
RefSeq Genes
ILDR2
GPA33
LINC01363
MAEL
DUSP27
haplotype blocks
−log10 P (all EGPA)
−log10 P (ANCA −ve)
−log10 P (MPO +ve)
PP
GWAS
Sz
Astig
Q
L
H3K4Me1
H3K4Me3
H3K27ac
MPZL1
166.1Mb
FAM78B
CREG1
167.5Mb
CD247
166.5Mb
RP11-52A20.2
DUSP27
GPA33
rs72689399
MAEL

D

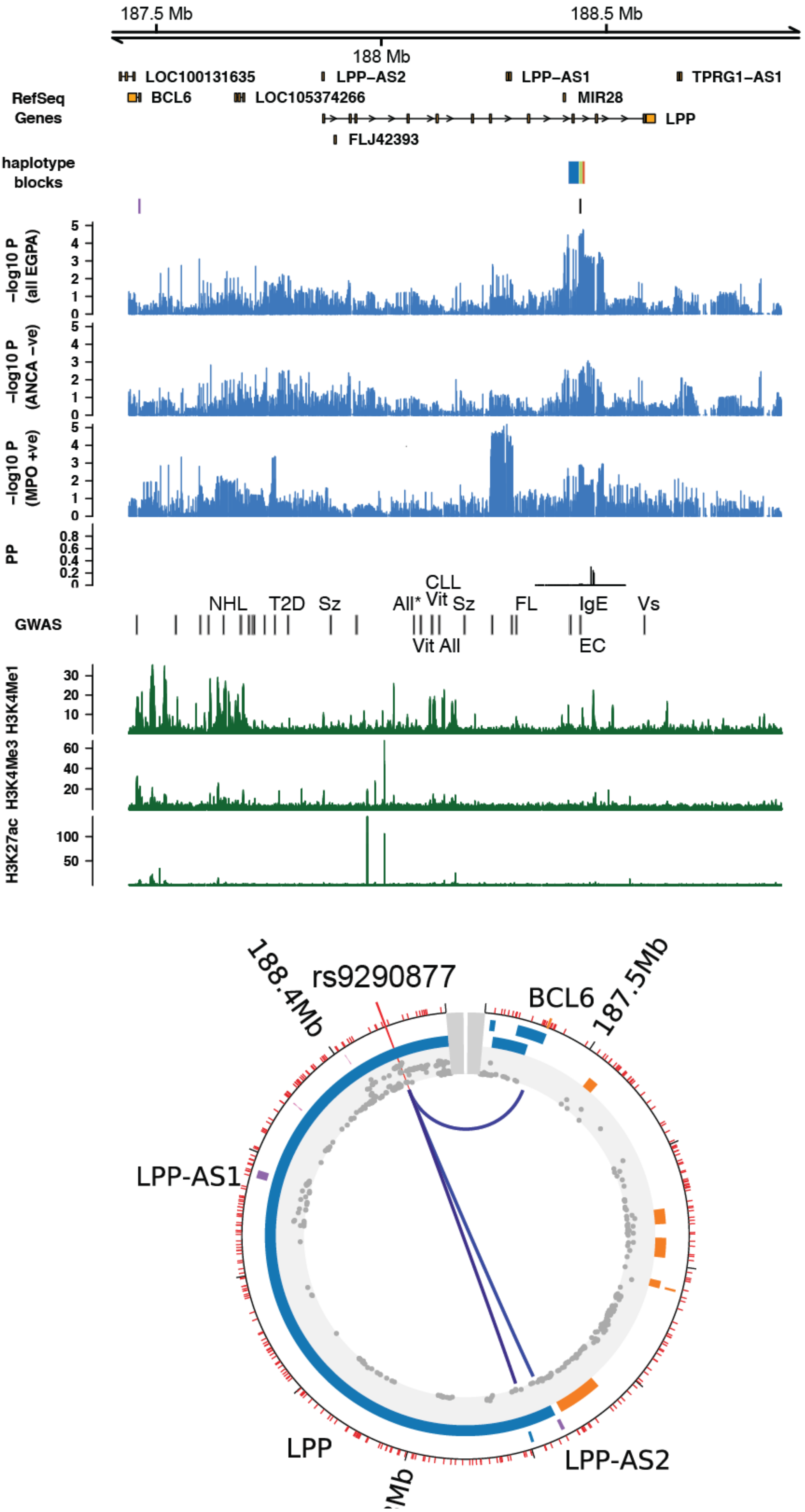

187.5 Mb
188.5 Mb
188 Mb
LOC100131635
LPP-AS2
LPP-AS1
TPRG1-AS1
RefSeq Genes
BCL6
LOC105374266
MIR28
LPP
FLJ42393
haplotype blocks
-log10 P (all EGPA)
-log10 P (ANCA -ve)
-log10 P (MPO +ve)
PP
CLL
NHL
T2D
Sz
All*
Vit
Sz
FL
IgE
Vs
GWAS
Vit
All
EC
H3K4Me1
H3K4Me3
H3K27ac
188.4Mb
rs9290877
BCL6
187.5Mb
LPP-AS1
LPP
LPP-AS2

E

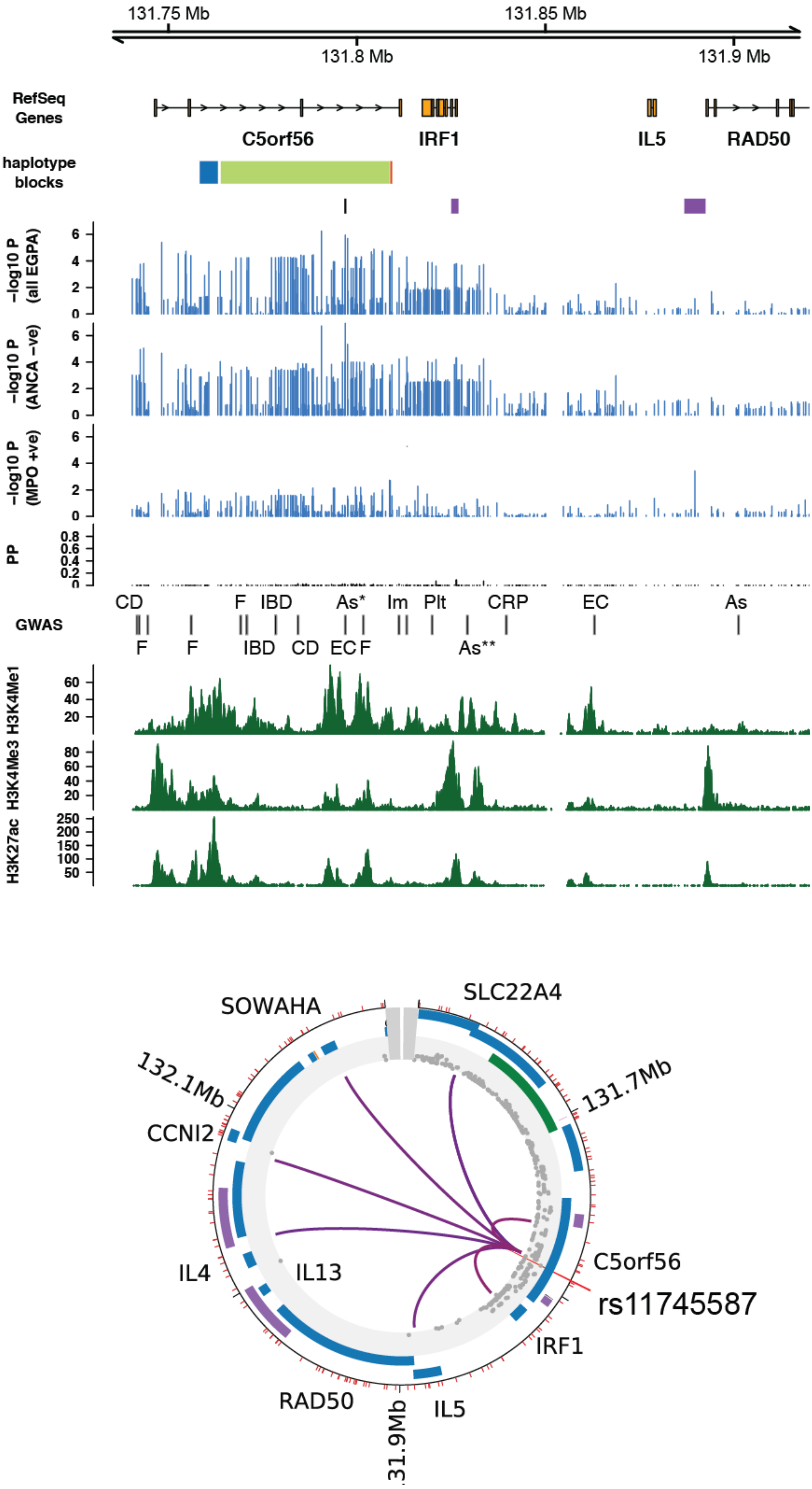

131.75 Mb
131.85 Mb
131.8 Mb
131.9 Mb
RefSeq Genes
C5orf56
IRF1
IL5
RAD50
haplotype blocks
−log10 P (all EGPA)
−log10 P (ANCA −ve)
−log10 P (MPO +ve)
PP
CD
F
IBD
As*
Im
Plt
CRP
EC
As
GWAS
F
F
IBD
CD
EC
F
As**
H3K4Me1
H3K4Me3
H3K27ac
SOWAHA
SLC22A4
132.1Mb
131.7Mb
CCNI2
IL4
IL13
C5orf56
rs11745587
IRF1
RAD50
IL5
131.9Mb

F

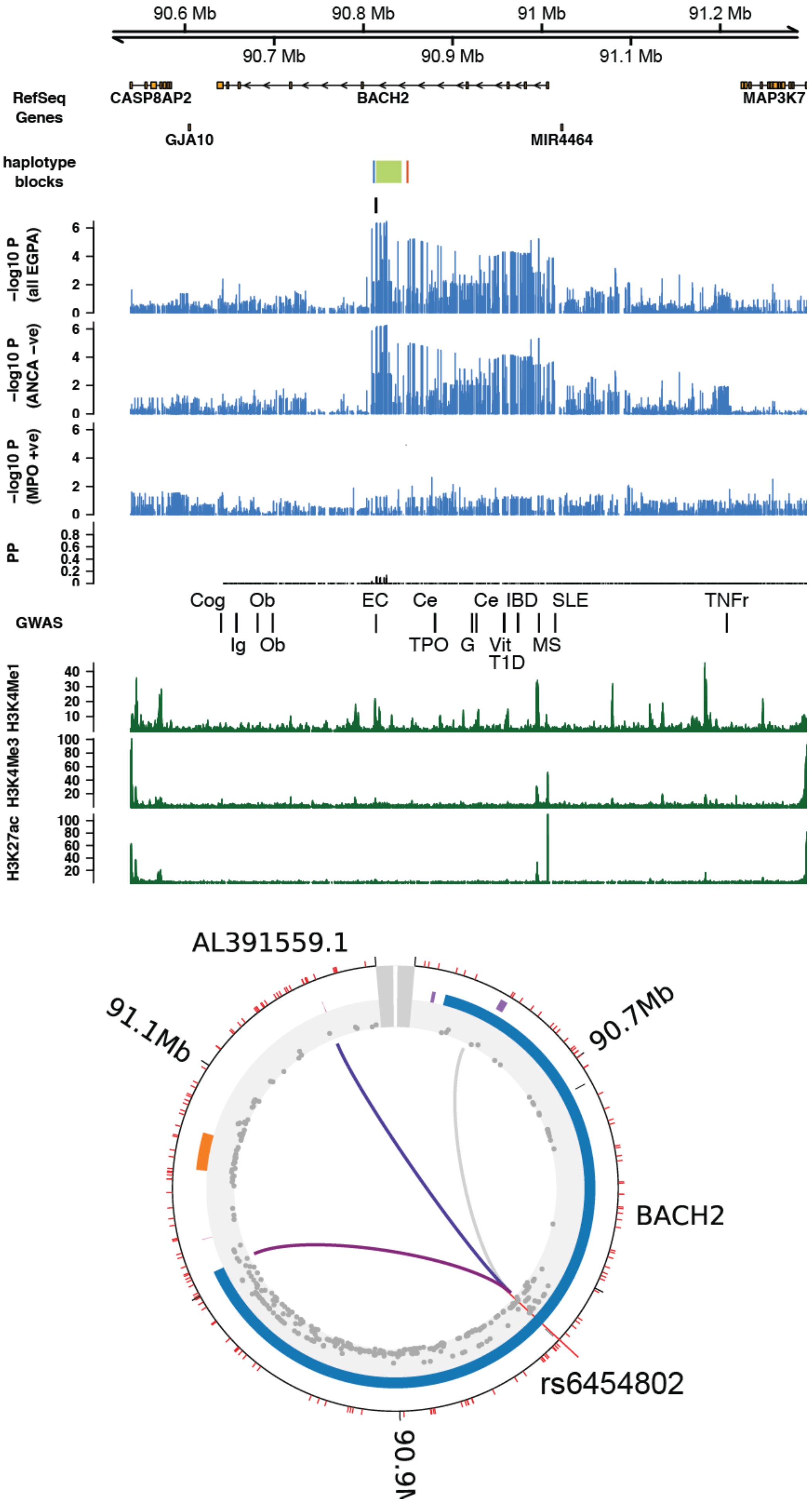

90.6 Mb
90.8 Mb
91 Mb
91.2 Mb
90.7 Mb
90.9 Mb
91.1 Mb
RefSeq Genes
CASP8AP2
BACH2
MAP3K7
GJA10
MIR4464
haplotype blocks
−log10 P (all EGPA)
−log10 P (ANCA −ve)
−log10 P (MPO +ve)
PP
GWAS
Cog
Ob
EC
Ce
Ce
IBD
SLE
TNFr
Ig
Ob
TPO
G
Vit
MS
T1D
H3K4Me1
H3K4Me3
H3K27ac
AL391559.1
91.1Mb
90.7Mb
BACH2
rs6454802
90.9Mb

G

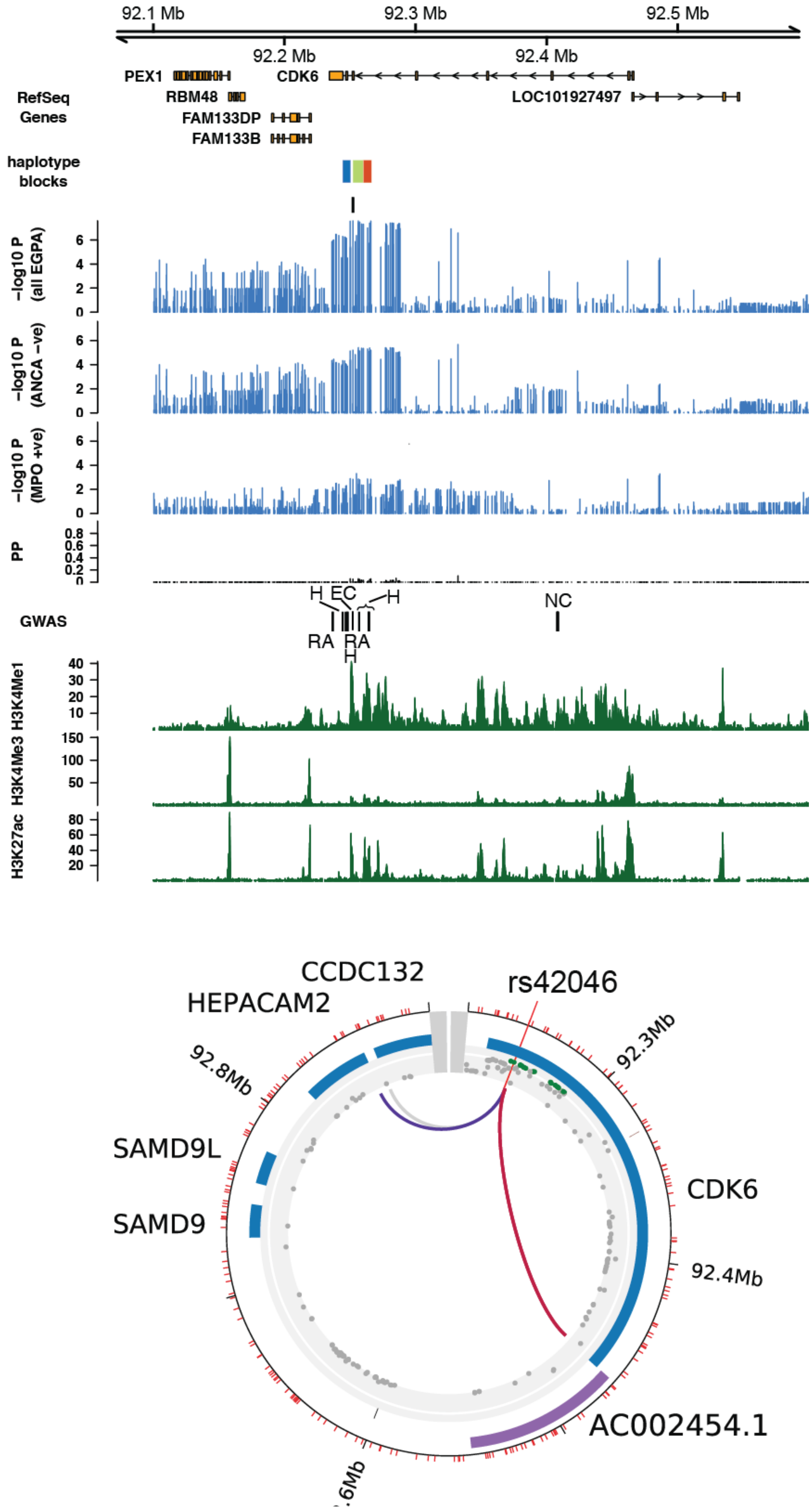

92.1 Mb
92.2 Mb
92.3 Mb
92.4 Mb
92.5 Mb
RefSeq Genes
PEX1
RBM48
FAM133DP
FAM133B
CDK6
LOC101927497
haplotype blocks
−log10 P (all EGPA)
−log10 P (ANCA −ve)
−log10 P (MPO +ve)
PP
GWAS
H EC
H
RA RA
H
NC
H3K4Me1
H3K4Me3
H3K27ac
CCDC132
HEPACAM2
rs42046
92.3Mb
92.8Mb
SAMD9L
SAMD9
CDK6
92.4Mb
AC002454.1

H

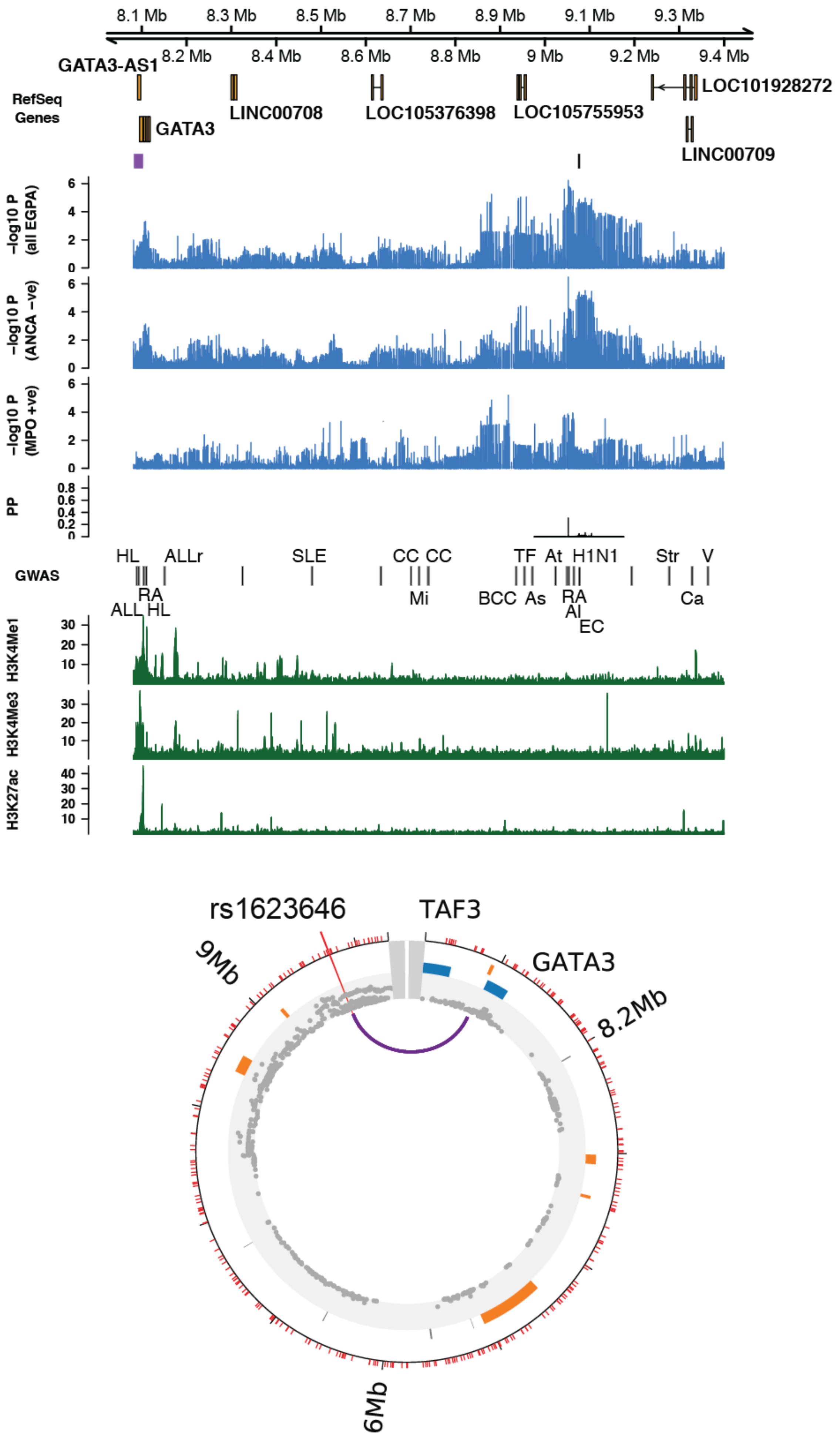

8.1 Mb
8.3 Mb
8.5 Mb
8.7 Mb
8.9 Mb
9.1 Mb
9.3 Mb
8.2 Mb
8.4 Mb
8.6 Mb
8.8 Mb
9 Mb
9.2 Mb
9.4 Mb
RefSeq Genes
GATA3-AS1
GATA3
LINC00708
LOC105376398
LOC105755953
LOC101928272
LINC00709
−log10 P (all EGPA)
−log10 P (ANCA −ve)
−log10 P (MPO +ve)
PP
GWAS
HL
ALLr
SLE
CC
CC
TF
At
H1N1
Str
V
RA
ALL
HL
Mi
BCC
As
RA
AI
EC
Ca
H3K4Me1
H3K4Me3
H3K27ac
rs1623646
TAF3
GATA3
9Mb
8.2Mb
6Mb

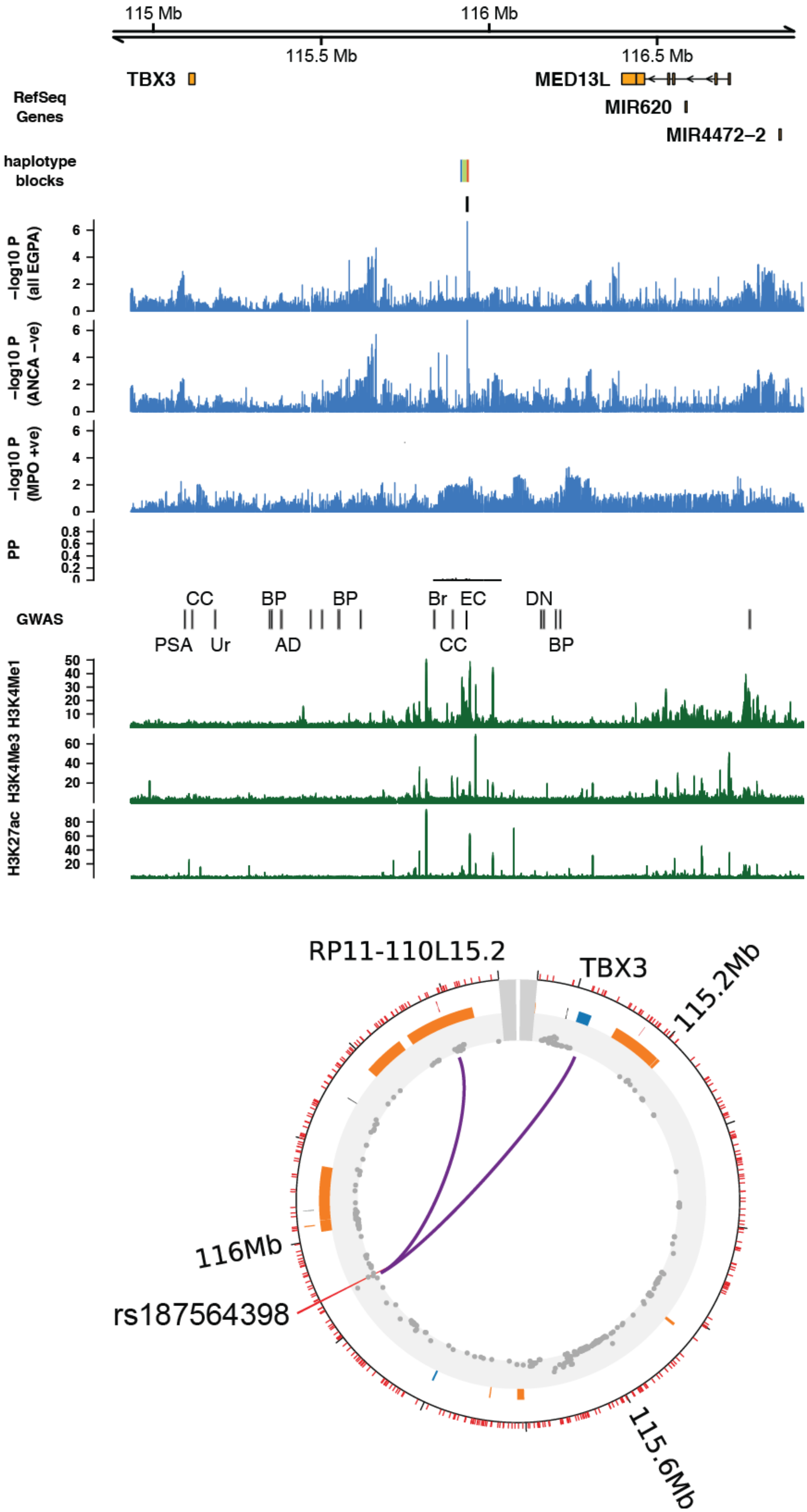

115 Mb
116 Mb
115.5 Mb
116.5 Mb
TBX3
MED13L
MIR620
MIR4472–2
RefSeq Genes
haplotype blocks
−log10 P (all EGPA)
−log10 P (ANCA −ve)
−log10 P (MPO +ve)
PP
GWAS
CC
BP
BP
Br
EC
DN
PSA
Ur
AD
CC
BP
H3K4Me1
H3K4Me3
H3K27ac
RP11-110L15.2
TBX3
115.2Mb
116Mb
rs187564398
115.6Mb

J

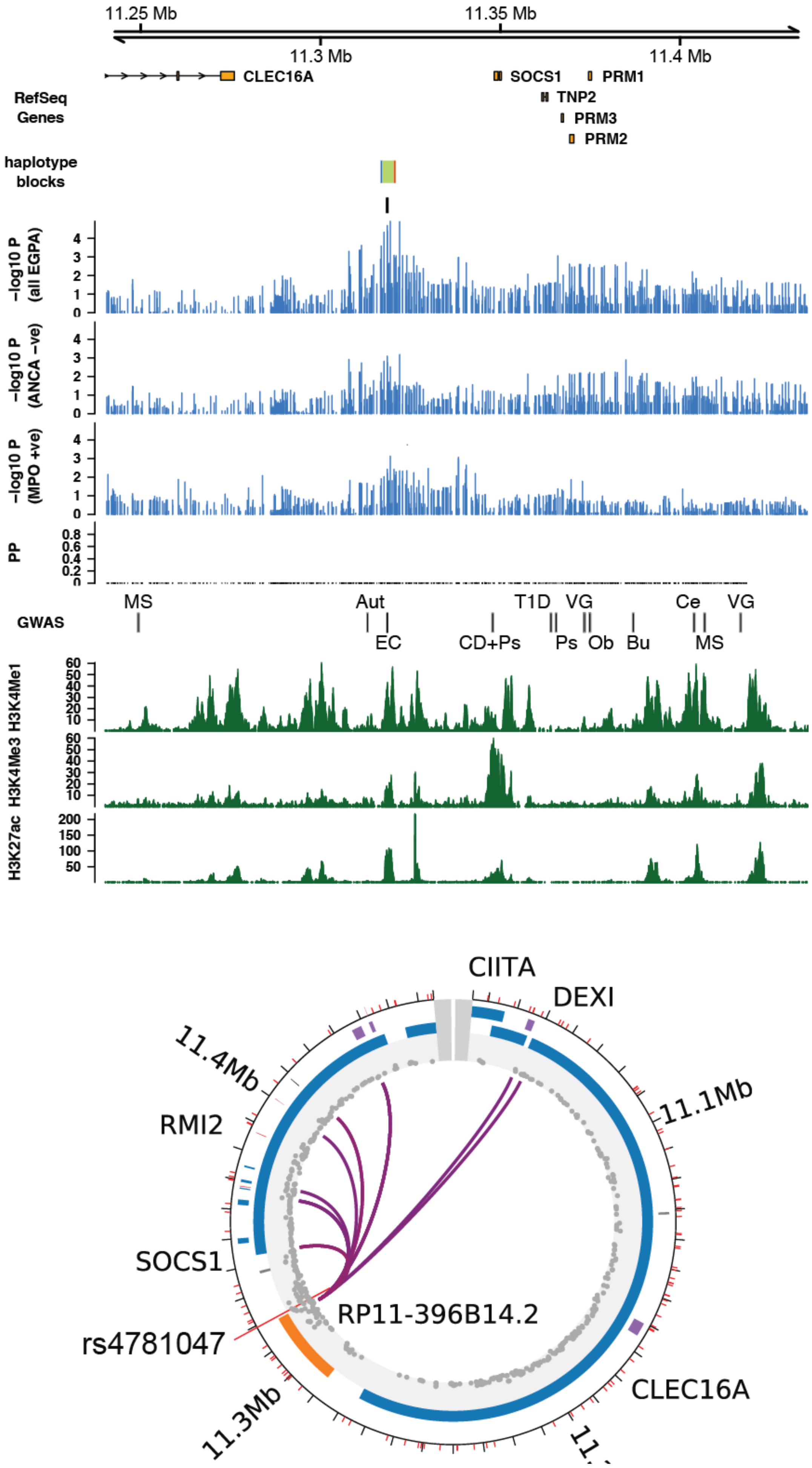

11.25 Mb
11.35 Mb
11.3 Mb
11.4 Mb
CLEC16A
SOCS1
PRM1
TNP2
PRM3
PRM2
RefSeq Genes
haplotype blocks
−log10 P (all EGPA)
−log10 P (ANCA −ve)
−log10 P (MPO +ve)
PP
MS
Aut
T1D
VG
Ce
VG
GWAS
EC
CD+Ps
Ps
Ob
Bu
MS
H3K4Me1
H3K4Me3
H3K27ac
CIITA
DEXI
11.4Mb
11.1Mb
RMI2
SOCS1
RP11-396B14.2
rs4781047
CLEC16A
11.3Mb

**Supplementary Figure 6. Genomic features and associations with other traits at non-MHC EGPA-associated loci.**

Upper panels from the top

- Genomic positions are from the hg19 genome build.
- The position of the EGPA-associated SNP reported in Table 2 is indicated by a vertical black bar. Vertical purple bars indicate the site(s) of DNA-DNA interaction with the EGPA-associated SNP identified by CHIC-P. Where the SNP lies in a haplotype block, the block is shown in green, and the proximal and distal blocks are shown in blue and red, respectively.
- Associations with EGPA as a whole, and with MPO+ and ANCA- subsets, are shown by vertical blue lines.
- PP= posterior probability from fine-mapping analysis.
- 'GWAS' panel indicates trait-associated SNPs from the NHGRI GWAS Catalog (those with p-values < $1 \times 10^{-5}$) and SNPs associated with eosinophil count in the study by Astle *et al*.(9) (see also Supplementary Table 5).
- Histone marks from ENCODE are shown in green (GM12878 lymphoblastoid cell line) or purple (NHEK epithelial cell line). H3K4me1 = associated with enhancer function and with active transcription. H3K4me3 = marker of promoters. H3K27ac = marker of active enhancers.

Lower panels: "circos plots" showing genomic segments displayed in a circular format. Adjacent parallel gray blocks indicate the proximal and distal ends of the segment. The position of the EGPA-associated SNP reported in Table 2 is indicated by a red line and rs number. Points on the inner circle indicate –log10 p-values for association with EGPA. Genes are shown as colored blocks (see panel A for legend). Traversing lines indicate DNA-DNA physical interactions identified by CHiC-P.

(**A**) **BCL2L11 region.** IgAN= IgA nephropathy, RA= rheumatoid arthritis, CLL= chronic lymphocytic leukemia, Mn= peripheral blood monocyte count, MCV= mean red cell volume, PSC= primary sclerosing cholangitis, At= atopic dermatitis, EC= peripheral blood eosinophil count. CHiC-P data shown for total activated CD4 T cells (10).

(**B**) **TSLP-WDR36 region.** As= asthma, AsH= joint phenotype of both asthma and hayfever, EO= eosinophilic esophagitis, AR= allergic rhinitis, EC= peripheral blood eosinophil count, Al= self-reported allergy. CHiC-P data shown for total activated CD4 T cells (10).

(**C**) **GPA33 region.** Sz= schizophrenia, Astig= refractory astigmastism, Q= QRS duration in Tripanosoma cruzi seropositiivty, L = liver enzyme levels. CHiC-P data shown for GM12878 cell line (11).

(**D**) **LPP-BCL6 region.** NHL= non-Hodkgin's lymphoma, T2D= type 2 diabetes mellitus, Vit= vitiligo, CLL= chronic lymphocytic leukaemia, Sz= schizophrenia, All= self-reported allergy, All*= allergic sensitization, FL= follicular lymphoma, IgE= plasma immunoglobulin E levels, EC= peripheral blood eosinophil count, Vs= interferon response to smallpox vaccine. CHiC-P data shown for total B cells (10).

(**E**) **C5orf56-IRF1-IL5 region.** CD= Crohn's disease, IBD= inflammatory bowel disease, F= fibrinogen levels, As= asthma, As*= severe asthma, Im= composite phenotype of multiple autoimmune diseases, Plt= platelet count, As**= asthma (interaction of genotype with sex), CRP= C-reactive protein levels, EC= peripheral blood eosinophil count. CHiC-P data shown for GM12878 cell line (11).

(**F**) **BACH2 region.** Cog= cognitive performance, Ig= IgG glycosylation, Ob= obesity-related traits, EC= peripheral blood eosinophil count, Ce= celiac disease, TPO= thyroid peroxidase autoantibody positivity, G= Grave's disease, Vit= vitligo, T1D= type 1 diabetes, IBD= inflammatory bowel disease, MS= multiple sclerosis, SLE= systemic lupus erythematosus, TNFr= response to anti-TNF therapy in rheumatoid arthritis. CHiC-P data shown for total activated CD4 T cells (10).

(**G**) **CDK6 region.** RA= rheumatoid arthritis, H= height, EC= eosinophil count, NC= neutrophil count. CHiC-P data shown for total CD4 T cells (10).

(**H**) **10p14 intergenic region.** HL= Hodgkin's lymphoma, ALL = acute lymphoblastic leukemia, RA= rheumatoid arthritis, ALLr = response to chemotherapy in ALL, SLE = systemic lupus erythematosus, CC = colorectal cancer, Mi= migraine, BCC = basal cell carcinoma, TF= tetralogy of Fallot, As =asthma, At= recalcitrant atopic dermatitis in Koreans, Al = self-reported allergy, H1N1= susceptibility to H1N1 influenza, EC = peripheral blood eosinophil count, Str = stroke, Ca = serum calcium, V= IL1beta response to smallpox vaccine. CHiC-P data shown for naive CD4 T cells (10).

**(I) TBX3-MED13L intergenic region.** PSA= prostate specific antigen levels, CC= colorectal carcinoma, Ur= urate levels, BP= blood pressure, AD= Alzheimer's disease, Br= breast carcinoma, EC= peripheral blood eosinophil count, DN= diabetic nephropathy. CHiC-P data shown for GM12878 cell line (11).

(**J**) **CLEC16A-SOCS1 intergenic region.** MS=multiple sclerosis, Aut= autistic spectrum disorders, EC= eosinophil count, CD+Ps= combined phenotype of Crohn's disease and psoriasis, T1D= type 1 diabetes, Ps= psoriasis, VG= vein graft stenosis post coronary artery bypass graft, Ob= obesity-related traits, Bu= bulimia nervosa, Ce= celiac disease. CHiC-P data shown for GM12878 cell line (11).

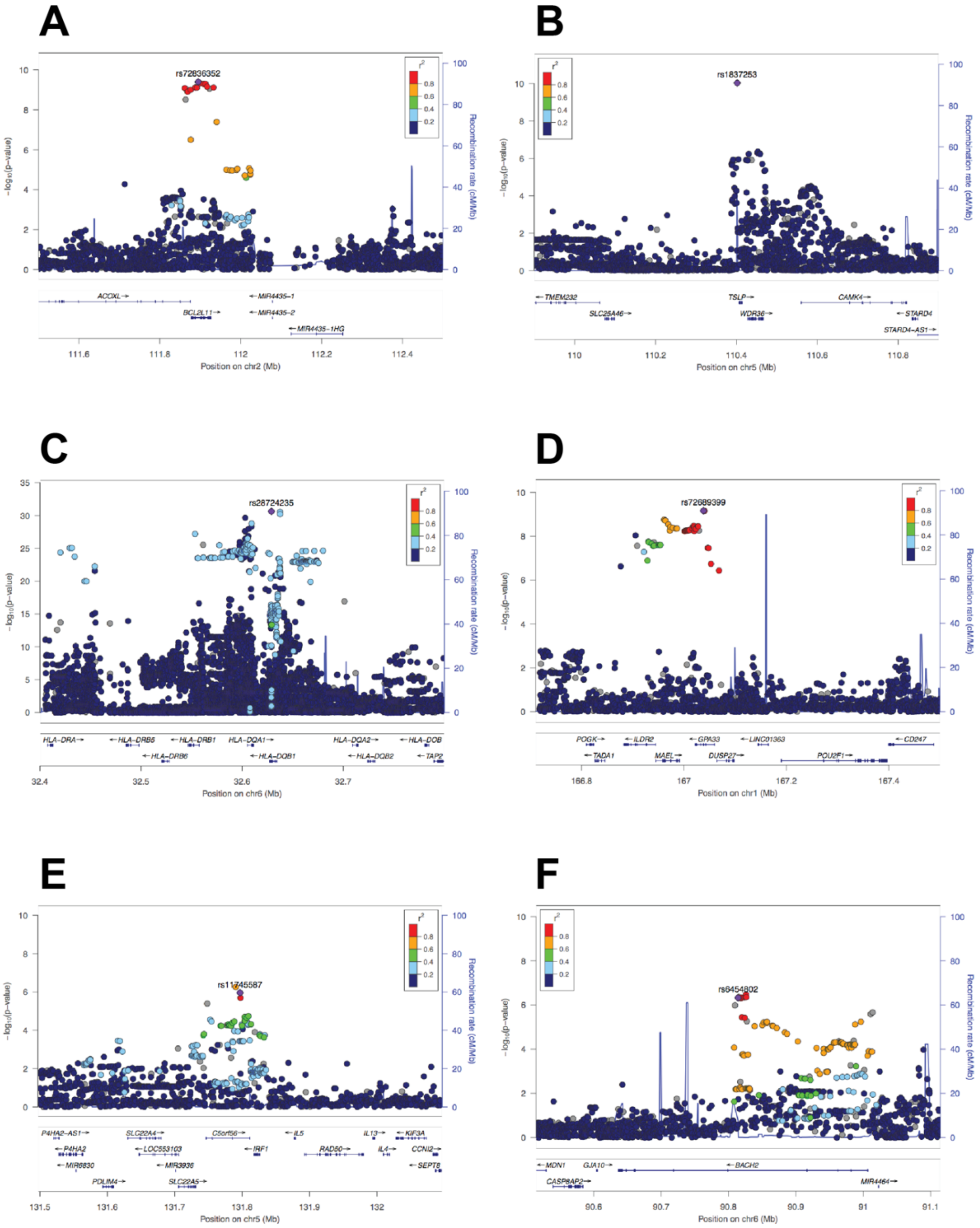
A
rs72836352
ACOXL
BCL2L11
MIR4435-1
MIR4435-2
MIR4435-1HG
Position on chr2 (Mb)
B
rs1837253
TMEM232
SLC25A46
TSLP
WDR36
CAMK4
STARD4
STARD4-AS1
Position on chr5 (Mb)
C
rs28724235
HLA-DRA
HLA-DRB5
HLA-DRB1
HLA-DQA1
HLA-DRB6
HLA-DQB1
HLA-DQA2
HLA-DQB2
HLA-DOB
TAP2
Position on chr6 (Mb)
D
rs72689399
POGK
TADA1
ILDR2
MAEL
GPA33
DUSP27
LINC01363
POU2F1
CD247
Position on chr1 (Mb)
E
rs11745587
P4HA2-AS1
P4HA2
MIR6830
PDLIM4
SLC22A4
LOC553103
MIR3936
SLC22A5
C5orf56
IRF1
IL5
RAD50
IL13
IL4
KIF3A
CCNI2
SEPT8
Position on chr5 (Mb)
F
rs6454802
MDN1
GJA10
CASP8AP2
BACH2
MIR4464
Position on chr6 (Mb)
−log10(p-value)
Recombination rate (cM/Mb)
r2

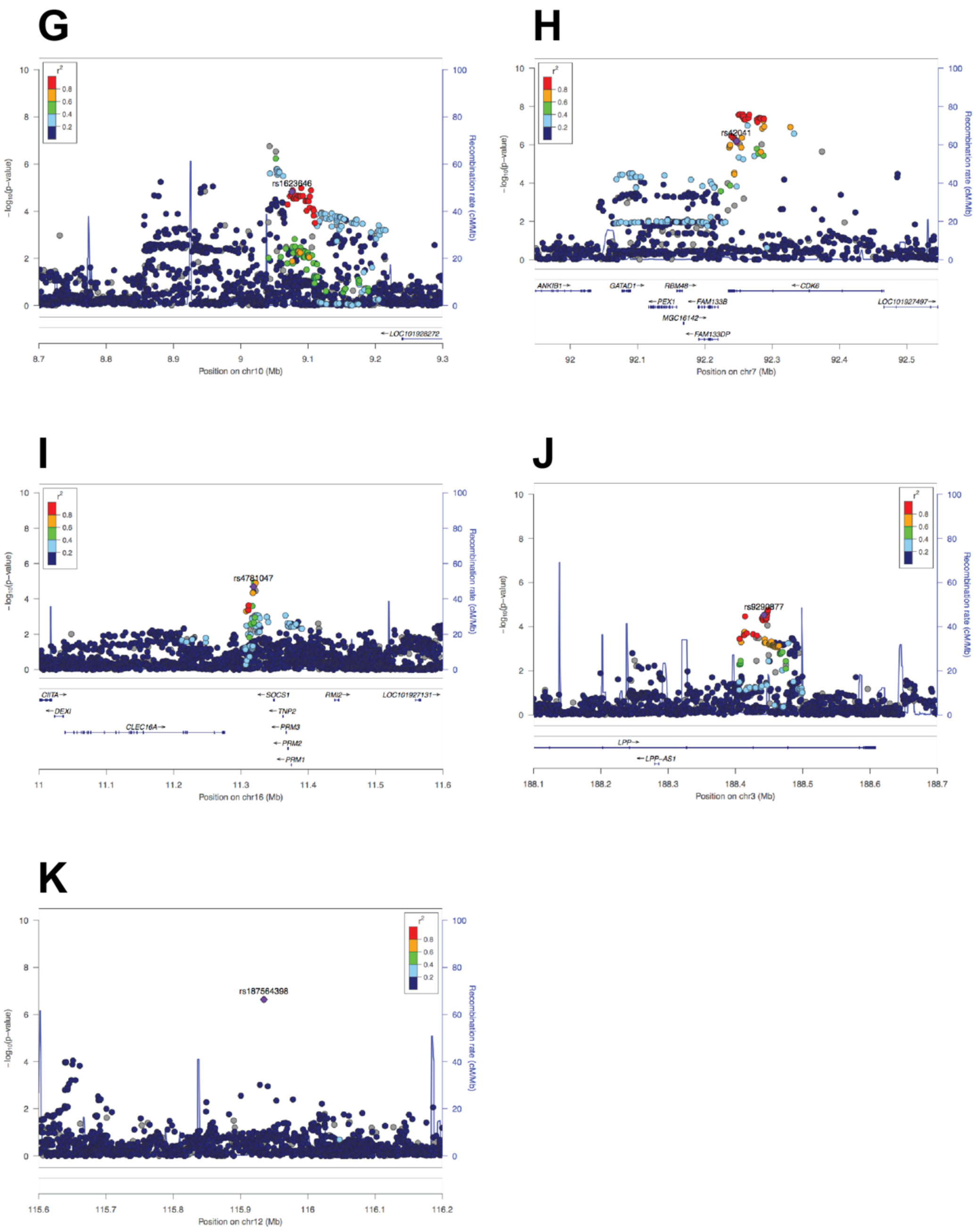

**Supplementary Figure 7. Locus zoom plots for loci associated with EGPA.** The strength of association with EGPA and LD structure relative to the lead SNP at the A) *BCL2L11*, B) *TSLP*, C) *MHC*, D) GPA33, E) *IRF1/IL5*, F) *BACH2*, G) Chr 10, H) *CDK6*, I) *SOCS1*, J) *LPP* and K) *TBX3* loci.

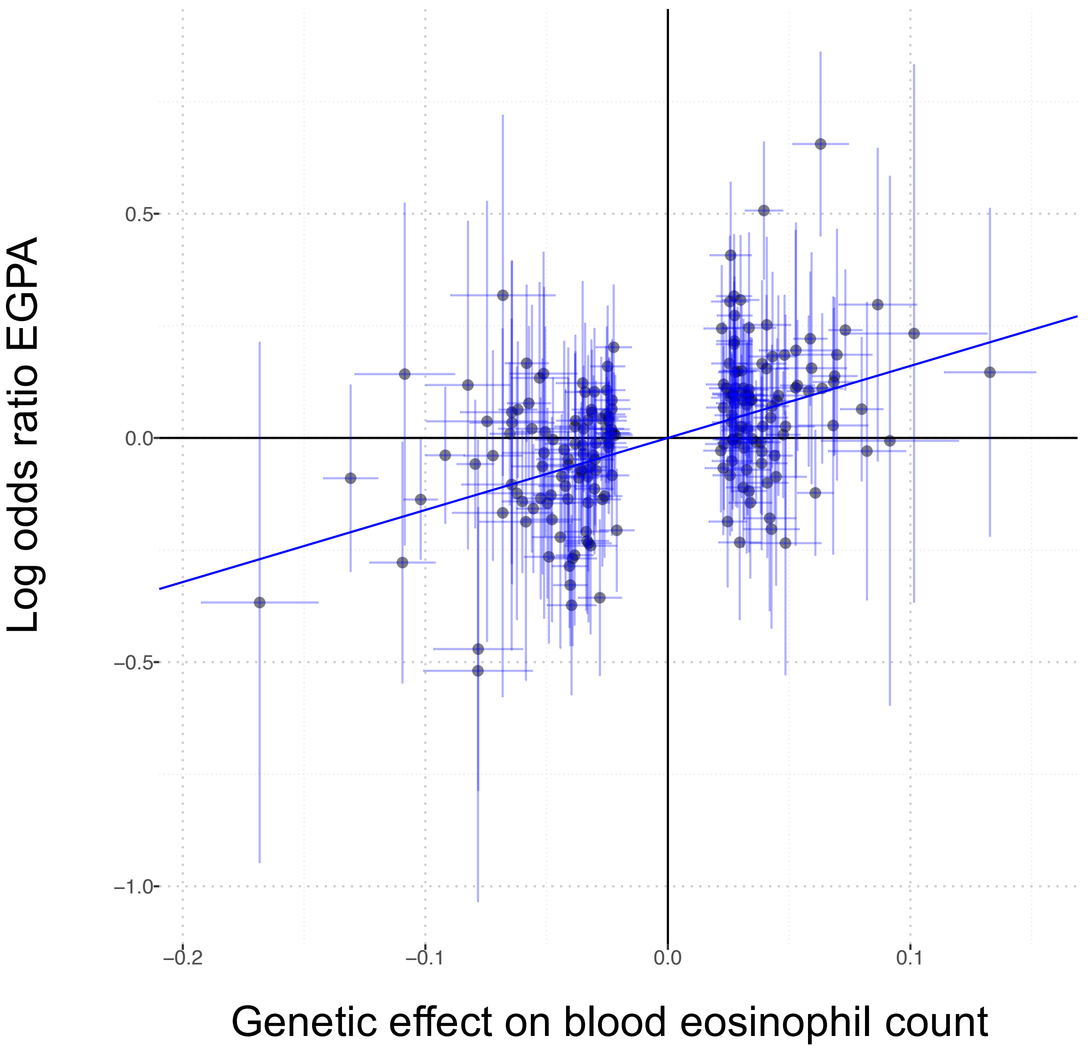

**Supplementary Figure 8. Genetic effects on eosinophil count correlate with risk of EGPA.** Of the 209 conditionally independent genetic variants associated with peripheral blood eosinophil count in the analysis by Astle *et al* (9), 193 (78%) were typed or imputed with INFO score >0.8 in the EGPA dataset. The point estimates for the effect sizes on eosinophil count and on EGPA risk for these 193 SNPs are shown here. Each circle represents a genetic variant. 95% confidence intervals are indicated by the horizontal and vertical bars. The effect size for eosinophil count (x-axis) is the coefficient (the 'beta') for the genotype term in the meta-analysis by Astle *et al*. The natural log of the odds ratio (y-axis) gives the coefficient for the genotype term in the logistic regression in the EGPA GWAS.

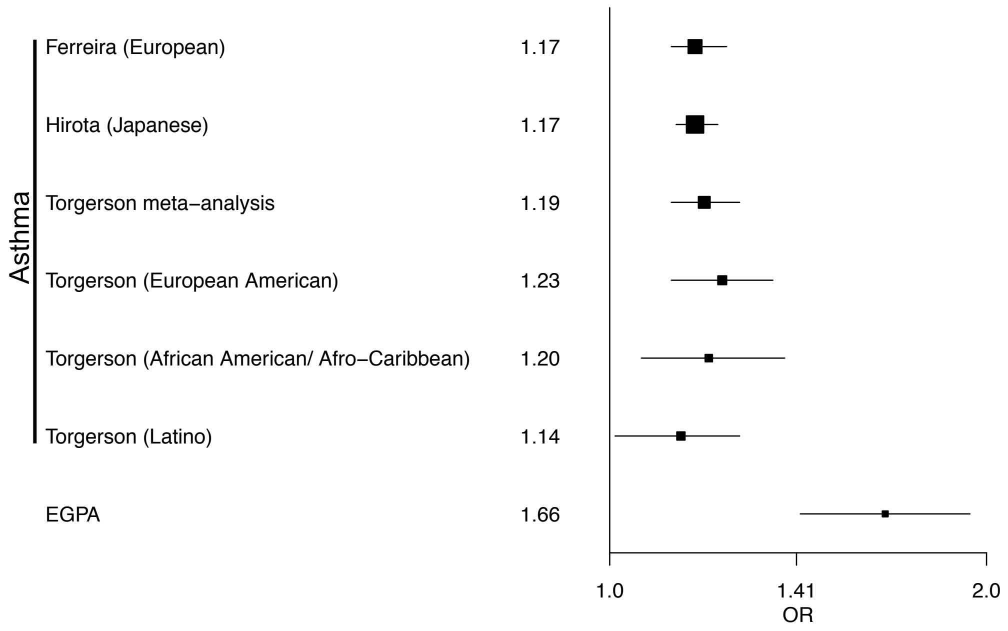

**Supplementary Figure 9. The *TSLP* promoter region variant rs1837253 has a greater effect size in EGPA than in asthma.** Forest plot comparing odds ratio estimates from genome-wide association studies of asthma to EGPA. Black squares indicate estimated odds ratios (also printed numerically in the column to the left of the plot). Horizontal lines indicate 95% confidence intervals. Asthma studies are indicated by the name of the first author, with the ancestry of the cohort studied in parentheses.

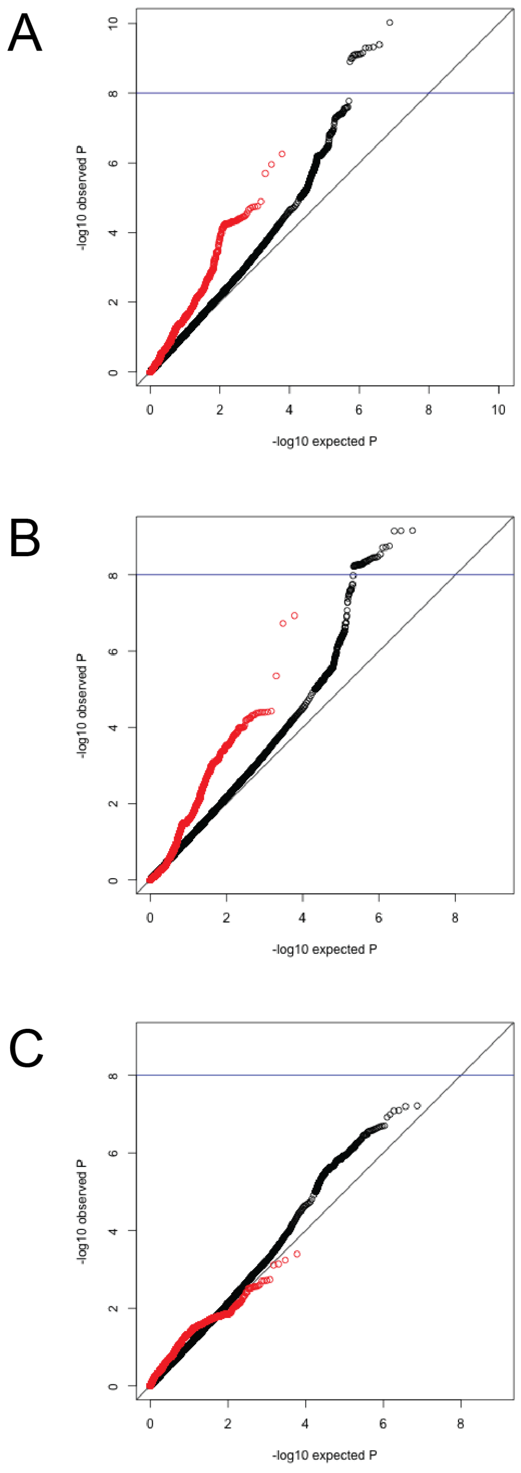

**Supplementary Figure 10. QQ plots of genetic associations in EGPA according to association in IBD**. A) All EGPA cases vs controls. B) ANCA –ve EGPA vs controls. C) MPO+ EGPA vs controls. Black circles indicate all genetic variants in the EGPA study. Red circles represent the subset of genetic variants with genome-wide significance (P <5x10-8) in IBD. IBD summary statistics were taken from the GWAS by Liu et al (12)(European-ancestry individuals).

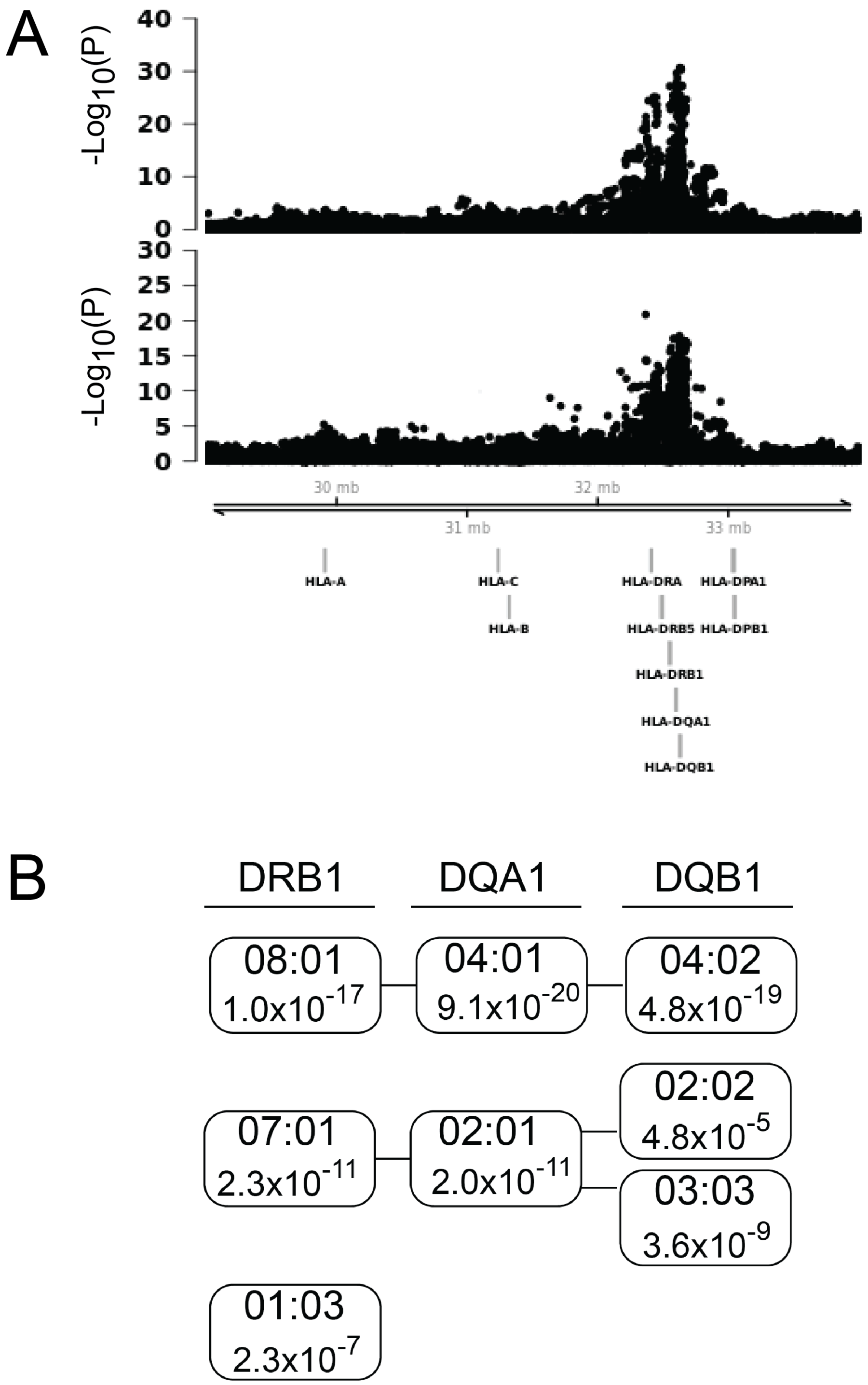

**Supplementary Figure 11. The MHC association with MPO+ve EGPA is localized to the Class II region. (A)** The MHC association signal in MPO+ EGPA (upper plot), and MPO+ ANCA-associated vasculitis (from Lyons et al (13) middle plot) localize to the same interval. The positions of selected Class I and Class II loci are indicated in the lower plot, all co-ordinates are from the hg19 genome build. **(B)** Imputation of classical alleles identifies three MHC haplotypes that confer susceptibility to EGPA.

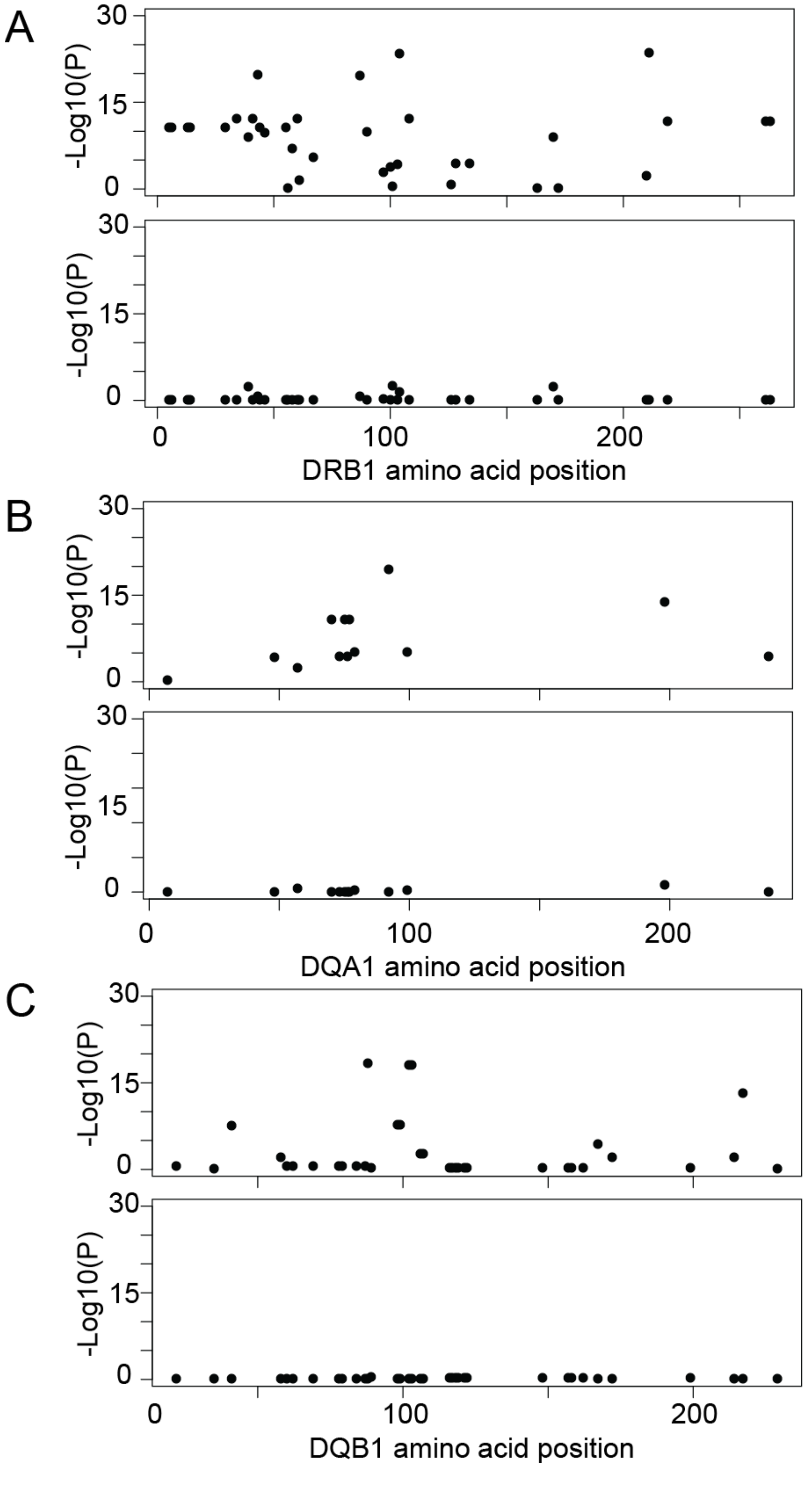

**Supplementary Figure 12. Amino acid positions in HLA-DRB1, HLA-DQA1 and HLA-DQB1 associated with susceptibility to EGPA.** Amino acid positions in (**A**) HLA-DRB1, (**B**) HLA-DQA1 and (**C**) HLA-DQB1 associated with susceptibility to EGPA (upper panels) and following conditioning on position 181 in HLA-DRB1 (lower panels)

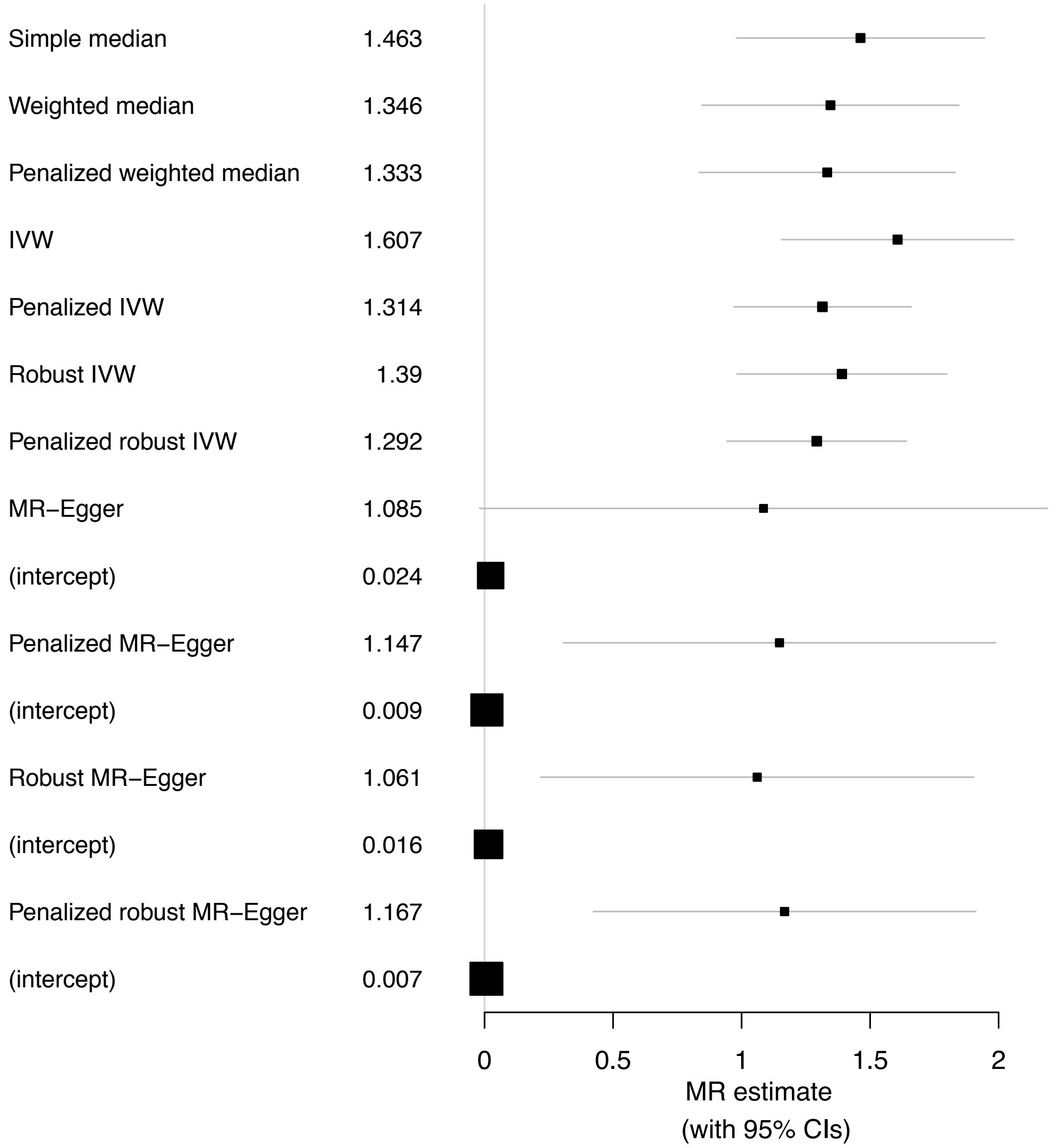

**Supplementary Figure 13. Forest plot of Mendelian randomization estimates for the causal effect of eosinophil count on EGPA**. Point estimates with 95% confidence intervals are shown for multiple Mendelian randomization methods. IVW = inverse variance weighted.

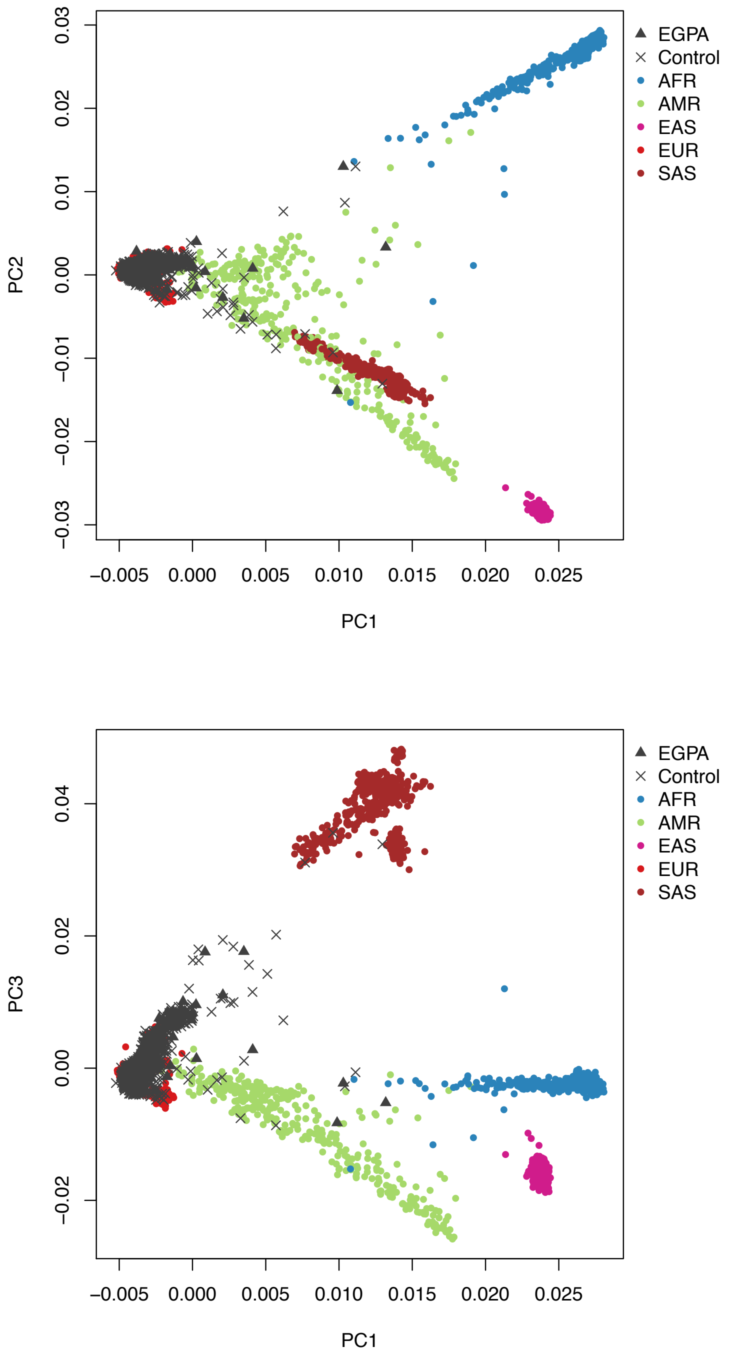

**Supplementary Figure 14. Principal components analysis (PCA) of genotype data.** PCA plots show ancestry of EGPA patients and controls (before removal of non-European ancestry individuals) in relation to 1000 Genomes Project individuals. EGPA patients and controls are coloured dark grey. Non-European ancestry cases and controls were removed prior to subsequent analysis.

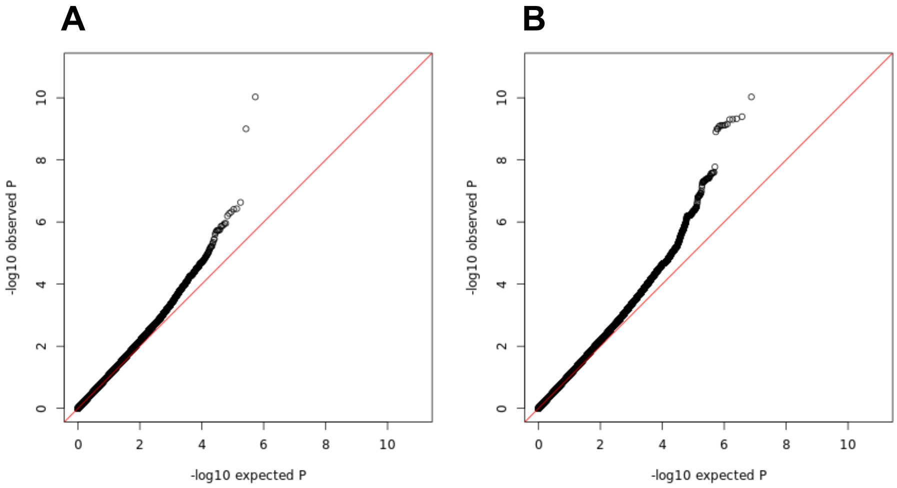

**Supplementary Figure 15. QQ plots for all EGPA cases vs controls.** (A) Directly genotyped SNPs. (B) Directly genotyped SNPs and imputed SNPs with ‘info’ metric >0.9. The line y = x is shown in red. The MHC region has been excluded.

### Supplementary Tables

#### Supplementary Table 1. Criteria for the diagnosis of EGPA from the 'Study to Investigate Mepolizumab in the Treatment of Eosinophilic Granulomatosis With Polyangiitis' (MIRRA[$])

| | |
|---|---|
| A diagnosis of EGPA requires both:<br>-**Asthma**<br>AND<br>-**Eosinophilia** (>1.0x$10^9$/L and/or >10% of total blood leucocytes)<br>PLUS at least 2 of the following additional features of EGPA: | |
| **Positive biopsy** | A biopsy showing histopathological evidence of eosinophilic vasculitis, or perivascular eosinophilic infiltration, or eosinophil-rich granulomatous inflammation. |
| **Neuropathy** | Either mononeuritis or polyneuropathy demonstrated by a motor deficit or nerve conduction abnormality |
| **Pulmonary infiltrates** | Non-fixed |
| **Sino-nasal abnormality** | |
| **Cardiomyopathy** | Established by echocardiography or cardiac magnetic resonance imaging |
| **Glomerulonephritis** | Hematuria, red cell casts, proteinuria |
| **Alveolar haemorrhage** | Confirmed by bronchoalveolar lavage |
| **Palpable purpura** | |
| **Positive ANCA** | Positive MPO or PR-3 ANCA |

[$] https://clinicaltrials.gov/ct2/show/record/NCT02020889

**Supplementary Table 2. Breakdown of 542 cases and 6717 controls by country and center (primary cohort).**

| Country | Cases | Controls |
|---|---|---|
| United Kingdom and Republic of Ireland | 98 | 5466 (EPIC) |
| Germany | 152 | 285 |
| Czech Republic | 5 | 144 |
| Poland | 48 | 119 |
| France | 69 | 0 |
| Italy | 120 | 267 |
| Spain | 29 | 93 |
| Sweden | 21 | 343 |

UK controls from the European Prospective Investigation into Cancer and Nutrition (EPIC) study.

**Supplementary Table 3. Ethics approval from each contributing centre**

| Centre/DNA bank | Ethics number | Ethics committee |
|---|---|---|
| **Overarching GWAS ethics** | 10/H0308/1 | Cambridgeshire 2 Research Ethics Committee |
| **Watts DNA bank** | MREC 03/0/118 | MREC for Scotland |
| **University of Erlangen-Nuremberg** | No. 3604 | Ethics Committee of the University of Erlangen-Nuremberg |
| **Poland** | KBET/201/B/2011 | Jagiellonian University Ethics Committee |
| **Klinikum Bad Bramstedt** | AZ 13-114 | Ethics Committee of the University of Lübeck |
| **Lund University** | Dnr 2010/29 | The Regional Ethical Review Board, Lund, Sweden |
| **University Hospital of Parma** | 29932-08/10/2008 | Ethics Committee of Parma University Hospital |
| **Paris** | 2009-A01331-56 | Comite de Protection des Personnes Ile de France X |
| **Karolinska University Hospital** | 2008/1143-31 | Regional Ethics Committee in Stockholm. |
| **Hospital Clinic Barcelona** | HCB/2016/0274 | Ethics committee of the Hospital Clínic of Barcelona |
| **San Raffaele Hospital - Milan** | Autoimmuno-mol Protocol | Ethics Committee of San Raffaele Hospital, Milan, Italy |
| **Uppsala University** | 2011/241/2 | Regional Ethics Committee in Uppsala, Sweden |
| **University Hospital Prague** | 1738/07 (S-IV) | Ethics Committee of the General University Hospital in Prague |
| **St James's Hospital, Dublin** | 01/03/2010 | SJH/AMNCH Research Ethics Committee |

**Supplementary Table 4. Genetic association analysis using a linear mixed model (LMM-Bolt).**

| CHR | SNP | BP | P PCA | P MLM | | |
|---|---|---|---|---|---|---|
| | | | | EGPA | ANCA -ve | MPO +ve |
| 2 | rs72836352 | 111894874 | $4.03x^{-10}$ | $2.60x^{-11}$ | - | - |
| 5 | rs1837253 | 110401872 | $9.33x^{-11}$ | $1.20x^{-10}$ | - | - |
| 6 | rs28724235 | 32628915 | $1.63x^{-11}$ | $5.90x^{-09}$ | - | $8.5x^{-28}$ |
| 7 | rs42046 | 92252203 | $2.57x^{-08}$ | $6.50x^{-08}$ | - | - |
| 1 | rs72689399 | 167038121 | $1.80x^{-06}$ | $1.90x^{-06}$ | $8.6x^{-10}$ | - |
| 5 | rs11745587 | 131796922 | $1.10x^{-06}$ | $6.30x^{-07}$ | - | - |
| 6 | rs6454802 | 90814199 | $4.72x^{-07}$ | $4.00x^{-07}$ | - | - |
| 10 | rs1623646 | 9076230 | $1.49x^{-05}$ | $1.10x^{-05}$ | - | - |
| 16 | rs4781047 | 11318537 | $2.04x^{-05}$ | $1.30x^{-03}$ | - | - |
| 3 | rs9290877 | 188442480 | $3.00x^{-05}$ | $1.80x^{-06}$ | - | - |
| 12 | rs187564398 | 115934855 | $2.32x^{-07}$ | $2.50x^{-05}$ | - | - |

**Supplementary Table 5. Replication cohort case demographics by country of origin**

| | Germany | Italy |
|---|---|---|
| Number | 49 | 101 |
| Gender (M/F) | 16/33 | 43/58 |
| ANCA -ve | 42 | 56 |
| ANCA +ve | 6 | 43 |
| MPO ANCA+ve | 4 | 41 |

**Supplementary Table 6. Meta-analysis of genetic associations with EGPA in the primary and replication cohorts**

| | | | Total EGPA | | | | | |
|---|---|---|---|---|---|---|---|---|
| | | | Primary Cohort N = 542 cases N= 6717 controls | | Replication Cohort N = 142 cases N= 121 controls | | Combined Cohort N = 684 cases N =6838 controls | |
| chr | Gene | SNP | P | Beta | P | Beta | P | Beta |
| 2 | *BCL2L11* | rs72836352 | $4.03\times10^{-10}$ | 0.66 | 0.008 | 0.72 | $1.07\times10^{-11}$ | 0.66 |
| 5 | *TSLP* | rs1837253 | $9.33\times10^{-11}$ | 0.51 | 0.015 | 0.53 | $4.67\times10^{-12}$ | 0.51 |
| 6 | *HLA-DQ* | rs28724235 | $1.63\times10^{-11}$ | 0.98 | 0.027 | 0.75 | $1.65\times10^{-12}$ | 0.95 |
| 1 | *GPA33* | rs72689399 | $1.80\times10^{-06}$ | 1.34 | 0.125 | 1.22 | $5.36\times10^{-07}$ | 1.34 |

**Supplementary Table 7. Direction of effect at EGPA variants on eosinophil count and asthma risk**

| Chr | SNP | Typed or imputed | Variant type* | Risk allele for EGPA (major/minor) | Effect on eosinophil count (9) | Effect on asthma risk (14) | Effect on asthma in UKBB |
|---|---|---|---|---|---|---|---|
| 2 | rs72836352 | imputed | *BCL2L11* intron variant, NMD transcript variant | T (minor) | ↑ | na | NS |
| 5 | rs1837253 | typed | *TSLP* upstream gene variant | C (major) | ↑ | ↑ | ↑ |
| 7 | rs42046 | typed | *CDK6* intron variant, upstream gene variant | G (minor) | ↑ | na | NS |
| 1 | rs72689399 | imputed | *GPA33* intron variant, non-coding transcript variant, NMD transcript variant | T (minor) | na | na | NS |
| 5 | rs11745587 | typed | *C5orf56* 3-prime UTR variant, intron variant, downstream gene variant | A (minor) | ↑ | ↑ | ↑ |
| 6 | rs6454802 | typed | *BACH2* intron variant, non-coding transcript variant | C (major) | ↑ | ↑ | ↑ |
| 10 | rs1623646 | typed | intergenic | T (major) | ↑ | ↑ | ↑ |
| 16 | rs4781047 | typed | Non-coding transcript exon variant, non-coding transcript variant | A (minor) | ↑ | na | NS |
| 3 | rs9290877 | typed | *LPP* intron variant, non-coding transcript variant | C (minor) | ↑ | na | ↑ |
| 12 | rs187564398 | typed | intergenic | A (minor) | ↑[$] | na | NS |

* from Ensembl Variant Effect Predictor tool; na, data not available; NS, not genome-wide significant, UTR, untranslated region; NMD, nonsense-mediated decay. [$]not genome-wide significant: P 0.007.

**Supplementary Table 8: ANCA status according to country of recruitment**

| Countries | ANCA negative (N) | MPO positive (N) | % MPO positive |
|---|---|---|---|
| Czech Republic | 2 | 3 | 60.0 |
| France | 49 | 19 | 27.9 |
| Germany | 129 | 22 | 14.6 |
| Italy | 56 | 58 | 50.9 |
| Poland | 36 | 5 | 12.2 |
| Spain | 12 | 16 | 57.1 |
| Sweden | 12 | 9 | 42.9 |
| UK & Republic of Ireland | 62 | 29 | 31.9 |

**Supplementary Table 9: associations of ANCA status with clinical features using logistic regression with adjustment for country of origin.**

| *Clinical feature* | *Nominal P value* | *Bonferroni adjusted P value* | *Odds ratio (95% CI)* |
|---|---|---|---|
| **Neuropathy** | **$1.66 \times 10^{-5}$** | **$1.32 \times 10^{-4}$** | **2.73 (1.73, 4.32)** |
| **Lung infiltrates** | **0.0031** | **$2.46 \times 10^{-2}$** | **0.52 (0.34, 0.80)** |
| ENT | 0.43 | 1.0 | 0.80 (0.47, 1.38) |
| **Cardiomyopathy** | **0.00051** | **$4.07 \times 10^{-3}$** | **0.39 (0.23, 0.66)** |
| **Glomerulonephritis** | **$5.49 \times 10^{-6}$** | **$4.39 \times 10^{-5}$** | **3.79 (2.13, 6.70)** |
| Lung haemorrhage | 0.47 | 1.0 | 1.47 (0.52, 4.11) |
| Purpura | 0.11 | 0.94 | 0.68 (0.42, 1.10) |
| Positive biopsy | 0.57 | 1.0 | 0.88 (0.58, 1.35) |

Analysis confined to ANCA –ve and MPO +ve patients (n = 358+161= 519). Patients who were PR3 positive, or who were ANCA positive by immunofluorescence without MPO or PR3 antibodies were excluded. Odds ratios are for MPO positivity (i.e. positive odds ratios indicate increased prevalence of the clinical feature in MPO positive cases). Significant associations are highlighted in bold.

**Supplementary Table 10. Meta-analysis of genetic associations with EGPA subsets in the primary and replication cohorts**

| EGPA subset | | | | Primary Cohort | | Replication Cohort | | Combined Cohort | |
|---|---|---|---|---|---|---|---|---|---|
| | chr | Gene | SNP | P | Beta | P | Beta | P | Beta |
| MPO+ve EGPA* | | | | | | | | | |
| | 6 | *HLA-DQ* | rs28724235 | $2.35 \times 10^{-31}$ | 3.03 | 0.003 | 1.43 | $2.04 \times 10^{-31}$ | 2.67 |
| ANCA -ve EGPA‡ | | | | | | | | | |
| | 1 | *GPA33* | rs72689399 | $6.85 \times 10^{-10}$ | 2.04 | 0.265 | 1.06 | $5.91 \times 10^{-10}$ | 1.94 |

* MPO+ve EGPA, primary cohort N = 161, replication cohort N = 43, combined cohort N = 204

‡ ANCA –ve EGPA, primary cohort N = 358, replication cohort N = 94, combined cohort N = 452

### Supplementary Table 11. Non-MHC EGPA-associated loci and other diseases

| Position (hg19 build) | Lead EGPA-associated SNP | Candidate gene(s) | | Other diseases or traits associated with the EGPA lead SNP or linked SNPs in high LD ($r^2$>0.6) | | | Selected potentially relevant diseases / traits associated with variants within 1 MB of EGPA associated variant but not in LD with EGPA associated variant |
|---|---|---|---|---|---|---|---|
| | | | **Disease or trait** | **Directional concordance of effect on EGPA vs effect on other trait** | **Trait-associated reported LD proxy SNP (where different from EGPA SNP)** | **$r^2$ to EGPA SNP (where applicable)** | |
| chr1: 167038121 | rs72689399 | GPA33 | - | N/A | - | - | Celiac disease, Systemic sclerosis (15-18), Celiac disease or Rheumatoid arthritis (19), *Rheumatoid arthritis** (20), Allergic disease (asthma, hay fever or eczema) (21), Asthma (22) |
| chr2: 111894874 | rs72836352 | *BCL2L11, ACOXL, MIR4435-2HG* | Eosinophil count (9) | + | N/A | N/A | Chronic lymphocytic leukemia (23, 24), Rheumatoid arthritis (25), IgA Nephropathy (26), Atopic dermatitis* (27), Allergic sensitization* (28), Monocyte count* (29) |
| | | | Monocyte count (9) | + | N/A | N/A | |
| | | | PSC (30) | + | N/A | N/A | |
| | | | *Asthma* (31)* | + | rs72837826 | 0.971 | |
| | | | *Self-reported nasal polyps* (31)* | + | rs72836346 | 0.73676 | |
| | | | PSC (32) | + | rs6720394 | 0.687 | |

| | | | | | | | |
|---|---|---|---|---|---|---|---|
| | | | *IBD** (12) | + | rs72837849 | 0.687 | |
| chr5: 110401872 | rs1837253 | *TSLP* | Eosinophil count (9) | + | N/A | N/A | Asthma* (33),<br>Asthma and hayfever (34),<br>Allergic sensitization (28),<br>IgE grass sensitization (35),<br>Allergic rhinitis (35),<br>Self-reported allergy (36)3),<br>Eosinophilic esophagitis (37, 38),<br>Eosinophilic esophagitis (pediatric) (39),<br>Blood eosinophil counts (40) |
| | | | Asthma (41, 42) (31) | + | N/A | N/A | |
| | | | Severe asthma* (14, 43)** | + | N/A | N/A | |
| | | | Allergic rhinitis (31) | | N/A | N/A | |
| | | | Phenotype of both asthma and hayfever (34) | | N/A | N/A | |
| | | | Nasal polyp (31) | + | N/A | N/A | |
| | | | Allergic disease (21) | + | N/A | N/A | |
| | | | Hayfever, allergic rhinitis or eczema (31) | + | N/A | N/A | |

| Position (hg19 build) | Lead EGPA-associated SNP | Candidate gene(s) | Other diseases or traits associated with the EGPA lead SNP or linked SNPs in high LD ($r^2$>0.6) | | | | Selected potentially relevant diseases / traits associated with variants in proximity to candidate gene(s) but not in LD with EGPA associated variant |
|---|---|---|---|---|---|---|---|
| | | | **Disease or trait** | **Directional concordance of effect on EGPA vs effect on other trait** | **Trait-associated reported LD proxy SNP (where different from EGPA SNP)** | **$r^2$ to EGPA SNP (where applicable)** | |
| chr3: 188442480 | rs9290877 | *LPP, BCL6* | Asthma (31) | + | N/A | N/A | Self-reported allergy (36), Allergic sensitization (28), Vitiligo (44, 45), Celiac disease (46), SLE (47), Autoimmune thyroid diseases (48), *Multiple sclerosis** (49), CLL (23) |
| | | | Eosinophil count ((9) | + | N/A | N/A | |
| | | | *Plasma IgE ** (50) | + | N/A | N/A | |
| | | | *Allergic disease * (21)* | + | N/A | N/A | |
| chr5: 131796922 | rs11745587 | *IL5, IRF1, C5orf56* | Eosinophil count (9) | + | N/A | N/A | Eosinophil counts (40)9), IgE levels (51), Atopic dermatitis (52), CRP levels (53), Fibrinogen levels (54), Giant cell arteritis (55), Platelet count (56) |
| | | | Platelet count | + | N/A | N/A | |
| | | | Hayfever, allergic rhinitis or eczema (31) | + | N/A | N/A | |
| | | | Asthma (31) | + | N/A | N/A | |
| | | | *Severe asthma** (43) | ? | N/A | N/A | |
| | | | *Plasma IgE** (50) | ? | N/A | N/A | |
| | | | *Allergic disease * (21)* | + | rs6894249 | 0.821 | |
| | | | Crohn's, IBD (12, 57) | - | N/A | N/A | |
| | | | JIA (58) | ? | rs6894249 | 0.821 | |
| chr6: 90814199 | rs6454802 | *BACH2* | Eosinophil count (9) | + | N/A | N/A | Celiac disease (46), Type 1 diabetes (59), Vitiligo (60), Thyroid autoantibody positivity (61), Grave's disease* (62)1), SLE* (63), Ankylosing spondylitis (64, 65), *Rheumatoid arthritis** (66) |
| | | | Lymphocyte count (9) | + | N/A | N/A | |
| | | | *Basophil count * (9)* | + | N/A | N/A | |
| | | | Asthma (31) | + | N/A | N/A | |
| | | | Allergic disease (21) | + | N/A | N/A | |
| | | | Hayfever, allergic rhinitis or eczema* (31) | + | rs12196749 | 0.86626 | |
| | | | Nasal polyp (31) | + | N/A | N/A | |

| | | | | | | | |
|---|---|---|---|---|---|---|---|
| | | | *Celiac disease** (67) | ? | rs7753008 | 0.971 | |
| | | | *IBD** (12) | + | rs72925996 | 0.673 | |
| | | | MS (68) | ? | rs12212193 | 0.641 | |
| | | | *PSC* (30)* | - | rs56353819 | 0.656 | |
| | | | Hypothyroidism | - | N/A | N/A | |
| chr7: 92252203 | rs42046 | *CDK6* | Eosinophil count (9) | + | rs4272 | 0.61 | Blood neutrophil count (29) |
| | | | Monocyte count (9) | + | N/A | N/A | |
| | | | *RA** (25) | - | N/A | N/A | |

| | Supplementary Table 11 (cont'd). Non-MHC EGPA-associated loci and other diseases | | | | | | |
|---|---|---|---|---|---|---|---|
| chr10: 9076230 | rs1623646 | Intergenic (*GATA3*) | Eosinophil count (9) | + | N/A | N/A | Asthma (41), Self-reported allergy (36), Allergic rhinitis (69), Allergic disease (asthma, hay fever or eczema) (21), Atopic dermatitis* (70), RA (25), SLE (63) |
| | | | Neutrophil count (9) | - | N/A | N/A | |
| | | | Allergic disease (21) | + | N/A | N/A | |
| | | | Hayfever, allergic rhinitis or eczema (31) | + | N/A | N/A | |
| | | | Self-reported asthma (31) | + | N/A | N/A | |
| | | | Doctor diagnosed asthma (31) | + | N/A | N/A | |
| chr12: 115934855 | rs187564398 | Intergenic (*TBX3-MED13L*) | - | | N/A | N/A | Breast cancer (71), Colorectal cancer* (72), Diabetic nephropathy* (73) |
| chr16: 11318537 | rs4781047 | *CLEC16A, SOCS1* | Eosinophil count (9) | + | N/A | N/A | Asthma and hayfever (34), Atopic dermatitis (52), Celiac disease (46), Type 1 diabetes (59, 74, 75), Multiple sclerosis (68), SLE (76), PBC (77, 78), IBD (12, 57), Leprosy (79), Psoriasis* (80), Self-reported allergy* (36), Allergic rhinitis* (35), Vein graft stenosis after CABG* (81) |
| | | | Nasal polyp (31) | + | rs193763 | 0.849 | |
| | | | *Asthma** | + | rs10451095 | 0.656 | |
| | | | *PSC* (30)* | + | rs10451095 | 0.656 | |

**Supplementary Table 12.** Evidence to support biological plausibility of EGPA-associated variants

| Chr | Variant | Other relevant traits with GWAS significant signals in the region | Candidate genes | eQTL or pQTL? | Strength of additional evidence | Experimental data | Other |
|---|---|---|---|---|---|---|---|
| 1 | rs72689399 | - | *GPA33* | Yes* | intermediate | GPA33 plays a role in maintaining epithelial barrier function; KO mice exhibit increased intestinal permeability and increased severity of DSS-induced colitis (82) | |
| 2 | rs72836344 | EC, Asthma, nasal polyps, PSC | *BCL2L11* | Yes | strong | KO mouse: defective apoptosis of immune cells; autoimmunity (83-86) | |
| | | | *MORRBID* | No | strong | MORRBID KO: deficient in eosinophils. (87)<br><br>MORRBID regulates myeloid cell survival via BCL2L11 expression (87)<br><br>MORRBID expression higher in HES patients cf controls. (87)<br><br>MORRBID expression correlates with IL5 expression. (87) | |
| | | | *ACOXL* | No | - | - | |
| 3 | rs9290877 | EC, Asthma, Allergic disease | *LPP* | Yes | - | - | GWAS signals for allergy and immune-mediated diseases in this region at variants independent of EGPA hit |
| | | | *BCL6* | No | strong | BCL6-deficient mice die of overwhelming eosinophilic inflammation characterized by | |

| | | | | | | | |
|---|---|---|---|---|---|---|---|
| | | | | | | myocarditis and pulmonary vasculitis (88) | |
| 5 | rs1837253 | EC,<br>Asthma,<br>Nasal polyps,<br>Combined asthma & hayfever | *TSLP* | Yes | strong | TSLP drives eosinophilia and enhanced TH2 responses through effects on mast cells, group 2 innate lymphoid cells (ILC2), and dendritic cells | Drugs targeting TSLP in development for asthma (89, 90)<br><br>GWAS signals for other eosinophilic diseases in this region at variants independent of EGPA hit (28, 34-40) |
| 5 | rs11745587 | EC,<br>Asthma<br><br>QTL for plasma tryptophan levels$ | *IRF1* | Yes | intermediate | Important immune transcriptional regulator | |
| | | | *IL5* | No | strong | Archetypal “eosinophilic” cytokine (91, 92) | RCT evidence for anti-IL5 therapy in EGPA and eosinophilic asthma (8, 93) |
| | | | *C5orf56* | Yes | no | - | |
| | | | *IL4* | No | strong | | |
| 6 | rs6454802 | EC<br>Asthma<br>Nasal polyps<br>PSC | *BACH2* | Yes | strong | In B cells, BACH2 represses the transcriptional regulator BLIMP1. (94)<br><br>BACH2 also influences multiple facets of T cell differentiation and activity (95, 96)<br><br>BACH2 deficient mice die of eosinophilic pneumonitis (97) | |
| 7 | rs42046 | EC | *CDK6* | Yes | circumstantial | Plays a role in cell cycle regulation | GWAS signal for neutrophil count in this region (9) |

| | | | | | | | |
|---|---|---|---|---|---|---|---|
| 10 | rs1623646 | EC<br>Asthma<br>Hayfever,<br>Allergic rhinitis or eczema | *GATA3* | Yes | strong | GATA3 activation is a key event for Th2 cell differentiation and development (98-101)).<br><br>Directly binds Th2 locus genes and drives pro-eosinophilic cytokines (101)<br><br>GATA3 overexpression leads to eosinophilia in mice (102)<br><br>GATA3 important for development and function of ILC2 cells (103)and invariant NKT cells (104) | |
| 12 | rs187564398 | - | *TBX3* | No | circumstantial | TBX3 upregulated in endothelial cultures in response to VLDL and oxidised VLDL (105) | |
| | | | *MED13L* | No | - | | |
| 16 | rs4781047 | EC<br>Asthma<br>Nasal polyps<br>PSC | *SOCS1* | Yes | intermediate | KO mouse: perinatal lethality as a result of uncontrolled IFN-gamma mediated inflammation (106)<br><br>Important for Treg function(107))<br><br>SOCS1 mimetics reduce inflammation in murine models of autoimmune disorders (108) | |
| | | | *CLECL16A* | Yes | indirect | | CLEC16A region associated with PID (109) |

Abbreviations: EC eosinophil count, IBD inflammatory bowel disease, JIA juvenile idiopathic arthritis, MS multiple sclerosis, PID primary immunodeficiency, PSC primary sclerosing cholangitis

*The eQTL here (bronchial) has low r2 but high D' to the disease hit.

$Tryptophan supplements have been linked to eosinophilic syndromes.

**Supplementary Table 13: Association of classical MHC alleles with MPO+ve EGPA**

| MHC Allele | Unconditioned | | Conditioned on | | | | | |
|---|---|---|---|---|---|---|---|---|
| | | | DQA1*04:01 | | DQA1*04:01 DQA1*02:01 | | DQA1*04:01 DQA1*02:01 DRB1*01:03 | |
| | OR | P | OR | P | OR | P | OR | P |
| HLA-DQA1*04:01 | 7.18 | $9.1 \times 10^{-20}$ | - | - | - | - | - | - |
| HLA-DQB1*04:02 | 6.73 | $4.8 \times 10^{-19}$ | 1.86 | 0.47 | 1.95 | 0.47 | 1.88 | 0.49 |
| HLA-DRB1*08 | 5.97 | $1.0 \times 10^{-17}$ | 0.82 | 0.83 | 1.03 | 0.97 | 1.07 | 0.95 |
| HLA-DQA1*02:01 | 3.05 | $2.0 \times 10^{-11}$ | 3.56 | $2.3 \times 10^{-13}$ | - | - | - | - |
| HLA-DRB1*07:01 | 3.04 | $2.3 \times 10^{-11}$ | 3.56 | $2.6 \times 10^{-13}$ | 0.81 | 0.95 | 0.80 | 0.95 |
| HLA-DQB1*03:03 | 3.35 | $3.6 \times 10^{-9}$ | - | - | - | - | - | - |
| HLA-DRB1*01:03 | 5.96 | $2.3 \times 10^{-7}$ | - | - | 8.96 | $8.0 \times 10^{-10}$ | - | - |
| HLA-DQA1*05:01 | 0.39 | $6.2 \times 10^{-7}$ | - | - | - | - | 0.62 | 0.02 |
| HLA-DQB1*03:01 | 0.46 | $2.6 \times 10^{-5}$ | 0.53 | $7.8 \times 10^{-4}$ | 0.68 | 0.05 | 0.68 | 0.05 |
| HLA-DQB1*02:02 | 2.22 | $4.8 \times 10^{-5}$ | 2.56 | $3.0 \times 10^{-6}$ | - | - | - | - |

**Supplementary Table 14.** Minor allele frequencies at HLA alleles associated with EGPA stratified by country

| HLA allele | | Total cohort | | Czech | | UK | | Germany | | Italy | | Poland | | Spain | | Sweden | |
|---|---|---|---|---|---|---|---|---|---|---|---|---|---|---|---|---|---|
| | | Case | Cont | Case | Cont | Case | Cont | Case | Cont | Case | Cont | Case | Cont | Case | Cont | Case | Cont |
| *HLA DQA1* | 02:01 | 0.19 | 0.13 | 0.30 | 0.15 | 0.23 | 0.14 | 0.13 | 0.09 | 0.22 | 0.13 | 0.16 | 0.17 | 0.29 | 0.16 | 0.14 | 0.07 |
| | 04:01 | 0.07 | 0.02 | 0.10 | 0.02 | 0.04 | 0.02 | 0.06 | 0.02 | 0.09 | 0.02 | 0.05 | 0.03 | 0.10 | 0.03 | 0.12 | 0.04 |
| *HLA DQB1* | 02:02 | 0.13 | 0.10 | 0.10 | 0.08 | 0.17 | 0.10 | 0.09 | 0.07 | 0.17 | 0.10 | 0.13 | 0.12 | 0.22 | 0.14 | nd | 0.05 |
| | 04:02 | 0.07 | 0.02 | 0.10 | 0.02 | 0.04 | 0.02 | 0.06 | 0.02 | 0.09 | 0.03 | 0.05 | 0.03 | 0.10 | 0.03 | 0.12 | 0.04 |
| | 03:03 | 0.06 | 0.05 | 0.20 | 0.07 | 0.08 | 0.05 | 0.04 | 0.04 | 0.05 | 0.04 | 0.04 | 0.05 | 0.07 | 0.02 | 0.19 | 0.05 |
| *HLA DRB1* | 01:03 | 0.02 | 0.01 | nd | 0.01 | 0.03 | 0.01 | 0.01 | 0.004 | 0.02 | 0.004 | nd | nd | 0.07 | 0.005 | 0.02 | 0.004 |

nd, Not detected

**Legend for Supplementary Data Item 1**: Cross-referencing of EGPA-associated variants with variants in high LD in the NHGRI GWAS Catalog.

**Key:**

**egpa.ref_rsid** = rsid of the EGPA associated variant
**ref_hg19_coordinates** = genomic position of the EGPA associated variant (hg19 build)
**ref_a1** = the effect allele of the input EGPA-associated variant with respect to the trait in the GWAS Catalog.
**ref_a2** = the non-effect allele of the input EGPA-associated variant with respect to the trait in the GWAS Catalog.
**gwas.rsid** = rsid of the trait-associated variant in the GWAS catalog.
**a1** = the effect allele (aligned to the + strand) of the trait associated variant.
**a2** = the non- effect allele (aligned to the + strand) of the trait associated variant.
**proxy** = whether the GWAS trait associated variant is an LD proxy for the EGPA SNP. 0= no (i.e. trait-associated variant is exactly the same as the EGPA-associated variant)
**r2** = LD (as r2) between EGPA-associated variant and the trait-associated variant based on the phased haplotypes from 1000 Genomes
**trait** = trait type
**efo** = experimental factor ontology term
**study** = the eQTL study
**pmid** = the PubMed ID of the study
**ancestry** = study population ancestry
**year** = year of study
**beta** = beta coefficient for genotype from the regression model i.e. estimated effect size of the effect allele
**se** = standard error for the genotype coefficient
**p** = p-value
**direction** = direction of effect allele on the trait: + = increases; - = decreases.
**dataset** = the GWAS dataset
**egpa.risk.allele** = the risk allele for EGPA of '**egpa.ref_rsid'**
**concordance** = directional concordance of the variant on EGPA and the GWAS trait. + indicates same direction; - indicates opposite direction.

**Legend for Supplementary Data Item 2**: cross-referencing of EGPA-associated variants with eQTLs.

**Key:**

**egpa.snp.rsid** = rsid of the EGPA associated variant
**ref_hg19_coordinates** = genomic position of the EGPA associated variant (hg19 build)
**ref_a1** = the effect allele of the input EGPA-associated variant with respect to gene expression.
**ref_a2** = the non- effect allele of the input EGPA-associated variant with respect to for gene expression.
**eqtl.rsid** = rsid of the eQTL variant
**hg19_coordinates** = genomic position of the eQTL variant (hg19 build)
**a1** = the effect allele (aligned to the + strand) for gene expression for the input eQTL variant.
**a2** = the non-effect allele (aligned to the + strand) for gene expression for the input eQTL variant.
**proxy** = whether the eQTL variant is an LD proxy for the EGPA SNP. 0= no (i.e. eQTL SNP is exactly the same as the EGPA-associated variant)
**r2** = LD (as r2) between EGPA-associated variant and the eQTL variant based on the phased haplotypes from 1000 Genomes
**dprime** = LD (as r2) between EGPA-associated variant and the eQTL variant based on the phased haplotypes from 1000 Genomes
**trait** = trait type
**efo** = experimental factor ontology term
**study** = the eQTL study
**pmid** = the PubMed ID of the study
**ancestry** = study population ancestry
**year** = year of study
**tissue** = tissue in which the eQTL was identified
**exp_gene** = gene symbol of gene whose expression is associated with the eQTL variant
**exp_ensembl** = ensemble id of gene whose expression is associated with the eQTL variant
**probe** = probe id (where gene expression was measured using microarrays)
**beta** = beta coefficient for genotype i.e. estimated effect size of the effect allele
**se** = standard error for the genotype coefficient
**p** = p-value
**direction** = direction of effect allele on gene expression: + = increases; - = decreases.
**dataset** = eQTL dataset

**Legend for Supplementary Data Item 3**: variants associated with traits in NHGRI GWAS Catalog that lie within +/- 1MB of the EGPA-associated variants).

**Legend for Supplementary Data Item 4**: Mendelian randomisation estimates.

**Legend for Supplementary Data Item 5**: Full EGPA GWAS summary statistics.

### Supplementary references

1. Jennette JC, Falk RJ, Bacon PA, Basu N, Cid MC, Ferrario F, Flores-Suarez LF, Gross WL, Guillevin L, Hagen EC, et al. 2012 revised International Chapel Hill Consensus Conference Nomenclature of Vasculitides. *Arthritis Rheum.* 2013;65(1):1-11.
2. Churg J, and Strauss L. Allergic granulomatosis, allergic angiitis, and periarteritis nodosa. *Am J Pathol.* 1951;27(2):277-301.
3. Reid AJ, Harrison BD, Watts RA, Watkin SW, McCann BG, and Scott DG. Churg-Strauss syndrome in a district hospital. *QJM.* 1998;91(3):219-29.
4. Lanham JG, Elkon KB, Pusey CD, and Hughes GR. Systemic vasculitis with asthma and eosinophilia: a clinical approach to the Churg-Strauss syndrome. *Medicine (Baltimore).* 1984;63(2):65-81.
5. Masi AT, Hunder GG, Lie JT, Michel BA, Bloch DA, Arend WP, Calabrese LH, Edworthy SM, Fauci AS, Leavitt RY, et al. The American College of Rheumatology 1990 criteria for the classification of Churg-Strauss syndrome (allergic granulomatosis and angiitis). *Arthritis Rheum.* 1990;33(8):1094-100.
6. Khoury P, Zagallo P, Talar-Williams C, Santos CS, Dinerman E, Holland NC, and Klion AD. Serum biomarkers are similar in Churg-Strauss syndrome and hypereosinophilic syndrome. *Allergy.* 2012;67(9):1149-56.
7. Rao JK, Allen NB, and Pincus T. Limitations of the 1990 American College of Rheumatology classification criteria in the diagnosis of vasculitis. *Ann Intern Med.* 1998;129(5):345-52.
8. Wechsler ME, Akuthota P, Jayne D, Khoury P, Klion A, Langford CA, Merkel PA, Moosig F, Specks U, Cid MC, et al. Mepolizumab or Placebo for Eosinophilic Granulomatosis with Polyangiitis. *N Engl J Med.* 2017;376(20):1921-32.
9. Astle WJ, Elding H, Jiang T, Allen D, Ruklisa D, Mann AL, Mead D, Bouman H, Riveros-Mckay F, Kostadima MA, et al. The Allelic Landscape of Human Blood Cell Trait Variation and Links to Common Complex Disease. *Cell.* 2016;167(5):1415-29 e19.
10. Javierre BM, Burren OS, Wilder SP, Kreuzhuber R, Hill SM, Sewitz S, Cairns J, Wingett SW, Varnai C, Thiecke MJ, et al. Lineage-Specific Genome Architecture Links Enhancers and Non-coding Disease Variants to Target Gene Promoters. *Cell.* 2016;167(5):1369-84 e19.
11. Mifsud B, Tavares-Cadete F, Young AN, Sugar R, Schoenfelder S, Ferreira L, Wingett SW, Andrews S, Grey W, Ewels PA, et al. Mapping long-range promoter contacts in human cells with high-resolution capture Hi-C. *Nat Genet.* 2015;47(6):598-606.
12. Liu JZ, van Sommeren S, Huang H, Ng SC, Alberts R, Takahashi A, Ripke S, Lee JC, Jostins L, Shah T, et al. Association analyses identify 38 susceptibility loci for inflammatory bowel disease and highlight shared genetic risk across populations. *Nat Genet.* 2015;47(9):979-86.

13. Lyons PA, Rayner TF, Trivedi S, Holle JU, Watts RA, Jayne DR, Baslund B, Brenchley P, Bruchfeld A, Chaudhry AN, et al. Genetically distinct subsets within ANCA-associated vasculitis. *N Engl J Med.* 2012;367(3):214-23.
14. Moffatt MF, Gut IG, Demenais F, Strachan DP, Bouzigon E, Heath S, von Mutius E, Farrall M, Lathrop M, Cookson WO, et al. A large-scale, consortium-based genomewide association study of asthma. *N Engl J Med.* 2010;363(13):1211-21.
15. Allanore Y, Saad M, Dieude P, Avouac J, Distler JH, Amouyel P, Matucci-Cerinic M, Riemekasten G, Airo P, Melchers I, et al. Genome-wide scan identifies TNIP1, PSORS1C1, and RHOB as novel risk loci for systemic sclerosis. *PLoS Genet.* 2011;7(7):e1002091.
16. Gorlova O, Martin JE, Rueda B, Koeleman BP, Ying J, Teruel M, Diaz-Gallo LM, Broen JC, Vonk MC, Simeon CP, et al. Identification of novel genetic markers associated with clinical phenotypes of systemic sclerosis through a genome-wide association strategy. *PLoS Genet.* 2011;7(7):e1002178.
17. Gorlova OY, Li Y, Gorlov I, Ying J, Chen WV, Assassi S, Reveille JD, Arnett FC, Zhou X, Bossini-Castillo L, et al. Gene-level association analysis of systemic sclerosis: A comparison of African-Americans and White populations. *PLoS One.* 2018;13(1):e0189498.
18. Radstake TR, Gorlova O, Rueda B, Martin JE, Alizadeh BZ, Palomino-Morales R, Coenen MJ, Vonk MC, Voskuyl AE, Schuerwegh AJ, et al. Genome-wide association study of systemic sclerosis identifies CD247 as a new susceptibility locus. *Nat Genet.* 2010;42(5):426-9.
19. Zhernakova A, Stahl EA, Trynka G, Raychaudhuri S, Festen EA, Franke L, Westra HJ, Fehrmann RS, Kurreeman FA, Thomson B, et al. Meta-analysis of genome-wide association studies in celiac disease and rheumatoid arthritis identifies fourteen non-HLA shared loci. *PLoS Genet.* 2011;7(2):e1002004.
20. Stahl EA, Raychaudhuri S, Remmers EF, Xie G, Eyre S, Thomson BP, Li Y, Kurreeman FA, Zhernakova A, Hinks A, et al. Genome-wide association study meta-analysis identifies seven new rheumatoid arthritis risk loci. *Nat Genet.* 2010;42(6):508-14.
21. Ferreira MA, Vonk JM, Baurecht H, Marenholz I, Tian C, Hoffman JD, Helmer Q, Tillander A, Ullemar V, van Dongen J, et al. Shared genetic origin of asthma, hay fever and eczema elucidates allergic disease biology. *Nat Genet.* 2017;49(12):1752-7.
22. Pickrell JK, Berisa T, Liu JZ, Segurel L, Tung JY, and Hinds DA. Detection and interpretation of shared genetic influences on 42 human traits. *Nat Genet.* 2016;48(7):709-17.
23. Berndt SI, Skibola CF, Joseph V, Camp NJ, Nieters A, Wang Z, Cozen W, Monnereau A, Wang SS, Kelly RS, et al. Genome-wide association study identifies multiple risk loci for chronic lymphocytic leukemia. *Nat Genet.* 2013;45(8):868-76.
24. Speedy HE, Di Bernardo MC, Sava GP, Dyer MJ, Holroyd A, Wang Y, Sunter NJ, Mansouri L, Juliusson G, Smedby KE, et al. A genome-wide association study identifies multiple susceptibility loci for chronic lymphocytic leukemia. *Nat Genet.* 2014;46(1):56-60.
25. Okada Y, Wu D, Trynka G, Raj T, Terao C, Ikari K, Kochi Y, Ohmura K, Suzuki A, Yoshida S, et al. Genetics of rheumatoid arthritis contributes to biology and drug discovery. *Nature.* 2014;506(7488):376-81.
26. Yu XQ, Li M, Zhang H, Low HQ, Wei X, Wang JQ, Sun LD, Sim KS, Li Y, Foo JN, et al. A genome-wide association study in Han Chinese identifies multiple susceptibility loci for IgA nephropathy. *Nat Genet.* 2011;44(2):178-82.

27. Hirota T, Takahashi A, Kubo M, Tsunoda T, Tomita K, Sakashita M, Yamada T, Fujieda S, Tanaka S, Doi S, et al. Genome-wide association study identifies eight new susceptibility loci for atopic dermatitis in the Japanese population. *Nat Genet.* 2012;44(11):1222-6.
28. Bonnelykke K, Matheson MC, Pers TH, Granell R, Strachan DP, Alves AC, Linneberg A, Curtin JA, Warrington NM, Standl M, et al. Meta-analysis of genome-wide association studies identifies ten loci influencing allergic sensitization. *Nat Genet.* 2013;45(8):902-6.
29. Okada Y, Hirota T, Kamatani Y, Takahashi A, Ohmiya H, Kumasaka N, Higasa K, Yamaguchi-Kabata Y, Hosono N, Nalls MA, et al. Identification of nine novel loci associated with white blood cell subtypes in a Japanese population. *PLoS Genet.* 2011;7(6):e1002067.
30. Ji SG, Juran BD, Mucha S, Folseraas T, Jostins L, Melum E, Kumasaka N, Atkinson EJ, Schlicht EM, Liu JZ, et al. Genome-wide association study of primary sclerosing cholangitis identifies new risk loci and quantifies the genetic relationship with inflammatory bowel disease. *Nat Genet.* 2017;49(2):269-73.
31. Neale BM. Rapid GWAS of thousands of phenotypes for 337000 samples in the UK biobank. http://www.nealelab.is/blog/2017/7/19/rapid-gwas-of-thousands-of-phenotypes-for-337000-samples-in-the-uk-biobank.
32. Melum E, Franke A, Schramm C, Weismuller TJ, Gotthardt DN, Offner FA, Juran BD, Laerdahl JK, Labi V, Bjornsson E, et al. Genome-wide association analysis in primary sclerosing cholangitis identifies two non-HLA susceptibility loci. *Nat Genet.* 2011;43(1):17-9.
33. Ferreira MA, Matheson MC, Duffy DL, Marks GB, Hui J, Le Souef P, Danoy P, Baltic S, Nyholt DR, Jenkins M, et al. Identification of IL6R and chromosome 11q13.5 as risk loci for asthma. *Lancet.* 2011;378(9795):1006-14.
34. Ferreira MA, Matheson MC, Tang CS, Granell R, Ang W, Hui J, Kiefer AK, Duffy DL, Baltic S, Danoy P, et al. Genome-wide association analysis identifies 11 risk variants associated with the asthma with hay fever phenotype. *J Allergy Clin Immunol.* 2014;133(6):1564-71.
35. Ramasamy A, Curjuric I, Coin LJ, Kumar A, McArdle WL, Imboden M, Leynaert B, Kogevinas M, Schmid-Grendelmeier P, Pekkanen J, et al. A genome-wide meta-analysis of genetic variants associated with allergic rhinitis and grass sensitization and their interaction with birth order. *J Allergy Clin Immunol.* 2011;128(5):996-1005.
36. Hinds DA, McMahon G, Kiefer AK, Do CB, Eriksson N, Evans DM, St Pourcain B, Ring SM, Mountain JL, Francke U, et al. A genome-wide association meta-analysis of self-reported allergy identifies shared and allergy-specific susceptibility loci. *Nat Genet.* 2013;45(8):907-11.
37. Kottyan LC, Davis BP, Sherrill JD, Liu K, Rochman M, Kaufman K, Weirauch MT, Vaughn S, Lazaro S, Rupert AM, et al. Genome-wide association analysis of eosinophilic esophagitis provides insight into the tissue specificity of this allergic disease. *Nat Genet.* 2014;46(8):895-900.
38. Sleiman PM, Wang ML, Cianferoni A, Aceves S, Gonsalves N, Nadeau K, Bredenoord AJ, Furuta GT, Spergel JM, and Hakonarson H. GWAS identifies four novel eosinophilic esophagitis loci. *Nat Commun.* 2014;5(5593.

39. Rothenberg ME, Spergel JM, Sherrill JD, Annaiah K, Martin LJ, Cianferoni A, Gober L, Kim C, Glessner J, Frackelton E, et al. Common variants at 5q22 associate with pediatric eosinophilic esophagitis. *Nat Genet.* 2010;42(4):289-91.
40. Gudbjartsson DF, Bjornsdottir US, Halapi E, Helgadottir A, Sulem P, Jonsdottir GM, Thorleifsson G, Helgadottir H, Steinthorsdottir V, Stefansson H, et al. Sequence variants affecting eosinophil numbers associate with asthma and myocardial infarction. *Nat Genet.* 2009;41(3):342-7.
41. Hirota T, Takahashi A, Kubo M, Tsunoda T, Tomita K, Doi S, Fujita K, Miyatake A, Enomoto T, Miyagawa T, et al. Genome-wide association study identifies three new susceptibility loci for adult asthma in the Japanese population. *Nat Genet.* 2011;43(9):893-6.
42. Torgerson DG, Ampleford EJ, Chiu GY, Gauderman WJ, Gignoux CR, Graves PE, Himes BE, Levin AM, Mathias RA, Hancock DB, et al. Meta-analysis of genome-wide association studies of asthma in ethnically diverse North American populations. *Nat Genet.* 2011;43(9):887-92.
43. Wan YI, Shrine NR, Soler Artigas M, Wain LV, Blakey JD, Moffatt MF, Bush A, Chung KF, Cookson WO, Strachan DP, et al. Genome-wide association study to identify genetic determinants of severe asthma. *Thorax.* 2012;67(9):762-8.
44. Jin Y, Birlea SA, Fain PR, Gowan K, Riccardi SL, Holland PJ, Mailloux CM, Sufit AJ, Hutton SM, Amadi-Myers A, et al. Variant of TYR and autoimmunity susceptibility loci in generalized vitiligo. *N Engl J Med.* 2010;362(18):1686-97.
45. Tang XF, Zhang Z, Hu DY, Xu AE, Zhou HS, Sun LD, Gao M, Gao TW, Gao XH, Chen HD, et al. Association analyses identify three susceptibility Loci for vitiligo in the Chinese Han population. *J Invest Dermatol.* 2013;133(2):403-10.
46. Dubois PC, Trynka G, Franke L, Hunt KA, Romanos J, Curtotti A, Zhernakova A, Heap GA, Adany R, Aromaa A, et al. Multiple common variants for celiac disease influencing immune gene expression. *Nat Genet.* 2010;42(4):295-302.
47. Morris DL, Sheng Y, Zhang Y, Wang YF, Zhu Z, Tombleson P, Chen L, Cunninghame Graham DS, Bentham J, Roberts AL, et al. Genome-wide association meta-analysis in Chinese and European individuals identifies ten new loci associated with systemic lupus erythematosus. *Nat Genet.* 2016;48(8):940-6.
48. Cooper JD, Simmonds MJ, Walker NM, Burren O, Brand OJ, Guo H, Wallace C, Stevens H, Coleman G, Wellcome Trust Case Control C, et al. Seven newly identified loci for autoimmune thyroid disease. *Hum Mol Genet.* 2012;21(23):5202-8.
49. International Multiple Sclerosis Genetics C, Beecham AH, Patsopoulos NA, Xifara DK, Davis MF, Kemppinen A, Cotsapas C, Shah TS, Spencer C, Booth D, et al. Analysis of immune-related loci identifies 48 new susceptibility variants for multiple sclerosis. *Nat Genet.* 2013;45(11):1353-60.
50. Granada M, Wilk JB, Tuzova M, Strachan DP, Weidinger S, Albrecht E, Gieger C, Heinrich J, Himes BE, Hunninghake GM, et al. A genome-wide association study of plasma total IgE concentrations in the Framingham Heart Study. *J Allergy Clin Immunol.* 2012;129(3):840-5 e21.

51. Weidinger S, Gieger C, Rodriguez E, Baurecht H, Mempel M, Klopp N, Gohlke H, Wagenpfeil S, Ollert M, Ring J, et al. Genome-wide scan on total serum IgE levels identifies FCER1A as novel susceptibility locus. *PLoS Genet.* 2008;4(8):e1000166.
52. Paternoster L, Standl M, Waage J, Baurecht H, Hotze M, Strachan DP, Curtin JA, Bonnelykke K, Tian C, Takahashi A, et al. Multi-ancestry genome-wide association study of 21,000 cases and 95,000 controls identifies new risk loci for atopic dermatitis. *Nat Genet.* 2015;47(12):1449-56.
53. Dehghan A, Dupuis J, Barbalic M, Bis JC, Eiriksdottir G, Lu C, Pellikka N, Wallaschofski H, Kettunen J, Henneman P, et al. Meta-analysis of genome-wide association studies in >80 000 subjects identifies multiple loci for C-reactive protein levels. *Circulation.* 2011;123(7):731-8.
54. Dehghan A, Yang Q, Peters A, Basu S, Bis JC, Rudnicka AR, Kavousi M, Chen MH, Baumert J, Lowe GD, et al. Association of novel genetic Loci with circulating fibrinogen levels: a genome-wide association study in 6 population-based cohorts. *Circ Cardiovasc Genet.* 2009;2(2):125-33.
55. Carmona FD, Vaglio A, Mackie SL, Hernandez-Rodriguez J, Monach PA, Castaneda S, Solans R, Morado IC, Narvaez J, Ramentol-Sintas M, et al. A Genome-wide Association Study Identifies Risk Alleles in Plasminogen and P4HA2 Associated with Giant Cell Arteritis. *Am J Hum Genet.* 2017;100(1):64-74.
56. Gieger C, Radhakrishnan A, Cvejic A, Tang W, Porcu E, Pistis G, Serbanovic-Canic J, Elling U, Goodall AH, Labrune Y, et al. New gene functions in megakaryopoiesis and platelet formation. *Nature.* 2011;480(7376):201-8.
57. Jostins L, Ripke S, Weersma RK, Duerr RH, McGovern DP, Hui KY, Lee JC, Schumm LP, Sharma Y, Anderson CA, et al. Host-microbe interactions have shaped the genetic architecture of inflammatory bowel disease. *Nature.* 2012;491(7422):119-24.
58. Hinks A, Cobb J, Marion MC, Prahalad S, Sudman M, Bowes J, Martin P, Comeau ME, Sajuthi S, Andrews R, et al. Dense genotyping of immune-related disease regions identifies 14 new susceptibility loci for juvenile idiopathic arthritis. *Nat Genet.* 2013;45(6):664-9.
59. Cooper JD, Smyth DJ, Smiles AM, Plagnol V, Walker NM, Allen JE, Downes K, Barrett JC, Healy BC, Mychaleckyj JC, et al. Meta-analysis of genome-wide association study data identifies additional type 1 diabetes risk loci. *Nat Genet.* 2008;40(12):1399-401.
60. Jin Y, Birlea SA, Fain PR, Ferrara TM, Ben S, Riccardi SL, Cole JB, Gowan K, Holland PJ, Bennett DC, et al. Genome-wide association analyses identify 13 new susceptibility loci for generalized vitiligo. *Nat Genet.* 2012;44(6):676-80.
61. Medici M, Porcu E, Pistis G, Teumer A, Brown SJ, Jensen RA, Rawal R, Roef GL, Plantinga TS, Vermeulen SH, et al. Identification of novel genetic Loci associated with thyroid peroxidase antibodies and clinical thyroid disease. *PLoS Genet.* 2014;10(2):e1004123.
62. Chu X, Pan CM, Zhao SX, Liang J, Gao GQ, Zhang XM, Yuan GY, Li CG, Xue LQ, Shen M, et al. A genome-wide association study identifies two new risk loci for Graves' disease. *Nat Genet.* 2011;43(9):897-901.

63. Yang W, Tang H, Zhang Y, Tang X, Zhang J, Sun L, Yang J, Cui Y, Zhang L, Hirankarn N, et al. Meta-analysis followed by replication identifies loci in or near CDKN1B, TET3, CD80, DRAM1, and ARID5B as associated with systemic lupus erythematosus in Asians. *Am J Hum Genet.* 2013;92(1):41-51.
64. Ellinghaus D, Jostins L, Spain SL, Cortes A, Bethune J, Han B, Park YR, Raychaudhuri S, Pouget JG, Hubenthal M, et al. Analysis of five chronic inflammatory diseases identifies 27 new associations and highlights disease-specific patterns at shared loci. *Nat Genet.* 2016;48(5):510-8.
65. International Genetics of Ankylosing Spondylitis C, Cortes A, Hadler J, Pointon JP, Robinson PC, Karaderi T, Leo P, Cremin K, Pryce K, Harris J, et al. Identification of multiple risk variants for ankylosing spondylitis through high-density genotyping of immune-related loci. *Nat Genet.* 2013;45(7):730-8.
66. Eyre S, Bowes J, Diogo D, Lee A, Barton A, Martin P, Zhernakova A, Stahl E, Viatte S, McAllister K, et al. High-density genetic mapping identifies new susceptibility loci for rheumatoid arthritis. *Nat Genet.* 2012;44(12):1336-40.
67. Trynka G, Hunt KA, Bockett NA, Romanos J, Mistry V, Szperl A, Bakker SF, Bardella MT, Bhaw-Rosun L, Castillejo G, et al. Dense genotyping identifies and localizes multiple common and rare variant association signals in celiac disease. *Nat Genet.* 2011;43(12):1193-201.
68. International Multiple Sclerosis Genetics C, Wellcome Trust Case Control C, Sawcer S, Hellenthal G, Pirinen M, Spencer CC, Patsopoulos NA, Moutsianas L, Dilthey A, Su Z, et al. Genetic risk and a primary role for cell-mediated immune mechanisms in multiple sclerosis. *Nature.* 2011;476(7359):214-9.
69. Waage J, Standl M, Curtin JA, Jessen LE, Thorsen J, Tian C, Schoettler N, andMe Research T, collaborators A, Flores C, et al. Genome-wide association and HLA fine-mapping studies identify risk loci and genetic pathways underlying allergic rhinitis. *Nat Genet.* 2018;50(8):1072-80.
70. Kim KW, Myers RA, Lee JH, Igartua C, Lee KE, Kim YH, Kim EJ, Yoon D, Lee JS, Hirota T, et al. Genome-wide association study of recalcitrant atopic dermatitis in Korean children. *J Allergy Clin Immunol.* 2015;136(3):678-84 e4.
71. Michailidou K, Hall P, Gonzalez-Neira A, Ghoussaini M, Dennis J, Milne RL, Schmidt MK, Chang-Claude J, Bojesen SE, Bolla MK, et al. Large-scale genotyping identifies 41 new loci associated with breast cancer risk. *Nat Genet.* 2013;45(4):353-61, 61e1-2.
72. Peters U, Hutter CM, Hsu L, Schumacher FR, Conti DV, Carlson CS, Edlund CK, Haile RW, Gallinger S, Zanke BW, et al. Meta-analysis of new genome-wide association studies of colorectal cancer risk. *Hum Genet.* 2012;131(2):217-34.
73. Iyengar SK, Sedor JR, Freedman BI, Kao WH, Kretzler M, Keller BJ, Abboud HE, Adler SG, Best LG, Bowden DW, et al. Genome-Wide Association and Trans-ethnic Meta-Analysis for Advanced Diabetic Kidney Disease: Family Investigation of Nephropathy and Diabetes (FIND). *PLoS Genet.* 2015;11(8):e1005352.

74. Barrett JC, Clayton DG, Concannon P, Akolkar B, Cooper JD, Erlich HA, Julier C, Morahan G, Nerup J, Nierras C, et al. Genome-wide association study and meta-analysis find that over 40 loci affect risk of type 1 diabetes. *Nat Genet.* 2009;41(6):703-7.
75. Hakonarson H, Grant SF, Bradfield JP, Marchand L, Kim CE, Glessner JT, Grabs R, Casalunovo T, Taback SP, Frackelton EC, et al. A genome-wide association study identifies KIAA0350 as a type 1 diabetes gene. *Nature.* 2007;448(7153):591-4.
76. Bentham J, Morris DL, Cunninghame Graham DS, Pinder CL, Tombleson P, Behrens TW, Martin J, Fairfax BP, Knight JC, Chen L, et al. Genetic association analyses implicate aberrant regulation of innate and adaptive immunity genes in the pathogenesis of systemic lupus erythematosus. *Nat Genet.* 2015;47(12):1457-64.
77. Cordell HJ, Han Y, Mells GF, Li Y, Hirschfield GM, Greene CS, Xie G, Juran BD, Zhu D, Qian DC, et al. International genome-wide meta-analysis identifies new primary biliary cirrhosis risk loci and targetable pathogenic pathways. *Nat Commun.* 2015;6(8019.
78. Mells GF, Floyd JA, Morley KI, Cordell HJ, Franklin CS, Shin SY, Heneghan MA, Neuberger JM, Donaldson PT, Day DB, et al. Genome-wide association study identifies 12 new susceptibility loci for primary biliary cirrhosis. *Nat Genet.* 2011;43(4):329-32.
79. Liu H, Irwanto A, Fu X, Yu G, Yu Y, Sun Y, Wang C, Wang Z, Okada Y, Low H, et al. Discovery of six new susceptibility loci and analysis of pleiotropic effects in leprosy. *Nat Genet.* 2015;47(3):267-71.
80. Stuart PE, Nair RP, Tsoi LC, Tejasvi T, Das S, Kang HM, Ellinghaus E, Chandran V, Callis-Duffin K, Ike R, et al. Genome-wide Association Analysis of Psoriatic Arthritis and Cutaneous Psoriasis Reveals Differences in Their Genetic Architecture. *Am J Hum Genet.* 2015;97(6):816-36.
81. Shah AA, Haynes C, Craig DM, Sebek J, Grass E, Abramson K, Hauser E, Gregory SG, Kraus WE, Smith PK, et al. Genetic variants associated with vein graft stenosis after coronary artery bypass grafting. *Heart Surg Forum.* 2015;18(1):E1-5.
82. Williams BB, Tebbutt NC, Buchert M, Putoczki TL, Doggett K, Bao S, Johnstone CN, Masson F, Hollande F, Burgess AW, et al. Glycoprotein A33 deficiency: a new mouse model of impaired intestinal epithelial barrier function and inflammatory disease. *Dis Model Mech.* 2015;8(8):805-15.
83. Alfredsson J, Puthalakath H, Martin H, Strasser A, and Nilsson G. Proapoptotic Bcl-2 family member Bim is involved in the control of mast cell survival and is induced together with Bcl-XL upon IgE-receptor activation. *Cell Death Differ.* 2005;12(2):136-44.
84. Bouillet P, Metcalf D, Huang DC, Tarlinton DM, Kay TW, Kontgen F, Adams JM, and Strasser A. Proapoptotic Bcl-2 relative Bim required for certain apoptotic responses, leukocyte homeostasis, and to preclude autoimmunity. *Science.* 1999;286(5445):1735-8.
85. Bouillet P, Purton JF, Godfrey DI, Zhang LC, Coultas L, Puthalakath H, Pellegrini M, Cory S, Adams JM, and Strasser A. BH3-only Bcl-2 family member Bim is required for apoptosis of autoreactive thymocytes. *Nature.* 2002;415(6874):922-6.
86. Enders A, Bouillet P, Puthalakath H, Xu Y, Tarlinton DM, and Strasser A. Loss of the pro-apoptotic BH3-only Bcl-2 family member Bim inhibits BCR stimulation-induced apoptosis and deletion of autoreactive B cells. *J Exp Med.* 2003;198(7):1119-26.

87. Kotzin JJ, Spencer SP, McCright SJ, Kumar DB, Collet MA, Mowel WK, Elliott EN, Uyar A, Makiya MA, Dunagin MC, et al. The long non-coding RNA Morrbid regulates Bim and short-lived myeloid cell lifespan. *Nature.* 2016;537(7619):239-43.
88. Dent AL, Shaffer AL, Yu X, Allman D, and Staudt LM. Control of inflammation, cytokine expression, and germinal center formation by BCL-6. *Science.* 1997;276(5312):589-92.
89. Corren J, Parnes JR, Wang L, Mo M, Roseti SL, Griffiths JM, and van der Merwe R. Tezepelumab in Adults with Uncontrolled Asthma. *N Engl J Med.* 2017;377(10):936-46.
90. Parulekar AD, Diamant Z, and Hanania NA. Role of T2 inflammation biomarkers in severe asthma. *Curr Opin Pulm Med.* 2016;22(1):59-68.
91. Ochiai K, Kagami M, Matsumura R, and Tomioka H. IL-5 but not interferon-gamma (IFN-gamma) inhibits eosinophil apoptosis by up-regulation of bcl-2 expression. *Clin Exp Immunol.* 1997;107(1):198-204.
92. Sitkauskiene B, Johansson AK, Sergejeva S, Lundin S, Sjostrand M, and Lotvall J. Regulation of bone marrow and airway CD34+ eosinophils by interleukin-5. *Am J Respir Cell Mol Biol.* 2004;30(3):367-78.
93. Ortega HG, Liu MC, Pavord ID, Brusselle GG, FitzGerald JM, Chetta A, Humbert M, Katz LE, Keene ON, Yancey SW, et al. Mepolizumab treatment in patients with severe eosinophilic asthma. *N Engl J Med.* 2014;371(13):1198-207.
94. Ochiai K, Katoh Y, Ikura T, Hoshikawa Y, Noda T, Karasuyama H, Tashiro S, Muto A, and Igarashi K. Plasmacytic transcription factor Blimp-1 is repressed by Bach2 in B cells. *J Biol Chem.* 2006;281(50):38226-34.
95. Richer MJ, Lang ML, and Butler NS. T Cell Fates Zipped Up: How the Bach2 Basic Leucine Zipper Transcriptional Repressor Directs T Cell Differentiation and Function. *J Immunol.* 2016;197(4):1009-15.
96. Vahedi G, Kanno Y, Furumoto Y, Jiang K, Parker SC, Erdos MR, Davis SR, Roychoudhuri R, Restifo NP, Gadina M, et al. Super-enhancers delineate disease-associated regulatory nodes in T cells. *Nature.* 2015;520(7548):558-62.
97. Kim EH, Gasper DJ, Lee SH, Plisch EH, Svaren J, and Suresh M. Bach2 regulates homeostasis of Foxp3+ regulatory T cells and protects against fatal lung disease in mice. *J Immunol.* 2014;192(3):985-95.
98. Pai SY, Truitt ML, and Ho IC. GATA-3 deficiency abrogates the development and maintenance of T helper type 2 cells. *Proc Natl Acad Sci U S A.* 2004;101(7):1993-8.
99. Ting CN, Olson MC, Barton KP, and Leiden JM. Transcription factor GATA-3 is required for development of the T-cell lineage. *Nature.* 1996;384(6608):474-8.
100. Zheng W, and Flavell RA. The transcription factor GATA-3 is necessary and sufficient for Th2 cytokine gene expression in CD4 T cells. *Cell.* 1997;89(4):587-96.
101. Zhu J, Min B, Hu-Li J, Watson CJ, Grinberg A, Wang Q, Killeen N, Urban JF, Jr., Guo L, and Paul WE. Conditional deletion of Gata3 shows its essential function in T(H)1-T(H)2 responses. *Nat Immunol.* 2004;5(11):1157-65.

102. Ano S, Morishima Y, Ishii Y, Yoh K, Yageta Y, Ohtsuka S, Matsuyama M, Kawaguchi M, Takahashi S, and Hizawa N. Transcription factors GATA-3 and RORgammat are important for determining the phenotype of allergic airway inflammation in a murine model of asthma. *J Immunol.* 2013;190(3):1056-65.
103. McKenzie AN. Type-2 innate lymphoid cells in asthma and allergy. *Ann Am Thorac Soc.* 2014;11 Suppl 5(S263-70.
104. Kim PJ, Pai SY, Brigl M, Besra GS, Gumperz J, and Ho IC. GATA-3 regulates the development and function of invariant NKT cells. *J Immunol.* 2006;177(10):6650-9.
105. Norata GD, Pirillo A, Callegari E, Hamsten A, Catapano AL, and Eriksson P. Gene expression and intracellular pathways involved in endothelial dysfunction induced by VLDL and oxidised VLDL. *Cardiovasc Res.* 2003;59(1):169-80.
106. Marine JC, Topham DJ, McKay C, Wang D, Parganas E, Stravopodis D, Yoshimura A, and Ihle JN. SOCS1 deficiency causes a lymphocyte-dependent perinatal lethality. *Cell.* 1999;98(5):609-16.
107. Takahashi R, Nishimoto S, Muto G, Sekiya T, Tamiya T, Kimura A, Morita R, Asakawa M, Chinen T, and Yoshimura A. SOCS1 is essential for regulatory T cell functions by preventing loss of Foxp3 expression as well as IFN-{gamma} and IL-17A production. *J Exp Med.* 2011;208(10):2055-67.
108. Ahmed CM, Larkin J, 3rd, and Johnson HM. SOCS1 Mimetics and Antagonists: A Complementary Approach to Positive and Negative Regulation of Immune Function. *Front Immunol.* 2015;6(183.
109. Li J, Jorgensen SF, Maggadottir SM, Bakay M, Warnatz K, Glessner J, Pandey R, Salzer U, Schmidt RE, Perez E, et al. Association of CLEC16A with human common variable immunodeficiency disorder and role in murine B cells. *Nat Commun.* 2015;6(6804.
